# Androgen aggravates aortic aneurysms via suppressing PD-1 in mice

**DOI:** 10.1101/2023.01.22.525073

**Authors:** Xufang Mu, Shu Liu, Zhuoran Wang, Kai Jiang, Tim McClintock, Arnold J. Stromberg, Alejandro V. Tezanos, Eugene S Lee, John A. Curci, Ming C Gong, Zhenheng Guo

## Abstract

Androgen has long been recognized for its pivotal role in the sexual dimorphism of cardiovascular diseases, including aortic aneurysms, a devastating vascular disease with a higher prevalence and mortality rate in men than women. However, the molecular mechanism by which androgen mediates aortic aneurysms is largely unknown. Here, we report that male but not female mice develop aortic aneurysms in response to aldosterone and high salt (Aldo-salt). We demonstrate that both androgen and androgen receptors (AR) are crucial for the sexually dimorphic response to Aldo-salt. We identify T cells expressing programmed cell death protein 1 (PD-1), an immune checkpoint molecule important in immunity and cancer immunotherapy, as a key link between androgen and aortic aneurysms. We show that intraperitoneal injection of anti-PD-1 antibody reinstates Aldo-salt-induced aortic aneurysms in orchiectomized mice. Mechanistically, we demonstrate that AR binds to the PD-1 promoter to suppress its expression in the spleen. Hence, our study reveals an important but unexplored mechanism by which androgen contributes to aortic aneurysms by suppressing PD-1 expression in T cells. Our study also suggests that cancer patients predisposed to the risk factors of aortic aneurysms may be advised to screen for aortic aneurysms during immune checkpoint therapy.

**Graphical Abstract:** 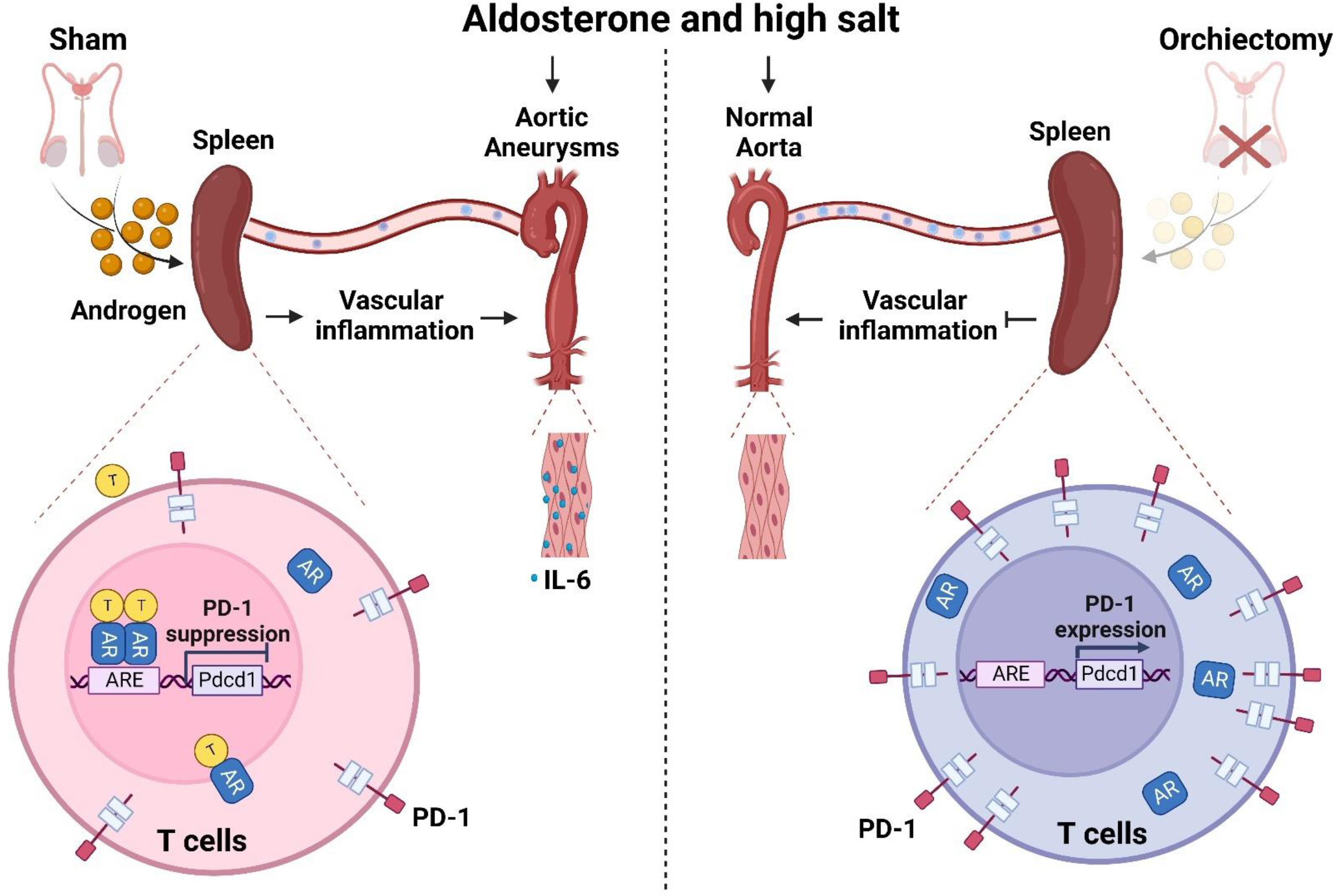

## Introduction

Aortic aneurysms are defined as a permanent localized dilation of the aorta with at least 3 mm or a 50% increase in diameter compared with a normal aortic diameter (1). Aortic aneurysms can be classified according to their location as thoracic aortic aneurysms (TAA) and abdominal aortic aneurysms (AAA) (2, 3). Aortic aneurysms are usually asymptomatic until they rupture, often lethal, resulting in over 85% mortality (4), affecting 4 to 8% of men and 0.5 to 1.5% of women over the age of 60, and accounting for nearly 2% of all deaths (1, 3, 5). Unfortunately, no medication except for open or endovascular surgeries has been approved to treat aortic aneurysms, largely attributed to the lack of mechanistic insight into this devastating vascular disease’s etiology and developmental pathogenesis.

Epidemiologic studies reveal aging, male sex, smoking, atherosclerosis, and hypertension as the risk factors for aortic aneurysms (5). In particular, the male sex is considered the most potent nonmodified risk factor for the sexual dimorphism of aortic aneurysms, with a 4:1 male-to-female ratio (5). In inflammatory aortic aneurysms, the male sex emerges as an even stronger risk factor, leading to male dominance in the development of AAA ranging from 6:1 to 30:1 (6). While the etiology of the sex difference in human aortic aneurysms remains to be elucidated, accumulated evidence from animal studies demonstrated that both sex chromosomes (X and Y chromosomes) and sex hormones (androgen and estrogen) contributed to the initiation and progression of aortic aneurysms (7). In particular, it is well documented that gonadal androgen deprivation via orchiectomy or deletion of androgen receptor (AR), but not gonadal estrogen deprivation via ovariectomy, protects angiotensin II (Ang II)-induced aortic aneurysms in hyperlipidemia mice (8-10) and elastase-induce aortic aneurysms in rats (11), indicating that androgen relative to estrogen seems to be mainly responsible for sexual dimorphism of aortic aneurysms in AAA animal models. However, how androgen mechanistically aggravates Ang II or elastin-induced aortic aneurysms at the molecular level remains largely unknown.

Accumulated clinical evidence manifests that elevated plasma concentration of aldosterone (Aldo), an essential component of the renin-angiotensin-aldosterone system, and excessive dietary sodium intake are associated with an increased risk for hypertension, stroke, coronary heart disease, heart failure, and renal disease (12). Consistent with these human studies, we developed a mouse model of aortic aneurysms in which we administered mineralocorticoid receptor (MR) agonists (i.e., Aldo to mimic elevated plasma concentration of Aldo in human patients) and high salt (to mimic excessive dietary sodium intake in human population) to 10-month-old male C57BL/6J mice (to imitate AAA in older men) to induce AAA, TAA, and aneurysmal rupture (13, 14). We demonstrated that Aldo and high salt (Aldo-salt)-induced aortic aneurysms are dependent on MR and aging but independent of Ang II (13, 14). However, whether Aldo-salt-induced aortic aneurysms have a sexual dimorphism has not been investigated. As a result, the role of androgen in Aldo-salt-induced aortic aneurysms is also completely unknown.

Here, we report that Aldo-salt-induced aortic aneurysms mostly occur in male but not female mice. We demonstrate that androgen and AR are crucial for this sexual dimorphism, which IL-6 partially mediates. Moreover, we identify splenic T cells expressing programmed cell death protein 1 (PD-1), an immune checkpoint molecule important in immunity and cancer immunotherapy (15), as a new androgen target gene critical for Aldo-salt-induced aortic aneurysms. Mechanistically, we demonstrate that AR binds to the PD-1 promoter to suppress its expression in the spleen. Our findings suggest that cancer patients with immune checkpoint therapy have a risk of developing aortic aneurysms.

## Results

### Sexual dimorphism in Aldo-salt-induced aortic aneurysms

To investigate whether there is sexual dimorphism in Aldo-salt-induced aortic aneurysms, 10-month-old male and female C57BL/6 mice were administered Aldo (200 µg/kg/day via osmotic pumps) and high salt (0.9% NaCl plus 0.2% KCl in drinking water) for four weeks to induce aortic aneurysms (13, 14). The effect of Aldo-salt on the intraluminal diameter of the abdominal aorta was quantified *in vivo* by a high-resolution ultrasound imaging system one week before (time 0; to set up a baseline) and weekly after Aldo-salt administration (to monitor the progression of aortic dilation). As shown in Figure 1A, Aldo-salt induced suprarenal aortic dilation in both male and female mice in a time-dependent manner. However, the suprarenal aortic dilation induced by Aldo-salt was much larger in male than in female mice (male *vs*. female, *p* < 0.0001). Based on the ultrasound data, we calculated the growth rate of the abdominal aortic diameters, an important clinical index for appraisal of aortic aneurysm progression and rupture in human patients (4). As expected, the suprarenal aortic growth rate was significantly accelerated in male than female mice (Figure 1B).

**Figure 1.**
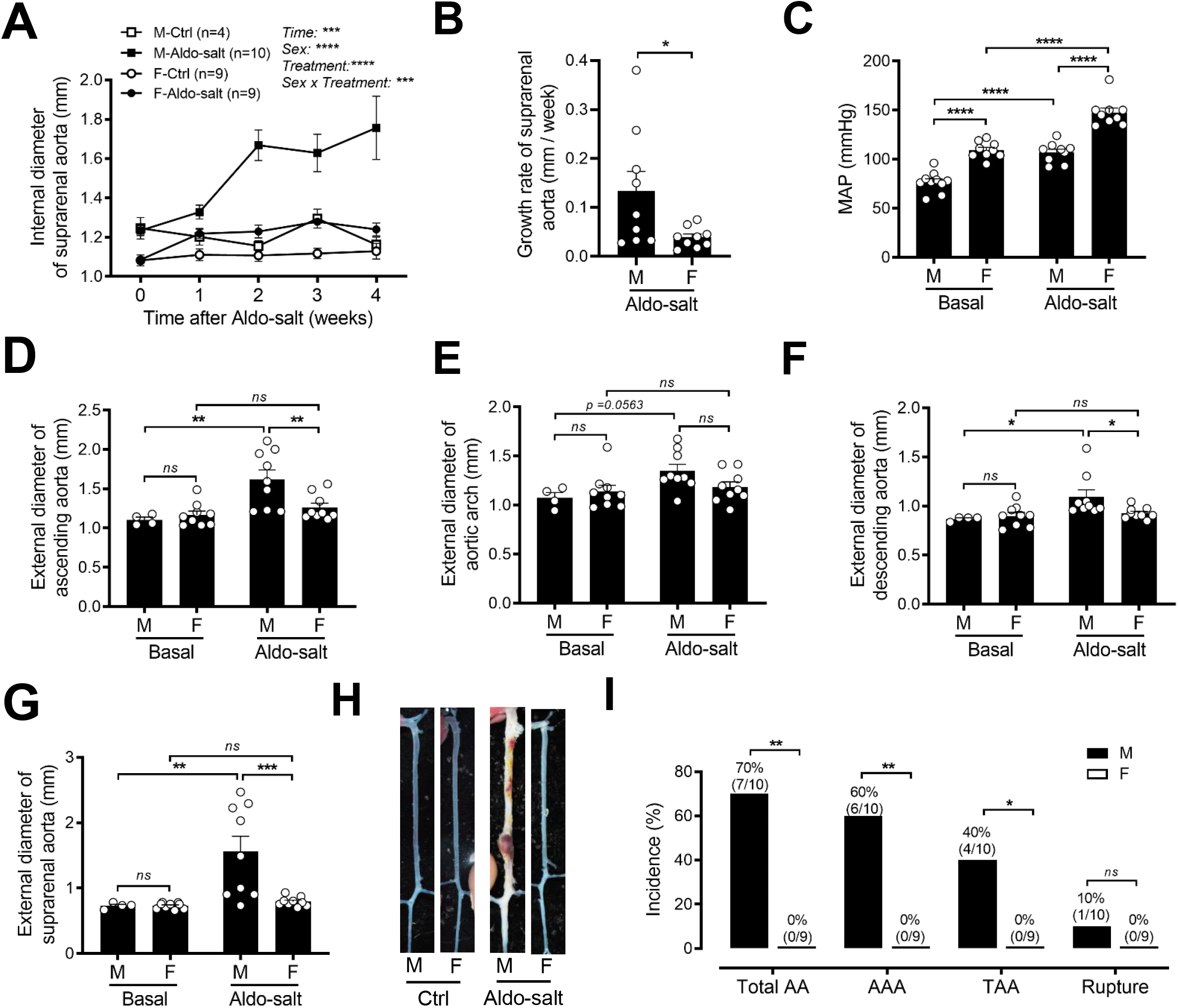
Sexual dimorphism in Aldo-salt-induced aortic aneurysms. (**A** and **B**) *In vivo* quantification of the maximal intraluminal diameters (A) and growth rate (B) of the suprarenal aortas weekly by ultrasound in 10-month-old male (M) and female (F) C57BL/6J mice one week before (0) and 1−4 weeks after Aldo-salt administration. (**C**) Mean arterial pressure (MAP) was measured by tail cuff one week before (basal) and three weeks after Aldo-salt administration. (**D−G**) *Ex vivo* measurement of the maximal external diameters of the ascending aorta (D), aortic arch (E), descending aorta (F), and suprarenal aorta (G) by microscopy in the isolated aortas four weeks after Aldo-salt administration. (**H**) Representative photographs of the aortas with or without aortic aneurysms in male and female mice four weeks after Aldo-salt administration. (**I**) The incidence of total aortic aneurysms (AA), abdominal aortic aneurysms (AAA), thoracic aortic aneurysms (TAA), and aortic aneurysm rupture. The percentages of the incidences (number of mice with total AA, AAA, TAA, and rupture / total number of mice) were indicated on the top of each bar. The data were expressed as mean ± standard error of the mean (SEM) and analyzed by thee-way ANOVA analysis (A), two-tailed unpaired *t*-test (B), two-way ANOVA with multiple comparison test (C−G), and two-sided Chi-square test (I). *\*, p < 0*.*05; **, p < 0*.*01; ***, p < 0*.*001; ****, p < 0*.*0001; ns, not significant*.

To determine whether the sexual dimorphism of Aldo-salt-induced progression of aortic dilation was related to the well-recognized sexual dimorphism of blood pressure (16), we measured blood pressure in the male and female mice by a non-invasive tail-cuff system one week before (to set up a baseline) and three weeks after Aldo-salt administration. Both male and female mice responded to Aldo-salt to develop hypertension (i.e., elevated mean arterial pressure (MAP)), but female mice surprisingly exhibited a higher blood pressure than male mice before and after Aldo-salt administration (Figure 1C), indicating that the larger increase in the suprarenal aortic dilation in response to Aldo-salt in male mice relative to female mice is not attributed to more severe hypertension in male mice than in female mice.

Four weeks after Aldo-salt administration, the aortas were isolated from the male and female mice and subjected to morphometric analysis to measure the maximal external diameters of the ascending aorta, aortic arch, descending aorta, and suprarenal aorta (Supplemental Figure 1). In agreement with the ultrasound results, there was a significant increase or an increasing trend in the external diameters of the ascending aorta, aortic arch, descending aorta, and suprarenal aorta in response to Aldo-salt in male mice but not in female mice (Figures 1D−1G). Isolated aortas were also subjected to pathological analysis to calculate the incidence of the total aortic aneurysms (total AA; total AA = AAA + TAA + aortic aneurysm rupture), AAA, TAA, and aortic aneurysm rupture based on either 50% increase in aortic internal or external diameters or evident aortopathy (Figure 1H) (13, 14, 17). Consistent with our previous reports (13, 14), of 10 male mice, 7 developed aortic aneurysms (70%), including 6 AAA (60%), 4 TAA (40%), and 1 aortic aneurysm rupture (10%) (Figure 1I). In contrast, of 9 female mice, none developed aortic aneurysms and aortic aneurysm ruptures (Figure 1I). Thus, Aldo-salt-induced aortic aneurysms exhibit a strong sex dimorphism with dominance in male mice.

### Gonadal androgen deprivation protects mice from Aldo-salt-induced aortic aneurysm

To investigate whether androgen, a group of male sex hormones, is involved in the sexual dimorphism in Aldo-salt-induced aortic aneurysms, 10-month-old male C57BL/6 mice were subjected to orchiectomy (to deprive gonadal androgen) or sham operation (to serve as a surgical control). After two weeks of orchiectomy, the mice were administered Aldo-salt for four weeks (to induce aortic aneurysms). To determine if orchiectomy succeeds, we measured seminal vesicle weight (SVW) in orchiectomized and sham-operated mice four weeks after Aldo-salt administration. Consistent with the previous report (18), both SVW and the SVW/body weight (BW) ratio (to rule out the effect of BW) were markedly suppressed in orchiectomized mice compared to sham-operated mice (Figure 2A and Supplemental Figures 2A and 2B). In addition, we measured serum testosterone levels with an ELISA kit in some of these mice four weeks after Aldo-salt administration. Consistent with the SVW data, serum testosterone levels were significantly lower in orchiectomized mice than in sham-operated mice (Supplemental Figure 2C).

**Figure 2.**
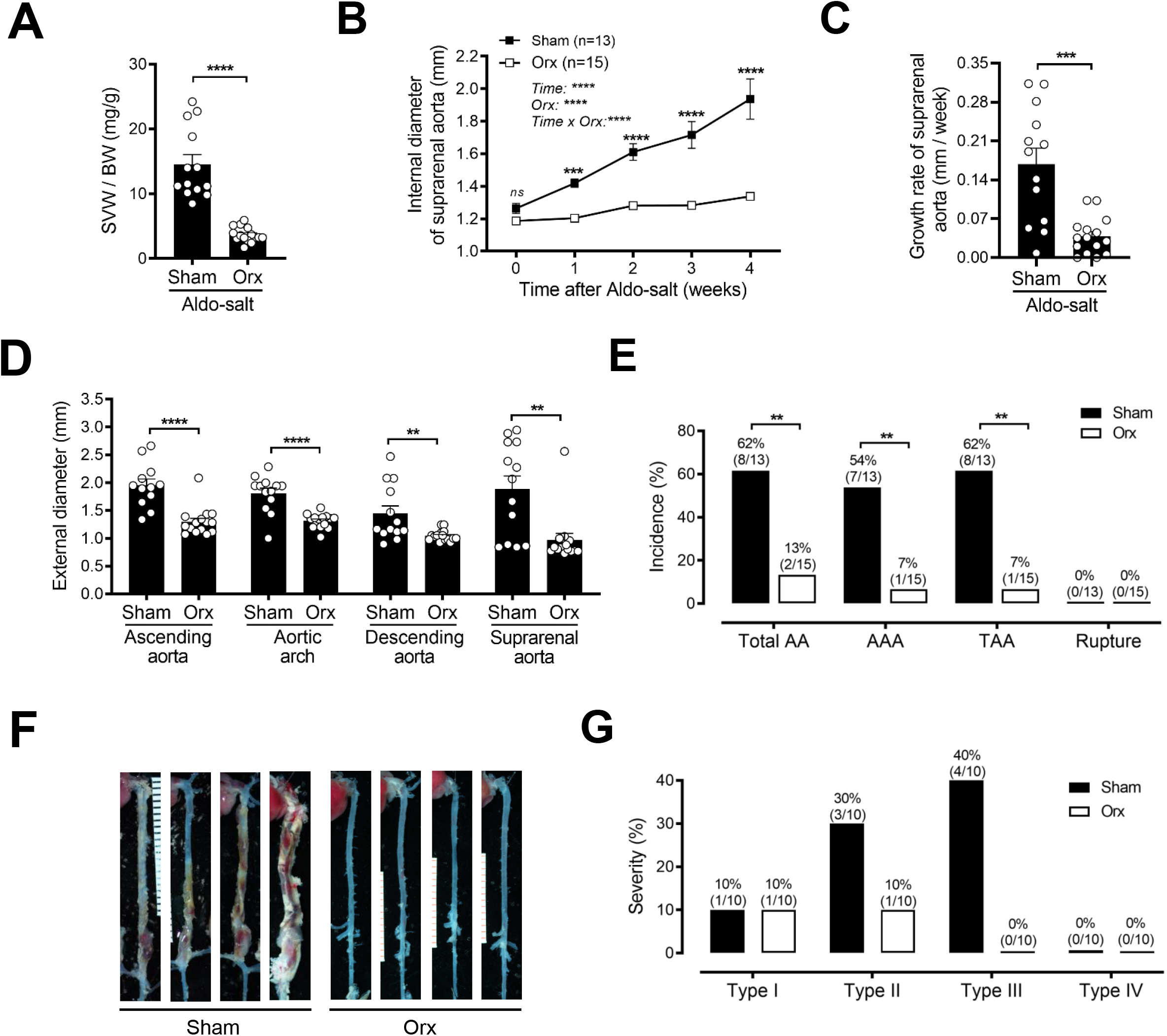
Orchiectomy protects mice from Aldo-salt-induced aortic dilation, progression, and aneurysm formation. (**A**) Seminal vesicle weight (SVW) to body weight (BW) ratio was determined in 10-month-old male C57BL/6J mice with orchiectomy (orx) or sham operation four weeks after Aldo-salt administration. (**B** and **C)** *In vivo* quantification of the maximal intraluminal diameters (B) and growth rate (C) of the suprarenal aortas weekly by ultrasound in mice with orchiectomy or sham operation one week before (0) and 1−4 weeks after Aldo-salt administration. (**D**) *Ex vivo* measurement of the maximal external diameters of the ascending aorta, aortic arch, descending aorta, and suprarenal aorta by microscopy in isolated aortas four weeks after Aldo-salt administration. (**E**) The incidence of total aortic aneurysms (AA), AAA, TAA, and aortic aneurysm rupture. The percentages of the incidences (number of mice with total AA, AAA, TAA, and rupture / total number of mice) were indicated on the top of each bar. (**F**) Representative photographs of the aortas with or without aortic aneurysms in orchiectomized or sham-operated mice four weeks after Aldo-salt administration. (**G**) The severity of aortic aneurysms was expressed as the percentages of Type I, II, III, and IV aortic aneurysms. The percentages of severity (number of mice with Type I, II, III, and IV aortic aneurysms / total number of mice with aortic aneurysms) were indicated on the top of each bar. The data were expressed as mean ± SEM and analyzed by two-tailed unpaired *t*-test (A, C, D), two-way ANOVA with multiple comparison test (B), and two-sided Chi-square test (E). *\*, p < 0*.*05; **, p < 0*.*01; ***, p < 0*.*001; ****, p < 0*.*0001; ns, not significant*.

To investigate whether gonadal androgen deprivation affects Aldo-salt-induced abdominal aortic aneurysms, we measured the internal and external diameters of the abdominal and thoracic aorta *in vivo* weekly by ultrasound and *ex vivo* by microscopy four weeks after Aldo-salt administration, respectively. Compared to the sham operation, orchiectomy completely abolished Aldo-salt-induced suprarenal aortic aneurysm formation and progression (Figures 2B and 2C). In line with these *in vivo* ultrasound results, orchiectomy also significantly suppressed the increase in the external diameters of the ascending aorta, aortic arch, descending aorta, and suprarenal aorta four weeks after Aldo-salt administration relative to the sham operation (Figure 2D), indicating the role of androgen in Aldo-salt-indued TAA and AAA. To further define the effect of gonadal androgen deprivation on aortic aneurysms, we compared the incidence of Aldo-salt-induced aortic aneurysms between orchiectomized and sham-operated mice. As shown in Figure 2E, of 13 sham-operated mice, 8 developed aortic aneurysms (62%), including 7 AAA (54%) and 8 TAA (62%). Conversely, of 15 orchiectomized mice, 2 developed aortic aneurysms (13%), including 1 AAA (7%) and 1 TAA (7%). No aortic aneurysm rupture was found in mice with orchiectomy and sham operation.

Resembling human aortic aneurysms and consistent with Ang II-induced aortic aneurysms (17), the severity of Aldo-salt-induced aortic aneurysms varied considerably (13, 14). As described in the Ang II AAA mouse model (17), Aldo-salt-induced aortic aneurysms can also be classified as Type I (the gross appearance of aortic dilation), Type II (at least a 2-time increase in external diameter of the aorta), Type III (a pronounced bulbous form of Type II with a thrombus), and Type IV (aortic aneurysm rupture) (Supplemental Figure 3). To define the effect of gonadal androgen deprivation on the severity of Aldo-salt-induced aortic aneurysms, we calculated the percentage of Type I, II, III, and IV aortic aneurysms from the total number of aortic aneurysms from orchiectomized and sham-operated mice. As shown in Figures 2F and 2G, in sham-operated mice, there were 10% Type I, 30% Type II, and 40% Type IV aortic aneurysms; in contrast, in orchiectomized mice, there were 10% Type I and 10% Type II aortic aneurysms. These data suggest that androgen may mainly affect Aldo-salt-induced aortic aneurysm progression.

We previously demonstrated that high salt is required for Aldo-salt-induced aortic aneurysms (13). Therefore, we wondered whether androgen augments Aldo-salt-induced sodium retention, thus promoting aortic aneurysms. We determined 24-h sodium retention in orchiectomized and sham-operated mice by subtracting their 24-h sodium excretion (via urine) from their sodium intake (via food and water intake) as described (19) one week before (basal) and three weeks after Aldo-salt administration. Interestingly, orchiectomy increased both 24-h sodium intake (mainly via high salt water intake but not food intake) and 24-urinary sodium excretion relative to the sham operation (Supplemental Figures 4A−4E). As a net balance, orchiectomy did not alter Aldo-salt-indued sodium retention (Supplemental figure 4F). Consistent with these results, we also did not find any significant difference in serum sodium levels between the orchiectomized and sham-operated mice four weeks after Aldo-salt administration (Supplemental Figures 4G). There was also no correlation of the serum sodium level with the internal diameter of the suprarenal aorta four weeks after Aldo-salt administration (Supplemental Figure 4H). Moreover, there were no significant differences in serum sodium levels between the orchiectomized and sham-operated mice, regardless of whether the mice developed aortic aneurysms (Supplemental Figure 4I). Hence, these results suggest that androgen unlikely exacerbates Aldo-salt-induced aortic aneurysms via sodium retention.

Hypertension is a risk factor for human aortic aneurysms (5). We, therefore, investigated whether androgen exacerbates Aldo-salt-induced hypertension, thus promoting aortic aneurysms. Blood pressures were measured by tail cuff in the orchiectomized and sham-operated mice one week before (basal) and three weeks after Aldo-salt administration. In line with its little effect on sodium retention, orchiectomy did not affect MAP before and after Aldo-salt administration (Supplemental Figure 4J). There was no correlation of the MAP with the internal diameter of the suprarenal aorta three weeks after Aldo-salt administration (Supplemental Figure 4K). There were also no significant MAP differences between orchiectomized and sham-operated mice, regardless of whether they developed aortic aneurysms (Supplemental Figure 4L). Similar effects of orchiectomy on blood pressure were also found with systolic and diastolic blood pressure (data not shown). Thus, these results suggest that androgen unlikely aggravates Aldo-salt-induced aortic aneurysms via increasing blood pressure.

### Restoration of androgen in orchiectomized mice reinstates Aldo-salt-induced aortic aneurysms

To further define the role of androgen in Aldo-salt-induced aortic aneurysms, 10-month-old male C57BL/6J mice were subjected to orchiectomy for two weeks (to deplete gonadal androgen) and then administered with Aldo-salt (to induce aortic aneurysms) with or without dihydrotestosterone (DHT) pellet implantation (10 mg, 60-day release) for four weeks (to restore androgen in orchiectomized mice) (9). We used DHT, but not testosterone, because DHT is more potent than testosterone and, more importantly, because DHT, rather than testosterone, cannot be converted to estrogens by aromatase (20), which makes the results more readily interpretable. To ensure implanted DHT pellets function in mice, we weighed SVW from orchiectomized mice with or without DHT pellet implantation four weeks after Aldo-salt administration. Consistent with the effect of gonadal androgen deprivation on SVW and SVW/BW in orchiectomized mice (Figure 2A and Supplemental Figures 2A and 2B), DHT pellet implantation effectively restored the SVW and SVW/BW ratio in orchiectomized mice to a level comparable with that in sham-operated mice (Figure 3A and Supplemental Figures 5).

**Figure 3.**
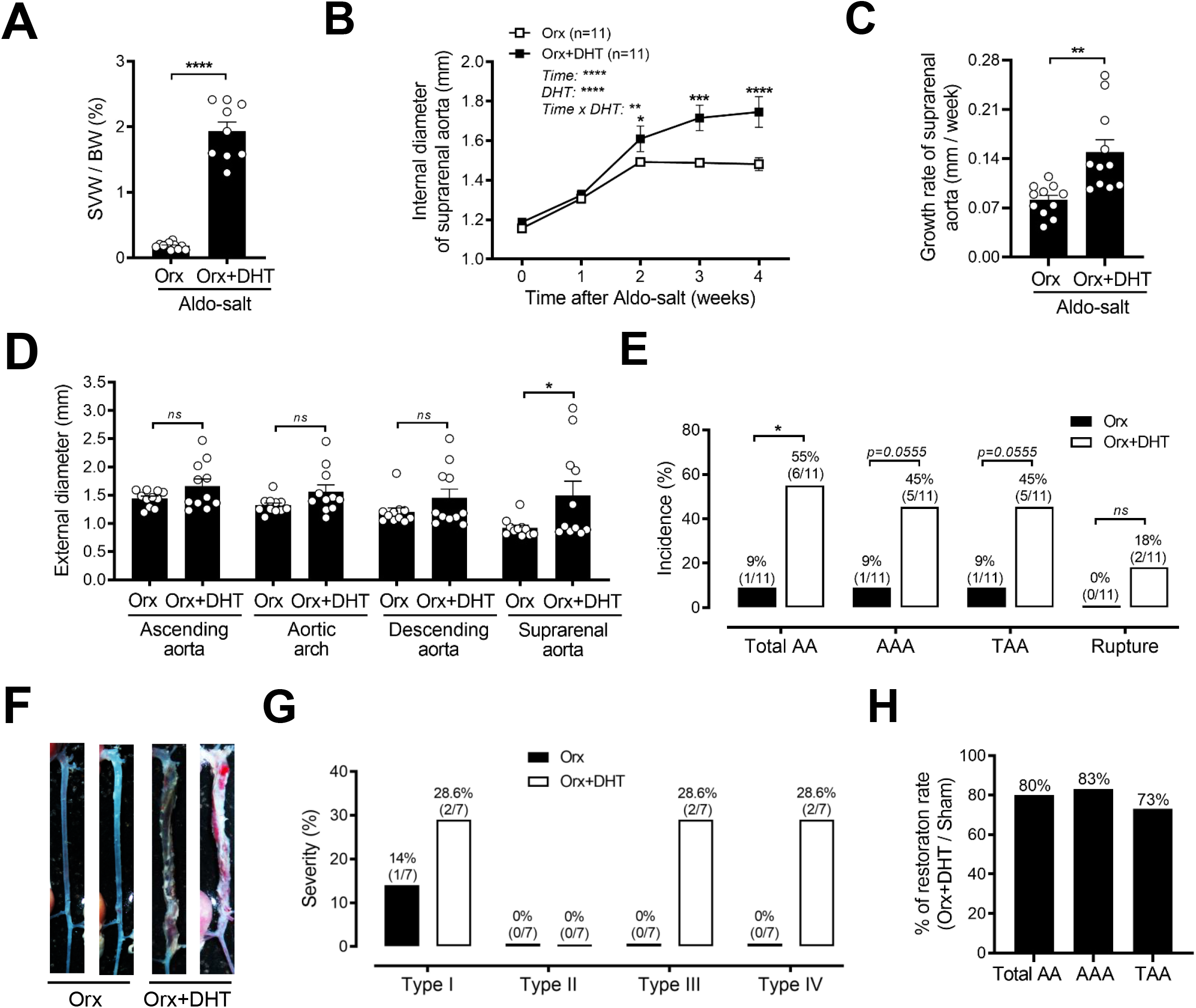
Exogenous dihydrotestosterone administration to orchiectomized male mice restores Aldo-salt-induced aortic aneurysms. (**A**) Seminal vesicle weight (SVW) to body weight (BW) ratio was determined in 10-month-old male C57BL/6J mice with orchiectomy (orx) 4 weeks after Aldo-salt with or without dihydrotestosterone (DHT) pellet implantation. (**B** and **C**) *In vivo* quantification of the maximal intraluminal diameters (B) and growth rate (C) of the suprarenal aortas weekly by ultrasound in orchiectomized mice one week before (0) and 1−4 weeks after Aldo-salt with or without DHT pellet implantation. (**D**) *Ex vivo* measurement of the maximal external diameters of the ascending aorta, aortic arch, descending aorta, and suprarenal aorta by microscopy in isolated aortas in orchiectomized mice four weeks after Aldo-salt with or without DHTpellet implantation. (**E**) The incidence of total aortic aneurysms (AA), AAA, TAA, and aortic aneurysm rupture. The percentages of the incidences (number of mice with total AA, AAA, TAA, and rupture / total number of mice) were indicated on the top of each bar. (**F**) Representative photographs of the aortas with or without aortic aneurysms in orchiectomized mice four weeks after Aldo-salt with or without DHT pellet implantation. The severity of aortic aneurysms was expressed as the percentages of Type I, II, III, and IV aortic aneurysms. The percentages of severity (number of mice with Type I, II, III, and IV aortic aneurysms / total number of mice with aortic aneurysms) were indicated on the top of each bar. (**H**) The percentage of the restoration rates (number of total AA, AAA, and TAA in orchiectomized mice with DHT / number of total AA, AAA, and TAA in sham-operated mice). The data were expressed as mean ± SEM and analyzed by two-tailed unpaired *t*-test (A, C, and D), two-way ANOVA with multiple comparisons test (B), and two-sided Chi-square test (E). *\*, p <0*.*05;* ; *\*\*, p < 0*.*01; ***, p < 0*.*001; ****, p < 0*.*0001; ns, not significant*.

The effect of DHT pellet implantation on Aldo-salt-induced aortic aneurysms was appraised weekly by *in vivo* ultrasound in orchiectomized mice before (basal) and after Aldo-salt administration. As shown in Figures 3B and 3C, restoration of androgen in orchiectomized mice partially but significantly restored Aldo-salt-induced suprarenal aortic dilation and progression. Interestingly, the significant increase in the internal diameters of the suprarenal aorta was only observed two weeks after Aldo-salt administration (Figure 3B), indicating that exogenous DHT administration to orchiectomized mice is mainly involved in the progression rather than the initiation of Aldo-salt-induced aortic aneurysms. Consistent with these data, there was a trend or significant increase toward the external diameters of the ascending aorta, aortic arch, descending aorta, and suprarenal aorta in orchiectomized mice with DHT implantation compared to orchiectomized mice without DHT implantation (Figures 3D).

Of 11 orchiectomies mice with DHT pellet implantation, 6 developed aortic aneurysms (55%), including 5 AAA (45%), 5 TAA (45%), and 2 aortic aneurysm rupture (18%). On the contrary, of 11 ovariectomized mice without DHT pellet implantation, only 1 developed with aortic aneurysms (9%), including AAA and TAA, and none developed aortic aneurysm rupture (Figure 3E). Of note, the aortopathies in orchiectomized mice with DHT pellet implantation also appeared more severe than that in sham-operated mice, as evidenced by 26.6% *vs*. 0% of mice developed Type IV aortic aneurysms in orchiectomized mice with DHT pellet implantation relative to sham-operated mice (Figures 3F and 3G *vs*. 2F and 2G). To quantify to what extent exogenous DHT administration restores Aldo-salt-induced aortic aneurysms, we calculated the percentage of aortic aneurysm restoration rates by normalizing the incidence of aortic aneurysms in orchiectomized mice with DHT pellets (Figure 3E) to that in sham-operated mice (Figure 2E). As a result, 80%, 83%, and 73% of aortic aneurysm restoration rates were obtained in orchiectomized mice with DHT pellet implantation for total AA, AAA, and TAA, respectively (Figure 3H). These results are consistent with the *in vivo* and *ex vivo* measurement of the aortic diameters and illustrate that restoration of androgen in orchiectomized mice reinstates Aldo-salt-induced aortic aneurysms.

To investigate whether Aldo-salt-induced sodium retention and hypertension are implicated in the exogenous DHT restoration of Aldo-salt-induced aortic aneurysms, we measured sodium retention, serum sodium level, and blood pressure in the orchiectomized mice with or without DHT pellet implantation. Surprisingly, exogenous DHT administration to orchiectomized mice significantly suppressed both 24-h sodium intake and 24-h urinary sodium excretion three weeks after Aldo-salt administration (Supplemental Figures 6A−6E). As a net result, exogenous DHT administration to orchiectomized mice did not affect Aldo-salt-induced sodium retention, serum sodium levels, and hypertension (Supplemental Figures 6F, 6G, and 6J). There was no correlation between the internal diameter of the suprarenal aortas and serum sodium or MAP levels in orchiectomized mice with and without DHT pellet implantation (Supplemental Figures 6H and 6K). There was also no significant difference in serum sodium or MAP levels in mice with or without DHT pellet implantation, regardless of whether they developed aortic aneurysms (Supplemental Figures 6I and 6L).

### Downregulation of androgen receptors ameliorates Aldo-salt-induced aortic aneurysms

To explore targeting androgen as a potential anti-aortic aneurysm therapy, 10-month-old male C57BL/6J mice were administered with Aldo-salt (to induce aortic aneurysms) with or without ASC-J9 (50 mg/kg, intraperitoneal injection, once a day) for four weeks (to downregulate AR) (10). ASC-J9 is a recently-developed AR-degradation enhancer and has been shown to selectively degrade AR without affecting other nuclear receptors in cultured cells and mice (21). The efficacy of ASC-J9 on AR protein expression was examined by immunocytochemistry (IHC) in the suprarenal aorta in the mice with or without ASC-J9 treatment four weeks after Aldo-salt administration. In the control mice without ASC-J9 treatment, AR immunostaining was found all over the crosssection of the suprarenal aorta (Figure 4A-a), which is consistent with its ubiquitous expression, including aortic smooth muscle cells (20). In contrast, in the mice treated with ASC-J9, little AR immunostaining was observed in the crosssection of the suprarenal aorta (Figure 4A-b), indicating that intraperitoneal injection of ASC-J9 effectively downregulates AR protein expression in the aorta.

**Figure 4.**
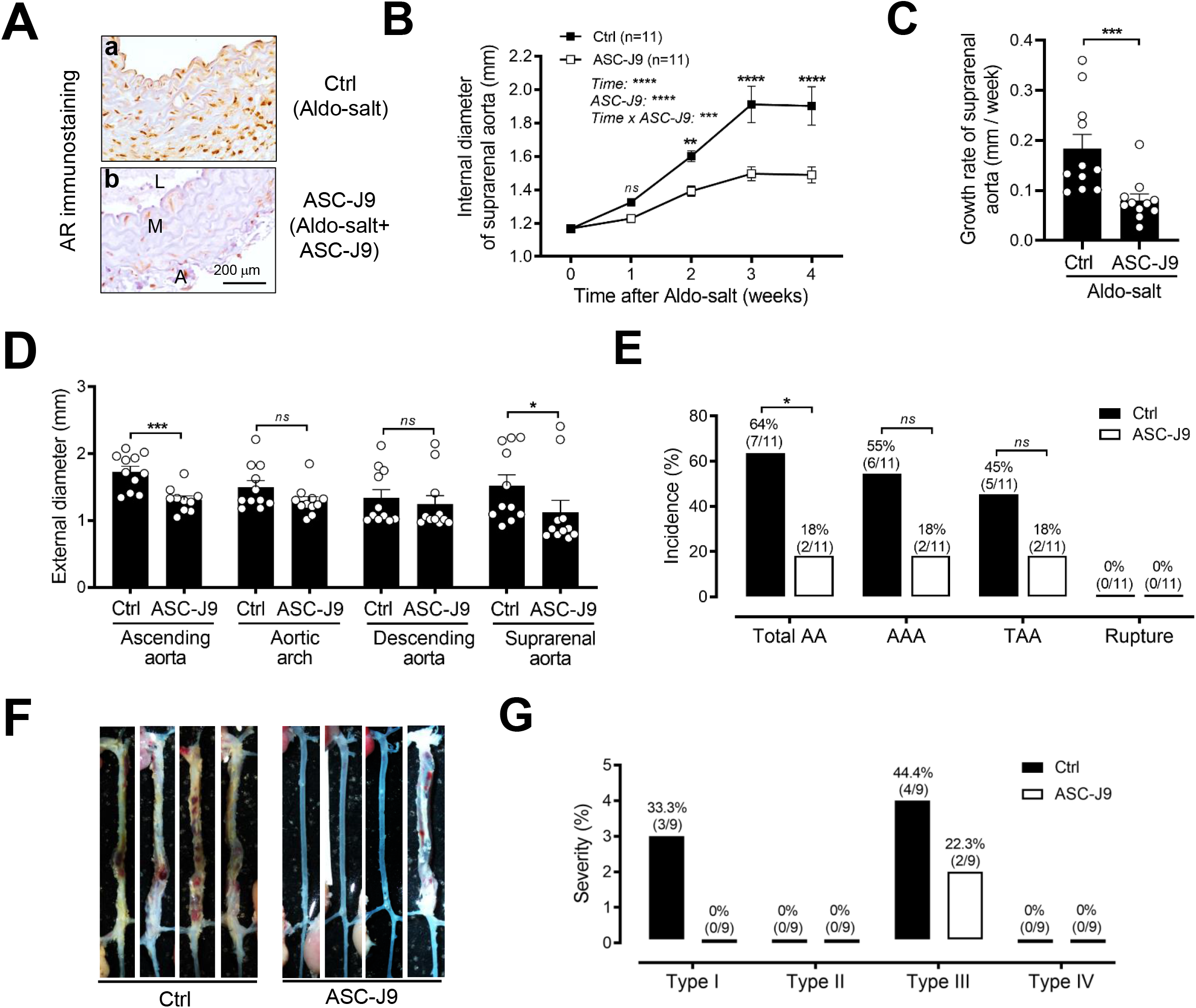
Downregulation of androgen receptors by ASC-J9 in mice inhibits Aldo-salt-induced aortic aneurysm. (**A**) Representative immunostaining of androgen receptor (AR) in the abdominal aorta from six 10-month-old male C57BL/6J mice four weeks after Aldo-salt with or without ASC-J9 administration (3 mice/group). L, lumen. M, media. A, adventitia. (**B** and **C**) *In vivo* quantification of the maximal intraluminal diameters (B) and growth rate (C) of the suprarenal aortas by ultrasound in mice one week before (0) and weekly after Aldo-salt with or without ASC-J9 administration. (**D**) *Ex vivo* measurement of the external diameters of the ascending aorta, aortic arch, and descending aorta by microscopy four weeks after Aldo-salt with or without ASC-J9 administration. (**E**) The incidence of total aortic aneurysms (AA), AAA, TAA, and aortic aneurysm rupture. The percentages of the incidences (number of mice with total AA, AAA, TAA, and rupture / total number of mice) were indicated on the top of each bar. (**F**) Representative photographs of the aortas with or without aortic aneurysm in mice four weeks after Aldo-salt with or without ASC-J9 administration. (**G**) The severity of aortic aneurysms was expressed as the percentages of Type I, II, III, and IV aortic aneurysms. The percentages of severity (number of mice with Type I, II, III, and IV aortic aneurysms / total number of mice with aortic aneurysms) were indicated on the top of each bar. The data were expressed as mean ± SEM and analyzed by two-way ANOVA with multiple comparisons test (B), two-tailed unpaired *t*-test (C and D), and two-sided Chi-square test (E). *\*, p<0*.*05; **, p<0*.*01; ***, p<0*.*001; ****, p < 0*.*0001; ns, not significant*.

In agreement with the crucial role of androgen in Aldo-salt-induced aortic aneurysms (Figure 2), downregulation of AR by ASC-J9 significantly suppressed Aldo-salt-induced increases in the internal diameters of the suprarenal aorta (Figure 4B), the growth rate of the suprarenal aorta (Figure 4C), the external diameters of the ascending and suprarenal aorta (Figure 4D), and the incidence of total aortic aneurysms (Figure 4E). Nevertheless, it was noted that, of 11 mice treated with ASC-J9, 2 still responded to Aldo-salt to develop aortic aneurysms (Figures 4E and 4G). Given that individual variation in pharmacokinetics is well recognized (22), we speculated that these two mice might resist ASC-J9-induced AR degradation and thereby develop aortic aneurysms. To explore this possibility, we examined AR protein expression in the suprarenal aorta by IHC from these two mice and two other mice treated with ASC-J9 but did not develop AAA. As speculated, abundant AR protein expression was readily detectable in the suprarenal aorta of these two mice but not in the other two mice treated with ASC-J9 but did not develop AAA (Supplemental Figures 7a−7d *vs*. 7e−7h).

We also determined the effect of ASC-J9 on Aldo-salt-induced sodium retention, serum sodium levels, and hypertension in mice treated with or without ASC-J9. Intriguingly, ASC-J9 suppressed Aldo-salt-induced 24-h sodium intake and urinary sodium excretion (Supplemental Figures 8A−8E), which is the opposite of the effect of orchiectomy (Supplemental Figure 4). However, ASC-J9 did not significantly affect Aldo-salt-induced sodium retention, serum sodium levels, and hypertension (Supplemental Figures 8F, 8G, and 8J). There was no correlation between the internal diameters and serum sodium or MAP levels in mice treated with and without ASC-J9 (Supplemental Figures 8H and 8K). There was also no difference in serum sodium or MAP levels between mice treated with and without ASC-J9, regardless they developed aortic aneurysms (Supplemental Figures 8I and 8L).

### IL-6 is implicated in Aldo-salt-induced and androgen-mediated aortic aneurysms

To investigate the molecular mechanism by which androgen mediates Aldo-salt-induced aortic aneurysms, we determined mRNA expressions in the aorta from 10-month-old male C57BL/6 mice administration for ten days after orchiectomy and sham operation. We isolated the aortas ten days rather than four weeks after Aldo-salt administration because we aimed to identify androgen-targeting genes that result in rather than result from Aldo-salt-induced aortic aneurysms.

Based on the literature (3, 10, 13, 14, 23, 24), we focused on a list of genes that are implicated in aortic aneurysms, including AR (*Ar*), MR (*Nr3c2*), serum and glucocorticoid-regulated Kinase 1 (*Sgk1*), sodium channel epithelial 1 subunit alpha (*Scnn1a*; also known as ENaCα), *Scnn1b* (also known as EnaCβ), *Scnn1g* (also known as ENaCγ), aryl hydrocarbon receptor nuclear translocator-like (*Arntl*; also known as Bmal1), TGF beta 2 (*Tgfb2*), matrix metallopeptidase 2 (*Mmp2*), IL 1 beta (*Il1b*; also known as IL-1β), *Il6* (also known as IL-6), IL-6 receptor alpha (*Il6ra*; also known as IL-6Rα), IL-6 signal transducer (*Il6st*; also known as IL-6Rβ or gp130), C-C motif chemokine ligand 2 (*Ccl2*; also known as MCP-1), *Ccl4*, and tumor necrosis factor (*Tnf*; also known as TNF-α).

Of 16 mRNAs examined, 12 mRNAs, *Ar, Nr3C2, Sgk1, Scnn1a, Scnn1b, Scnn1g, Arntl, Il1b, Il6, Il6ra, Ccl2*, and *Ccl4*, responded to Aldo-salt: 9 of them, *Ar, Nr3C2, Sgk1, Scnn1a, Scnn1b, Scnn1g, Il1b, Il6ra*, and *Ccl4*, were downregulated, whereas 3 of them, *Arntl, Il6*, and *Ccl2*, were upregulated (Supplemental Figure 9). Interestingly, 5 mRNAs, *Nr3C2, Scnn1a, Scnn1b, Il6*, and *Ccl2*, also responded to orchiectomy after Aldo-salt administration: 3 of them, *Nr3C2, Scnn1a, Scnn1b*, were upregulated, whereas 2 of them, *Il6*, and *Ccl2*, were downregulated (Supplemental Figures 9B, 9D, 9E, 9K, and 9N). In particular, *Il6* was most dramatically upregulated by Aldo-salt (i.e., up to 58-fold), which was completely abolished by orchiectomy (Supplemental Figure 9K).

Since IL-6 is implicated in human aortic aneurysms (24), we focused on IL-6 and further investigated whether IL-6 protein expression responds to Aldo-salt and/or androgen in the aorta. The suprarenal aortas were isolated from 10-month-old male C57BL/6 orchiectomized and sham-operated mice ten days after Aldo-salt administration and then subjected to IHC analysis. IL-6 protein was barely detectable in the aorta in sham-operated mice without Aldo-salt administration (Supplemental Figure 10a). Interestingly, IL-6 protein was markedly upregulated in the aorta in sham-operated mice with Aldo-salt administration (Supplemental Figure 10b *vs*. 10a), which is associated with aortic aneurysm formation (Figure 2). Surprisingly, an IL-6 protein basal level increase was found in the aorta in orchiectomized mice without Aldo-salt administration (Supplemental Figure 10c *vs*. 10a), indicating that androgen suppresses basal IL-6 protein expression. Importantly, in line with their effect on *Il6* mRNA expression, orchiectomy also prevented Aldo-salt-induced IL-6 protein upregulation in the aorta (Supplemental Figure 10d), which correlates with its protective effect on Aldo-salt-induced aortic aneurysms (Figure 2). In contrast to the dynamic regulation of IL-6 protein expression regulated by Aldo-salt and androgen, MR protein was not altered by orchiectomy before and after Aldo-salt administration (Supplemental Figure 10e−10h).

To investigate whether IL-6 is implicated in Aldo-salt-induced and androgen-mediated aortic aneurysms, 10-month-old male C57BL/6 mice were administered with Aldo-salt with LMT-28 (0.25 mg/kg, oral gavage, once a day) or vehicle (carboxymethyl cellulose [CMC] 0.5%) for four weeks (25). LMT-28 is a recently developed novel small molecule inhibitor and has been shown to specifically target IL-6β to disrupt IL-6α and IL6Rβ dimerization, thus inhibiting IL-6 signaling (25). The efficacy of LMT-28 on IL-6 signaling was appraised by immunostaining of signal transducer and activator of transcription 3 (STAT3) Tyr705 phosphorylation (p-STAT3), an index of the IL-6 signaling activation (25), in the suprarenal aortas by IHC in the mice four weeks after Aldo-salt with LMT-28 or vehicle administration. Supplemental Figure 11A illustrates that treatment of mice with LMT-28 completely abolished p-STAT3 phosphorylation in the aorta in mice administered Aldo-salt.

We next evaluated the effect of LMT-28 on Aldo-salt-induced aortic aneurysms. Treatment of mice with LMT28 partially but significantly suppressed Aldo-salt-induced suprarenal abdominal aortic dilation and progression (Supplemental Figures 11B−11C), the incidence of total aortic aneurysms, including TAA and AAA, and the severity of aortic aneurysms (Supplemental Figures 11E−11G). However, it should be pointed out that the inhibition of LMT-28 on Aldo-salt-induced aortic aneurysms was incomplete compared with gonadal androgen deprivation or targeting AR with ASC-J9 (i..e., Supplemental Figure 11D *vs*. Figures 2D and 4D).

We also determined the effect of LMT-28 on salt retention, serum sodium, and MAP one week before and/or after Aldo-salt administration. In agreement with the effect of orchiectomy, exogenous DHT administration to orchiectomized mice, and downregulation of AR by ASC-J9 (Supplemental Figures 4, 6, and 8), inhibiting IL-6 with LMT-28 did not affect Aldo-salt-induced sodium retention and hypertension (Supplemental Figure 12A−12C). There was no correlation between the internal diameter and serum sodium or MAP levels (Supplemental Figures 12E and 12H). Three was also no difference in serum sodium level or MAP between mice treated with and without LMT-28, regardless of whether they developed aortic aneurysms (Supplemental Figures 12D, 12F,12G, and 12L).

### Identification of T cell receptor signaling by transcriptome profile analysis of the mouse aorta as a link between AR and Aldo-salt-induced aortic aneurysms

Since treatment of mice with LMT-28 completely inhibited the IL-6 signaling but partially blocked Aldo-salt-induced aortic aneurysms (Supplemental Figure 11), we reasoned that additional signaling pathways are likely regulated by androgen and responsible for Aldo-salt-induced aortic aneurysms. To identify these putative signaling pathways in an unbiased way, 10-month-old male C57BL/6 mice were randomly divided into three groups: 1) Aldo-salt administration (to set up a baseline); 2) orchiectomy followed by Aldo-salt administration (to determine the effect of gonadal androgen deprivation); 3) orchiectomy followed by Aldo-salt administration plus DHT pellet implantation (to rescue from gonadal androgen deprivation). Whole aortas were isolated one week after the Aldo-salt administration to study the signaling events that cause, rather than result from, Aldo-salt-induced aortic aneurysms. These samples were subjected to RNA-seq.

Of a total of 18,841 mRNAs detected by RNA-seq, DESeq2 (26) identified 2,359 of them as significantly and differentially abundant (*P* < 0.01) between aortas from the three treatment groups (Figure 5A). Gonadal androgen deprivation caused 298 and 351 mRNAs to be upregulated and downregulated, respectively (Figure 5B). Administration of DHT to orchiectomized mice gave rise to 707 and 1,003 mRNAs being upregulated and downregulated, respectively (Figure 5C). Importantly, the rescue of gonadal androgen deprivation by administration of DHT to orchiectomized mice identified 180 androgen-sensitive mRNAs upregulated by gonadal androgen deprivation but downregulated by administration of DHT to orchiectomized mice (Figures 5D and 5E; Supplemental Table 1) and 150 androgen-sensitive mRNAs downregulated by gonadal androgen deprivation but upregulated (rescued) by administration of DHT to orchiectomized mice (Figures 5G and 5H; Supplemental Table 2).

**Figure 5.**
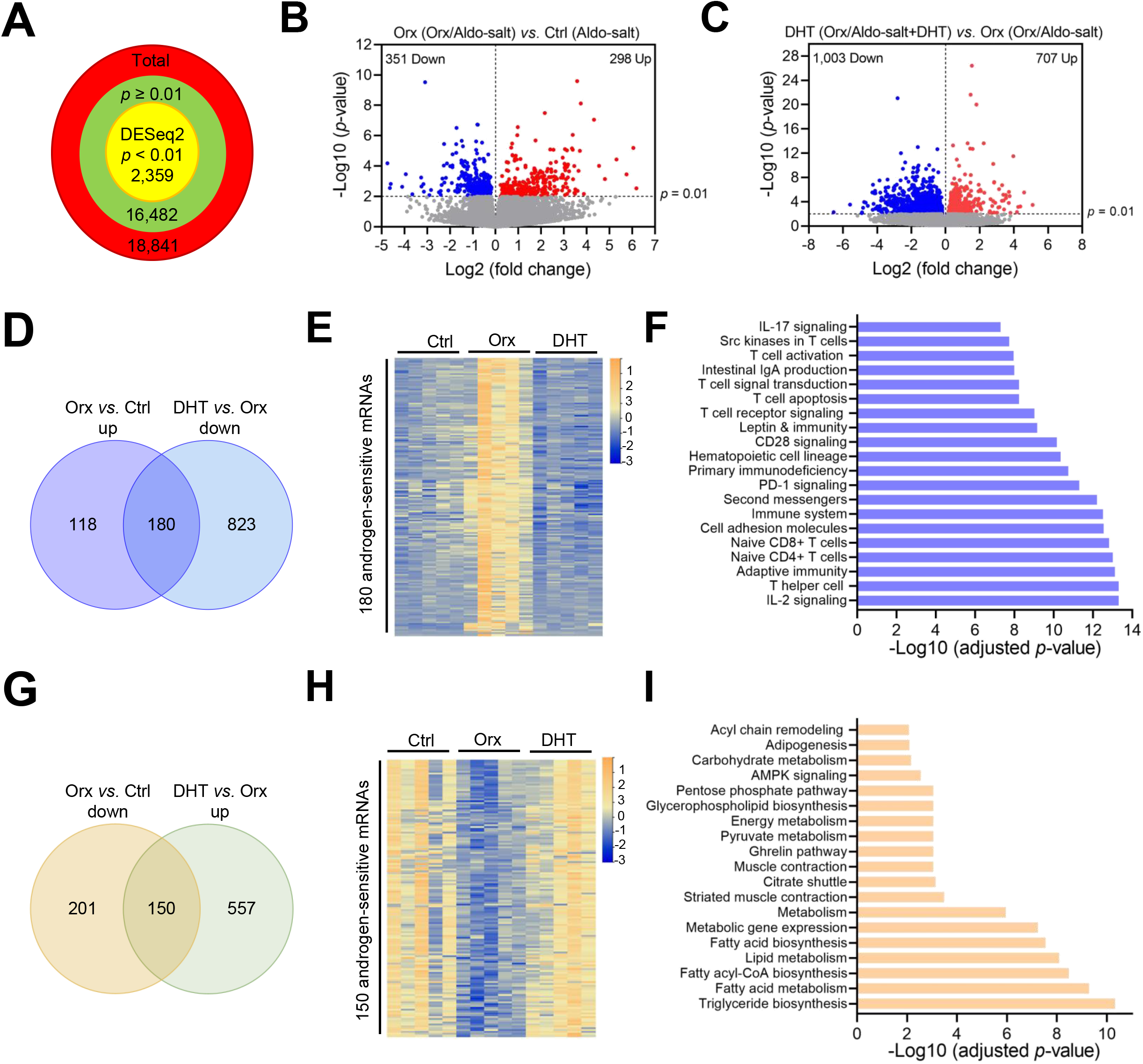
Profiling of aortic transcriptomes reveals T cell receptor signaling as a link between AR and Aldo-salt-induced aortic aneurysms. (**A**) The total number of genes whose mRNAs were detected by RNA-seq and analyzed to be differentially abundant by DESeq2 in the whole aortas from 10-month-old male C57BL/6 mice with or without orchiectomy (orx) followed by one-week Aldo-salt with or without DHT implantation. (**B** and **C**) Volcano plot illustration of the number of genes whose mRNAs with statistical significance (y-axis) versus effect size (fold change, x-axis) in the experiment. (**D** and **G**) Venn diagram identifies the number of genes whose mRNAs are upregulated by orchiectomy but downregulated by DHT (D) and downregulated by orchiectomy but upregulated by DHT (G), respectively. (**E** and **H**) Heatmap of 180 genes whose mRNAs were upregulated by orchiectomy but downregulated DHT (E) and 150 genes whose mRNAs were downregulated by orchiectomy but upregulated by DHT (H), respectively. (**F** and **I**) Pathway enrichment analysis using Enrichr showing the top 20 pathways among the mRNAs that were upregulated by orchiectomy but downregulated by DHT (F) and the 19 pathways among the mRNAs that were downregulated by orchiectomy but upregulated by DHT (I), respectively.

To gain mechanistic insight into the androgen-sensitive mRNAs identified by the RNA-seq analysis, we used Enrichr, a widely-used search engine for comprehensive pathway enrichment analysis (27), to identify the signaling pathways and other functions or processes responsive to androgen and potentially implicated in Aldo-salt-induced aortic aneurysms. The 180 androgen-sensitive mRNAs upregulated by androgen deprivation but downregulated by administration of DHT to orchiectomized mice (Figures 5D and 5E) proved to have 65 overrepresented functional annotations (Supplemental Table 3). Surprisingly, most of these were related to adaptive immunity, particularly T cell receptor signaling (Figure 5F and Supplemental Table 3). In parallel with this finding, the 150 androgen-sensitive mRNAs that were affected by androgen in the opposite direction, downregulated by gonadal androgen deprivation but upregulated by administration of DHT to orchiectomized mice (Figures 5G and 5H), proved to have 19 overrepresented annotations, most of which were associated with triglyceride, fatty acid, and lipid biosynthesis or metabolism (Figure 5I and Supplemental Table 4).

### Splenectomy mitigates Aldo-salt-induced aortic aneurysms and augments PD-1 positive T cells and B cells in the aorta

To investigate whether T cell receptor signaling, identified by the RNA-seq analysis, is involved in Aldo-salt-induced aortic aneurysms,11-13-month-old male C57BL/6 mice were subjected to splenectomy (to remove spleen-derived immune cells, including T cells) or sham operation. After one month of surgery recovery to deplete spleen-derived immune cells, mice were administered Aldo-salt for four weeks to induce aortic aneurysms. Consistent with the previous report (28), splenectomy significantly lowered MAP before and after Aldo-salt administration (Figure 6A), indicating the splenectomy’s success.

**Figure 6.**
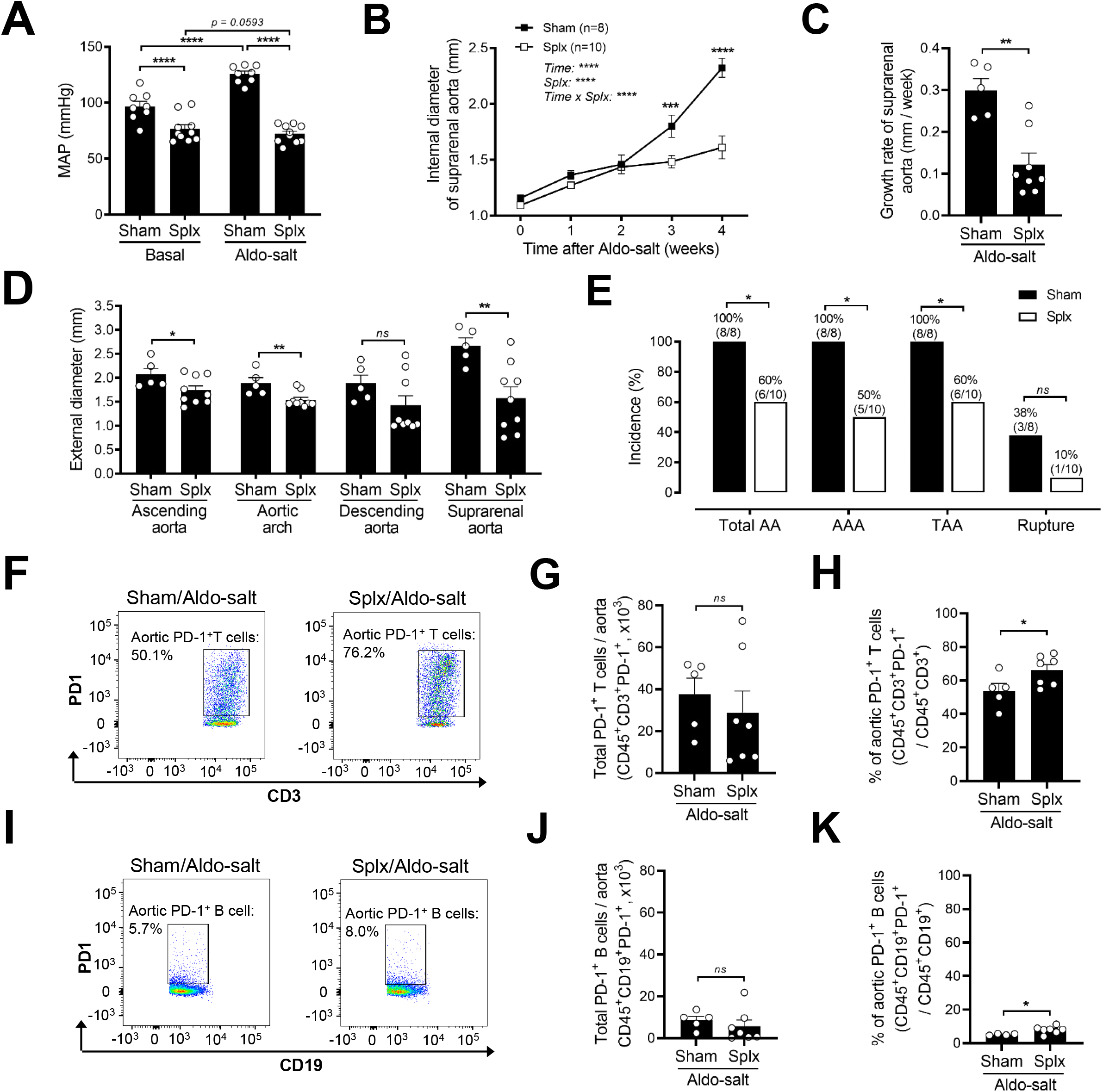
Splenectomy enriches PD-1 positive T cells and B cells in the aorta and mitigates Aldo-salt-induced aortic aneurysms. (**A**) MAP was measured in 11-13-month-old male C57BL/6J mice with splenectomy (splx) or sham operation one week before (basal) and three weeks after Aldo-salt administration. (**B** and **C)** *In vivo* quantification of the maximal intraluminal diameters (B) and growth rate (C) of the suprarenal aortas by ultrasound in splx and sham-operated mice before (0) and weekly after Aldo-salt administration. (**D**) *Ex vivo* measurement of the maximal external diameters of the ascending aorta, aortic arch, descending aorta, and suprarenal aorta by microscopy in isolated aortas four weeks after Aldo-salt administration. (**E**) The incidence of total aortic aneurysms (AA), AAA, TAA, and aortic aneurysm rupture. The percentages of the incidences (number of mice with total AA, AAA, TAA, and rupture / total number of mice) were indicated on the top of each bar. (**F**−**K**) Representative pseudocolor plots and quantitative data of flow cytometry analysis of the total numbers and percentages of PD-1^+^ T cells (F**−**H) and PD-1^+^ B cells (I**−**K) in the whole aortas of splx or sham-operated mice four weeks after Aldo-salt administration. The data were expressed as mean ± SEM and analyzed by two-way ANOVA with multiple comparisons test (A and B), two-tailed unpaired *t*-test (C, D, G, H, J, and K), and two-sided Chi-square test (E). *\*, p < 0*.*05; **, p< 0*.*01; ***, p < 0*.*001*; *\*\*\*\*, p<0*.*0001; ns, not significant*.

The effect of splenectomy on aortic aneurysms was then assessed by ultrasound one week before and weekly after Aldo-salt administration. Splenectomy significantly suppressed Aldo-salt-induced suprarenal aortic dilation and progression (Figures 6B and 6C). Interestingly, such an effect was only found three weeks after Aldo-salt administration (Figure 6B), indicating that spleen-derived immune cells may mainly participate in the progression rather than initiation of Aldo-salt-induced aortic aneurysms. As with its effect on the internal diameters of the suprarenal aorta in response to Aldo-salt (Figures 6B and 6C), splenectomy significantly reduced the external diameters of the ascending aorta, aortic arch, and suprarenal aorta (Figure 6D) and the incidence of Aldo-salt-induced aortic aneurysms, including TAA and AAA (Figure 6E).

To investigate whether splenectomy protects mice from Aldo-salt-induced aortic aneurysms via T cell-mediated vascular inflammation, a single-cell suspension was prepared from the whole aorta (29) of splenectomized and sham-operated mice four weeks after Aldo-salt administration and then subjected to the flow cytometry analysis. As illustrated for the gating strategy in Supplemental Figure 13, single aortic cells were first gated on CD45^+^ to sort leukocytes and then gated on CD3^+^, CD19^+^, F4/80^+^, and Ly6G^+^ to identify T cells, B cells, macrophages, and neutrophils, respectively, and finally gated on PD-1^+^ to identify PD-1^+^-T cells, PD-1^+^-B cells, PD-1^+^-macrophages, and PD-1^+^-neutrophils, respectively. A single-cell suspension was also prepared from the lymph nodes and analyzed by flow cytometry with fluorescence minus one (FMO) to serve as quality controls to identify gating boundaries and ensure the specificity of antibodies. We focused on PD-1^+^ immune cells rather than other immune cells identified by RNA-seq (Figure 5F) because PD-1^+^ immune cells, in contrast to many other immune cells, as an immune checkpoint molecule (15), potentially control rather than exacerbate vascular inflammation (30), thus may underlie the splenectomy protection on Aldo-salt-induced aortic aneurysms.

To our surprise, there was no significant difference in the percentages and total numbers of leukocytes, T cells, B cells, macrophages, and neutrophils in the aortas between splenectomized and sham-operated mice four weeks after Aldo-salt administration (Supplemental Figure 14). However, there was a significant increase in the percentages but not the total numbers of PD-1^+^ T cells and PD-1^+^ B cells in the aortas in splenectomized mice compared to that in sham-operated mice four weeks after Aldo-salt administration (Figures 6F−6K). In contrast, splenectomy did not affect PD-1^+^ macrophages (Supplemental Figures 15A−15C). Intriguingly, splenectomy decreased rather than increased the percentages and total numbers of PD-1^+^ neutrophils (Supplemental Figures 15D−15F).

### AR binds to the PD-1 promoter and suppresses its mRNA and protein expression in the spleen in mice administered Aldo-salt

While the resource of the increase in PD-1^+^ T and PD-1^+^ B cells in the aorta in splenectomized mice remains determined, the results that splenectomy inhibited Aldo-salt-induced aortic aneurysms concomitant with increased PD-1^+^ T and PD-1^+^ B cells in the aorta (Figures 6F−6K) suggest that PD-1^+^ T cells and B cells may play a role in Aldo-salt-induced aortic aneurysms. However, it should be pointed out that whether PD-1^+^ T cells and B cells are regulated by androgen is unknown since the splenectomy experiment was conducted in intact mice but not orchiectomized mice. As a result, the mechanism by which androgen regulates PD-1 is completely unknown. To address these important questions, we performed a series of experiments to investigate whether and how PD-1^+^ T cells and B cells are regulated by androgen in the spleen robustly. We focused on the spleen rather than other immune system organs because it has been shown that the spleen’s weight is increased by orchiectomy in mice (31), and more importantly, we (Figures 6A−6E) and others (32) demonstrated that the spleen is critical for Ang II- and Aldo-salt-induced aortic aneurysms.

First, to investigate a potential causal role of Aldo-salt-induced aortic aneurysms, we examined PD-1 protein expression by IHC in the spleens from 10-month-old male C57BL/6J mice with orchiectomy or sham operation ten days after Aldo-salt administration. Representative immunostaining and quantitative data demonstrate that PD-1 protein was mostly expressed in the white pulp of the spleen (Figures 7A and 7B), an immunological region of the spleen that mainly consists of lymphocytes, including T cells and B cells (33). Importantly, PD-1 protein was markedly upregulated in the spleen in orchiectomized mice relative to sham-operated mice (Figures 7A and 7B), indicating that androgen suppresses PD-1 protein expression in the spleen. To further define the role of androgen in regulating PD-1 protein expression, we tested whether the rescue of gonadal androgen deprivation by exogenous DHT administration to 10-month-old male orchiectomized mice was able to re-suppress PD-1 protein expression in the spleen four weeks after Aldo-salt administration. Consistent with the effect of orchiectomy on PD-1 protein expression in the spleen (Figures 7A and 7B), exogenous DHT administration to orchiectomized mice potently suppressed PD-1 protein upregulation in the spleen in orchiectomized mice (Figures 7C and 7D) to a level that was even lower than in sham-operated mice administered Aldo-salt (Figures 7C and 7D *vs*. Figures 7A and 7B).

**Figure 7.**
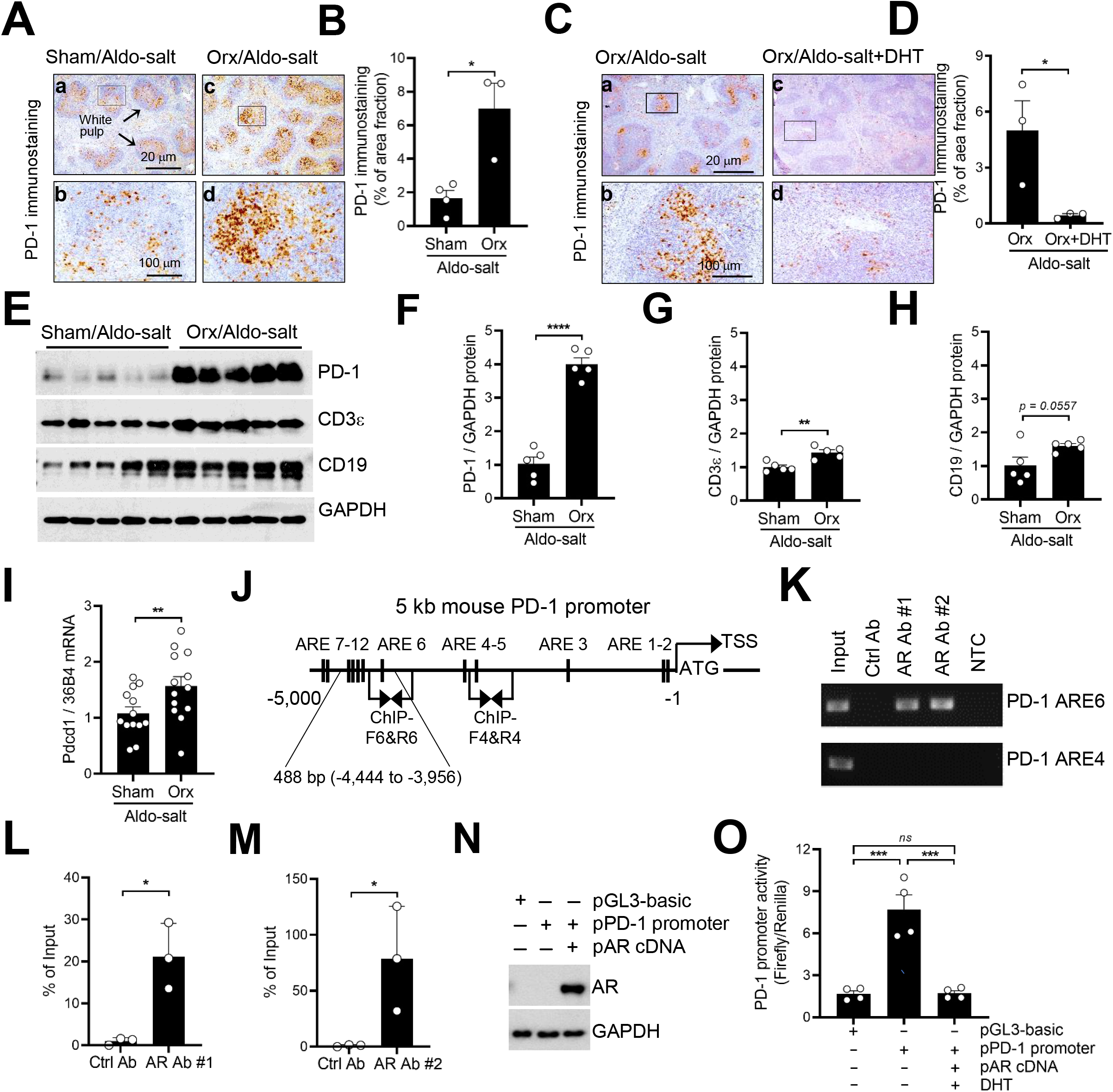
Androgen suppresses PD-1 mRNA and protein expression in the spleen in mice administered Aldo-salt. (**A−D**) Representative immunostainings and quantitative data of PD-1 protein expression in the spleens from 10-month-old orchiectomized (orx) and sham-operated C57BL/6J mice ten days after Aldo-salt administration (A and B) or orchiectomized mice four weeks after Aldo-salt with or without DHT pellet implantation (C and D). The percentage of areas fraction = (the PD-1 positive area / the area of fields of view) x 100%. The data were calculated from five fields of view randomly photographed per splenic section per mouse. (**E−H**) Representative Western blots and quantitative data of PD-1, CD3ε, CD19, and GAPDH protein expression in the spleens from 10-month-old male mice with orx or sham operation ten days after Aldo-salt administration. (**I**) Pdcd1 (the gene codes PD-1) mRNA expression in the spleens from 10-month-old male mice with orx or sham operation ten days after Aldo-salt administration. (**J**) A schematic diagram of the 12 androgen response elements (ARE) in the 5 kb mouse PD-1 promoter. TSS, transcription start site. ATG, translation start codon. ChIP-F, ChIP-PCR forward primers. ChiP-R, ChIP-PCR reverse primers. (**K−M**) Representative and quantitative ChIP-PCR with the control Ab, AR Ab #1, and AR Ab #2 in the mouse spleen. NTC, no template control. (**N** and **O**) Expression of AR protein in HEK293 cells suppressed the PD-1 promoter activity. The data were expressed as mean ± SEM and analyzed by two-tailed unpaired *t*-test (B, D, F−H, I, L, and M) and one-way ANOVA with multiple comparisons test (O). *\*, p < 0*.*05; **, p < 0*.*01; ***, p < 0*.*001*; *\*\*\*\*, p<0*.*0001; ns, not significant*.

Second, to quantify the effect of androgen on PD-1 protein expression in the spleen, we determined PD-1 protein expression by Western blot analysis in the spleens from 10-month-old orchiectomized or sham-operated C57BL/6J mice ten days after Aldo-salt administration. In agreement with our IHC data (Figures 7A and 7B), PD-1 protein was upregulated up to 4-fold in the spleen in orchiectomized mice relative to sham-operated mice (Figures 7E and 7F). To determine whether orchiectomy-induced PD-1 protein upregulation in the spleen results from T cells or B cells, we examined protein expressions of CD3ε, a T cell marker, and CD19, a B cell marker, in the same spleen lysate. Interestingly, CD3ε but not CD19 protein was significantly upregulated by orchiectomy in the spleen (Figures 7E, 7G, and 7H), indicating that orchiectomy-induced PD-1 upregulation may be mainly attributed to T cells but not B cells in the spleen. Moreover, the fold upregulation of CD3ε protein induced by orchiectomy in the spleen was 1.4 folds, much less than orchiectomy-induced PD-1 protein upregulation (4 folds) in the spleen (Figure 7F *vs*. Figure 7G). Thus, an increase in T-cell numbers (i.e., CD3ε protein upregulation) cannot fully account for PD-1 protein upregulation in T cells, and additional mechanisms (i.e., upregulation of PD-1 protein in T cells) likely also contribute to a higher PD-1 protein expression in the spleen in orchiectomized mice administered Aldo-salt.

Third, to further determine the cellular resource of PD-1 protein upregulation by orchiectomy in the spleen, a single cell suspension was prepared from the spleens in 10-month-old male C57BL/6J mice with orchiectomy or sham operation ten days after Aldo-salt administration and subjected to flow cytometry analysis of splenic PD-1^+^ T cells (CD45^+^CD3^+^PD-1^+^) and PD-1^+^ B cells (CD45^+^CD19^+^PD-1^+^; Supplemental Figure 16). Intriguingly, although orchiectomy did not affect the total number and percentage of splenic T cells (Supplemental Figures 17D−17F), it significantly increased the total number, not the percentage, of splenic PD-1^+^ T cells (Supplemental Figures 17I−17K). In contrast, although orchiectomy significantly elevated the total number, not the percentage, of splenic B cells (Supplemental Figures 17D, 17G, and 17H), it did not alter the total number and percentage of splenic PD-1^+^ B cells (Supplemental Figures 17L−17N). Thus, these data are consistent with our Western blot data (Figures 7E−7H), suggesting that orchiectomy-induced PD-1 protein upregulation in the spleen mainly results from splenic T cells, but not B cells, in mice administered Aldo-salt.

Fourth, to investigate whether PD-1 is regulated by androgen at the transcription level in the spleen in mice administered Aldo-salt, which may mechanistically underlie orchiectomy-induced PD-1 protein upregulation, we determined Pdcd1 (the gene that codes PD-1) mRNA expression by quantitative real-time PCR in the spleen from 10-month-old male C57BL/6J mice with orchiectomy or sham operation ten days after Aldo-salt administration. As shown in Figure 7I, Pdcd1 mRNA was significantly upregulated by orchiectomy in the spleen in orchiectomized mice compared to sham-operated mice, indicating that androgen suppresses splenic PD-1 protein expression via at least in part by suppressing Pdcd1 mRNA expression in the spleen in mice administered Aldo-salt.

Fifth, to investigate the mechanism by which androgen suppresses Pdcd1 mRNA expression in the spleen in mice administered Aldo-salt, we investigated whether AR binds to the PD-1 promoter to suppress its transcription. To address this important mechanistic issue, we analyzed a 5 kb mouse PD-1 promoter DNA sequence to identify androgen response elements (ARE) containing AGAACA or TGTTCT hexamers, which is sufficient to be bound by AR (34). We found 12 putative ARE in the 5 kb mouse PD-1 promoter (Figure 7J). To determine whether AR can bind to these putative ARE hexamers in the spleen, we performed chromatin immunoprecipitation (ChIP) assays using the mouse spleen with two different commercially available but epitope-distinct ChIP-grade anti-AR antibodies (to ensure the specificity of anti-AR antibodies) and two sets of ChIP-PCR primers that specifically amplify ARE4 and ARE6 in the mouse PD-1 promoter (Figure 7J). As shown in Figures 7K−7M), both anti-AR antibodies, but not a control antibody (nonspecific rabbit IgG), were capable of pulling down the chromatins containing ARE6 but not ARE4, indicating that AR can bind to the ARE6 but not ARE4 in the PD-1 promoter in the spleen.

Sixth, to investigate whether the binding of AR to the PD-1 promoter via ARE6 inhibits its transcriptional activity, we subcloned a 488 bp PD-1 mouse promoter (−4,444 to -3,956 bp relative to the transcription start site (TSS)) containing ARE6 to ARE10 (Figure 7J) into a pGL3-basic firefly luciferase report vector. The pGL3-basic-PD-1 promoter construct was co-transfected with the pRL-TK control vector (to express Renilla luciferase to control transfection efficiency) and a pcDNA Flag-M4-AR construct (35) (to express human AR protein) or a pcDNA3.1 vector (to maintain the equal amount of cDNA during transfection) into HEK293 cells. We used HEK293 cells rather than splenic cells because HEK293 cells, but not splenic cells, have a high transfection efficiency. Moreover, HEK293 cells expressed little AR protein (Figure 7N), which provides an additional advantage in assessing the effect of exogenous AR on the PD-1 promoter activity. Dual luciferase assays illustrate that the -488 bp PD-1 promoter exhibited a 4.6-fold higher luciferase activity relative to the pGL3-basic vector (Figure 7O), indicating that the cloned -488 bp PD-1 promoter can drive PD-1 transcription. Importantly, co-transfection of the PD-1 promoter-luciferase constructs with the human AR cDNA construct completely abolished the PD-1 promoter activity in the presence of DHT (to activate AR; Figure 7O), indicating that the binding of AR to the PD-1 promoter can suppress its transcriptional activity.

Seventh, to explore how orchiectomy-induced PD-1 upregulation in the spleen contributes to its protection against Aldo-salt-induced aortic aneurysms, the blood cells were collected from 10-month-old male C57BL/6J mice with orchiectomy or sham operation ten days after Aldo-salt administration and then subjected to the flow cytometry analysis of PD-1^+^ T cells (CD45^+^CD3^+^PD-1^+^) and PD-1^+^ B cells (CD45^+^CD19^+^PD-1^+^; Supplemental Figure 18). In line with its effect on PD-1^+^ T cells in the spleen (Supplemental Figures 17D−17F and 17I−17H), orchiectomy increased the total and percentage of PD-1^+^ T cells in the blood (Supplemental Figures 19I−19K), although it did not change the total and percentage of T cells (Supplemental Figures 19D−19F). Also, in agreement with its effect on PD-1^+^ B cells in the spleen (Supplemental Figures 17D, 17G−17H, and 17L−17N), orchiectomy did not alter the total and percentage of PD-1^+^ B cells in the blood (Supplemental Figures 19L−19N), even though it increased the total and percentage of B cells in the blood (Supplemental Figures 19D, 19G, and 19H). Together, these results suggest that orchiectomy mobilizes PD-1^+^ T cells, but not PD-1^+^ B cells, from the spleen to the blood, thereby protecting mice from Aldo-salt-induced aortic aneurysms.

Finally, to investigate whether orchiectomy-induced PD-1 upregulation also occurs in other immune system organs, we examined PD-1 protein expression by IHC and Western blot analysis in the periaortic lymph nodes from 10-month-old C57BL/6J mice with orchiectomy or sham operation ten days after Aldo-salt administration. IHC analysis showed an increasing trend toward PD-1 immunostaining in the periaortic lymph nodes in orchiectomized mice compared to sham-operated mice (Supplemental Figures 20A and 20B). Western blot analysis confirmed the IHC data and illustrated a significant PD-1 protein upregulation in the periaortic lymph nodes in orchiectomized mice compared to sham-operated mice (Supplemental Figures 20C and 20D). Hence, these results suggest that the suppression of androgen on PD-1 expression is not limited to the spleen and can occur in other immune organs, potentially contributing to Aldo-salt-induced aortic aneurysms.

### Blockade of the immune checkpoint with anti-PD-1 antibody reinstates Aldo-salt-induced aortic aneurysms in orchiectomized mice

To investigate whether PD-1 signaling mediates Aldo-salt-induced and androgen-mediated aortic aneurysms, 10-month-old C57BL/6J male mice were subjected to orchiectomy (to induce gonadal androgen deprivation and PD-1 upregulation in T cells) and then administered with Aldo-salt (to induce aortic aneurysms) with a specific rat anti-mouse PD-1 antibody (to inhibit PD-1 signaling in T cells) or an isotype control antibody (200 µg/mice, i.p. injection, twice a week for eight weeks) (36).

The effect of the anti-PD-1 antibody on Aldo-salt-induced aortic aneurysms was first examined by ultrasound in orchiectomized mice one week before and weekly four weeks after the Aldo-salt administration. Intraperitoneal injection of anti-PD-1 antibody partially but significantly reinstated Aldo-salt-induced suprarenal aortic dilation compared with the control antibody four weeks after Aldo-salt administration (anti-PD-1 Ab *vs*. Ctrl Ab, *p* < 0.01). However, the effect of the anti-PD-1 antibody on Aldo-salt-induced suprarenal aortic dilation was less potent and only distinguishable from that of the control antibody three weeks after Aldo-salt administration (Figure 8A), indicating that PD-1 is mainly implicated in the progression rather than the initiation of Aldo-salt-induced AAA. To further evaluate the effect of the anti-PD-1 antibody on the progression of aortic aneurysms, which have more clinical relevance, these mice were continuously administered Aldo-salt with anti-PD-1 or control antibodies for up to eight weeks. Compared with its effect from week 1 to week 4, the anti-PD-1 antibody did not lead to a further suprarenal aortic dilation from week 5 to week 8 (Figure 8A). To investigate whether intraperitoneal injection of anti-PD-1 antibody affects Aldo-salt-induced TAA, we measured the internal diameters of the aortic arch by ultrasound six weeks after Aldo-salt with anti-PD-1 or control antibody administration. A similar but more potent effect of the anti-PD-1 antibody was found on Aldo-salt-induced aortic arch dilation compared with its effect on the suprarenal aortic dilation (anti-PD-1 Ab *vs*. Ctrl Ab, *p* < 0.0001; Figure 8B).

**Figure 8.**
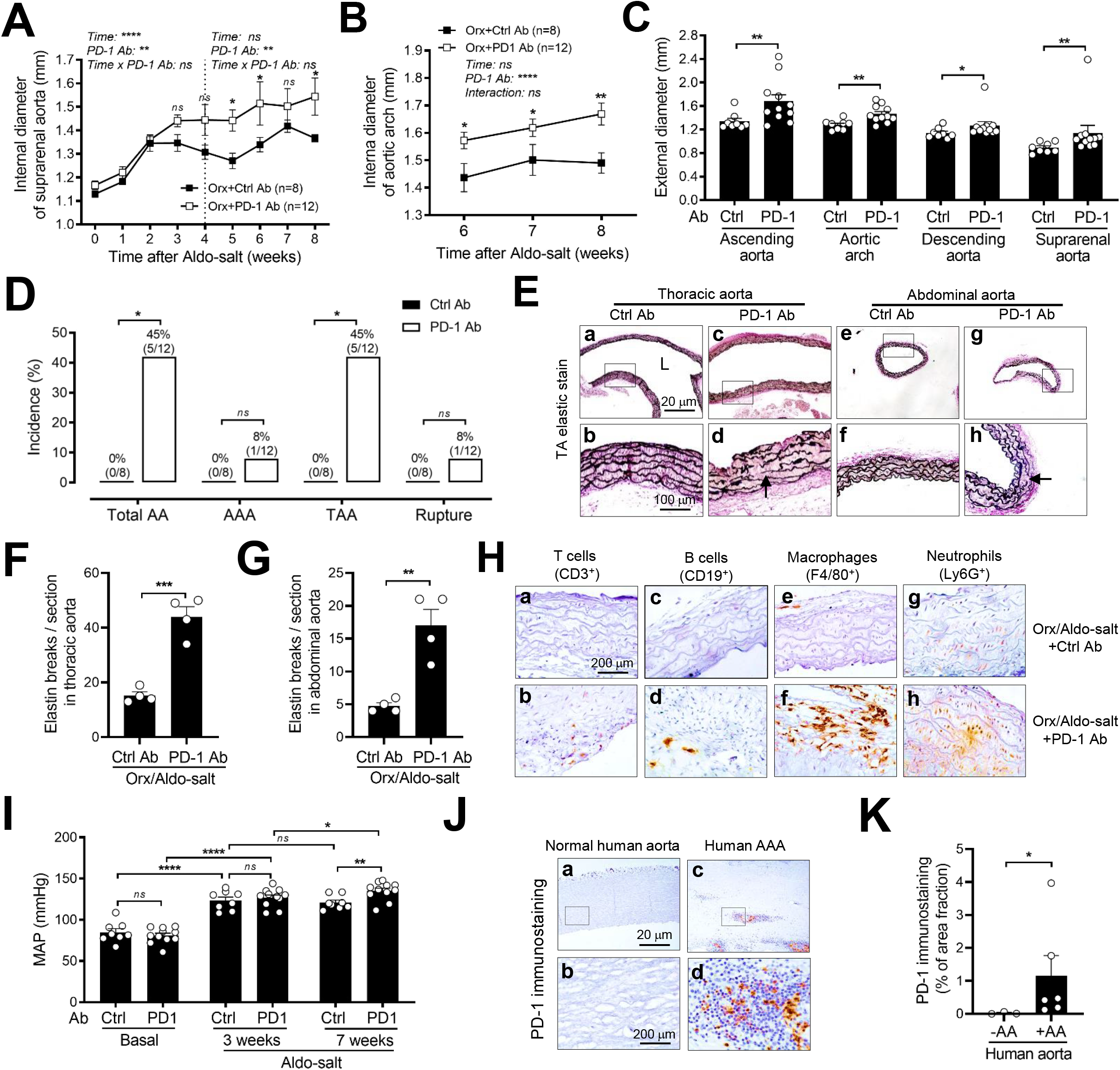
Intraperitoneal injection of anti-PD-1 antibody reinstates Aldo-salt-induced aortopathy in orchiectomized mice. (**A** and **B**) *In vivo* quantification of the maximal intraluminal diameters of the suprarenal aorta (A) and aortic arch (B) by ultrasound in 10-month-old male C57BL/6J mice with orchiectomy (orx) one week before (0) and weekly after Aldo-salt with anti-PD-1 or Ctrl antibodies (Ab). (**C**) *Ex vivo* measurement of the external diameters of the ascending aorta, aortic arch, descending aorta, and suprarenal aorta by microscopy in aortas eight weeks after Aldo-salt with anti-PD-1 or Ctrl Abs. (**D**) The incidences of AAA, TAA, and aortic aneurysm rupture were calculated based on the pathology of the aorta. (**E**−**G**) Representative and quantitative Verhoeff-Van Gieson staining of elastin in longitudinal sections of the thoracic aortas (E and F) and crosssections of the abdominal aortas (E and G) in orx mice with anti-PD-1 or ctrl Ab eight weeks after Aldo-salt administration. (**H**) Representative immunostaining of T cells, B cells, macrophage, and neutrophils in the thoracic aortas in orx mice with anti-PD-1 or ctrl Abs eight weeks after Aldo-salt administration. (**I**) MAP was measured by tail cuff one week before (basal) and three and seven weeks after Aldo-salt administration. (**J** and **K)** Representative immunostainings and quantitative data of PD-1 protein expression in the human aortas with or without aortic aneurysms. The data were expressed as mean ± SEM and analyzed by two-way ANOVA with multiple comparisons test (A, B, and I), two-tailed unpaired *t*-test (C, F, G, and K), and two-sided Chi-square test (D). *\*, p<0*.*05; **, p<0*.*01; ***, p<0*.*001; ***, p<0*.*001; ns, not significant*.

The effect of the anti-PD-1 antibody on Aldo-salt-induced aortic aneurysms was further investigated by morphometric analysis of the external diameters of the ascending aorta, aortic arch, descending aorta, and suprarenal aorta in orchiectomized mice eight weeks after Aldo-salt with the anti-PD-1 or control antibody administration. In agreement with its effect on the internal diameters of the suprarenal aorta and aortic arch (Figures 8A and 8B), intraperitoneal injection of anti-PD-1 antibody significantly increased the external diameters of the ascending aorta, aortic arch, descending aorta, and suprarenal aorta (Figure 8C). Moreover, pathological analysis of the aorta revealed that, of 12 mice treated with the anti-PD-1 antibody, 5 developed aortic aneurysms (45%), including 1 AAA (8%), 5 TAA (45%), and 1 aortic aneurysm rupture (8%). In contrast, none of 8 mice injected with the control antibody developed aortic aneurysms (Figure 8D). Thus, the anti-PD-1 antibody seems to have more effect on TAA than AAA in orchiectomized mice administered Aldo-salt.

One of the fundamental pathological characteristics of human aortic aneurysms is aortic elastin fiber fragmentation, which leads to aortic dissection and aortic aneurysm rupture (1, 2, 37). To investigate whether the anti-PD-1 antibody affects Aldo-salt-induced aortic elastin fiber fragmentation, the aortas were isolated from orchiectomized mice eight weeks after Aldo-salt with anti-PD-1 antibody or control antibody administration, cut into a longitudinal section of the thoracic aorta and a crosssection of the abdominal aorta, and subjected to Verhoeff-Van Gieson staining (13, 14). Representative photographs and quantitative data demonstrate an explicit increase in thoracic and abdominal aortic elastin fiber breakages in orchiectomized mice treated with the anti-PD-1 antibody and developed aortic aneurysms but not in mice administered with the control antibody and did not develop aortic aneurysms (Figures 8E−8G).

Given the pivotal role of the PD-1/PD-L1 pathway in immune checkpoint and vascular inflammation (15, 30, 38), we anticipated that blockage of the immune checkpoint with anti-PD-1 antibody might increase Aldo-salt-induced vascular inflammation, thus promoting Aldo-salt-induced aortic aneurysms. To test this possibility, the same sets of the longitudinal section of the thoracic aorta that were subjected to Verhoeff-Van Gieson staining (Figure 8E) were immunostained with anti-CD3ε, CD19, F4/80, and Ly6G immunostaining to identify T cells, B cells, macrophages, and neutrophils, respectively. As shown in Figure 8H, T cells, B cells, macrophages, and neutrophils were readily detectable in the thoracic aorta from mice administered the anti-PD-1 antibody but rarely found in the thoracic aorta from mice administered the control antibody. These results suggest that intraperitoneal injection of anti-PD-1 antibody to orchiectomized mice reinstates Aldo-salt-induced aortic aneurysms via unchecked vascular inflammation.

We also explored the effect of the anti-PD-1 antibody on blood pressure by tail cuff one week before (basal) and three or seven weeks after Aldo-salt administration. Interestingly, the anti-PD-1 antibody exhibited a pro-hypertensive effect relative to the control antibody in a time-dependent manner: it did not affect MAP before and three weeks after Aldo-salt administration, whereas it exacerbated Aldo-salt-induced hypertension seven weeks after Aldo-salt administration (Figure 8I). These findings suggest that the anti-PD-1 antibody may provoke Aldo-salt-induced aortic aneurysms via blood pressure. Paradoxically, this possibility is supported by a significant correlation in the internal diameters of the aortic arch and MAP (Supplemental Figure 21A) but is not supported by no difference in MAP between orchiectomized mice treated with the anti-PD-1 antibody regardless of whether they developed aortic aneurysms (Supplemental Figure 21B).

Finally, to explore the potential involvement of PD-1 in human aortic aneurysms, we examined PD-1 protein expression by IHC in the abdominal aorta specimens from human patients with or without AAA. Figures 8J and 8K illustrate that PD-1 immunoreactivity was barely detectable in normal abdominal aortas but could be found in human AAA. These results suggest that PD-1 may play a role in human aortic aneurysms. Future studies are required to address this important question.

## Discussion

The major new findings from current studies include 1) Aldo-salt-induced aortic aneurysms mimic human aortic aneurysms and exhibit a strong sex dimorphism; 2) gonadal androgen deprivation via orchiectomy abolishes Aldo-salt-induced aortic aneurysms; 3) restoration of androgen in orchiectomized mice by DHT pellet implantation reinstates Aldo-salt-induced aortic aneurysms; 4) downregulation of AR by ASC-J9 ameliorates Aldo-salt-induced aortic aneurysms; 5) inhibition of IL-6 signaling by LMT-28 attenuates Aldo-salt-induced and androgen-mediated aortic aneurysms; 6) RNA-seq identifies T cell receptor signaling as a link between AR and Aldo-salt-induced aortic aneurysms; 7) splenectomy mitigates Aldo-salt-induced aortic aneurysms and augments PD-1^+^T cells and B cells in the aorta; 8) orchiectomy increases PD-1 expression in T cells but not B cells, macrophages, and neutrophils in the spleen in mice administered Aldo-salt; 9) mechanistically, AR binds to the PD-1 promoter and suppresses its mRNA and protein expression in the spleen; and 10) blockade of the immune checkpoint with anti-PD-1 antibody restores Aldo-salt-induced aortopathies in orchiectomized mice.

It has long been recognized that androgen is implicated in cardiovascular diseases in addition to its effect on biological sex determination (39). It was also well documented that men with low androgen have a high prevalence of cardiovascular diseases (39), including aortic aneurysms (40). Whether low androgen causes human aortic aneurysms remains investigated, several lines of evidence from animal studies demonstrate that androgen aggravates rather than protects animals from aortic aneurysms. For instance, Henriques et al. reported that orchiectomy reduced AAA incidence from 85% to 18% in apolipoprotein E-deficient mice (ApoE^-/-^) infused with Ang II (8). Cho et al. reported that orchiectomy suppressed the external diameters of the infrarenal aortas compared with sham operation in Sprague Dawley rats infused with elastase (11). Huang et al. reported that global or cell-specific deletion of AR prevented ApoE^-/-^mice from Ang II-induced AAA (10). In agreement with these findings, the current study, using three different approaches, orchiectomy, restoration of androgen in orchiectomized mice by DHT pellet implantation, and downregulation of AR with ASC-J9, demonstrates that both androgen and AR are also critical for sexual dimorphism of aortic aneurysms in 10-month-old C57BL/6 mice administered Aldo-salt (Figure 2−4). Since the Aldo-salt AAA mouse model is independent of the Ang II and elastase AAA murine models (13), our study provides additional evidence that androgen aggravates but does not protect aortic aneurysms.

One of the most novel findings from the current study is that androgen aggravates Aldo-salt-induced aortic aneurysms, at least partially, via suppressing PD-1^+^ T cells. Several lines of evidence from the current study support this potential mechanism. First, using RNA-seq, an unbiased global gene expression profiling analysis, we identified 180 genes upregulated by gonadal androgen deprivation but downregulated by DHT pellet implantation in the aortas in orchiectomized mice one week after Aldo-salt administration (Figures 5D and 5E and Supplemental Tables 1). Of particular interest is these 180 androgen target genes were mostly mapped to T-cell receptor signaling (Figure 5F and Supplemental Tables 3). Although it has long been known that androgen can suppress T cells (20), and it is well documented that some subsets of T cells (i.e., CD4, Th17, and regulatory T cells (Treg)) participate in CaCl_2_-, elastase-, or Ang II-induced AAA (41-43), whether T cell receptor signaling links androgen and aortic aneurysms has not been demonstrated in any aortic aneurysm animal models. To the best of our knowledge, the current study is the first attempt to address this important question in the field. Second, in agreement with the potential role of T cells in AAA (41-43), we demonstrate that altering immune cells by splenectomy mitigates Aldo-salt-induced aortic aneurysms (Figures 6A−6E) and, more importantly, we found that PD-1^+^ T and PD-1^+^ B cells, but neither total T cells and total B cells nor PD-1^+^ macrophages and PD-1^+^ neutrophils, were increased by splenectomy in the aorta in mice administered Aldo-salt (Figures 6F−6K and Supplemental Figures 13−15), which provide the first experiment hint that PD-1^+^-T or PD-1^+^ B cells may respond to androgen and are implicated in Aldo-salt-indued aortic aneurysms. This possibility is also supported by our RNA-seq data, where PD-1 signaling was identified (Figure 5F). Third, using several distinct approaches, including IHC, Western bots, and flow cytometry, we consistently demonstrate that PD-1^+^-T cells but not PD-1^+^-B cells were upregulated by orchiectomy in the spleen, blood, and lymph nodes in mice administered Aldo-salt (Figures 7A−7H and Supplemental Figures 16−20). Finally, perhaps most importantly, we demonstrate that intraperitoneal injection of orchiectomized mice with an anti-PD1 antibody partially but significantly reinstates Aldo-salt-induced aortic aortopathies, including elastin degradation, vascular inflammation, and aortic aneurysms (Figure 8). Although intraperitoneal injection of anti-PD-1 antibody affects both TAA and AAA (Figures 8−8C), it affects TAA more potently than AAA (Figure 8D). While the reasons for this intriguing finding remain investigated, it is noted that the results from the current study suggest that PD-1^+^ T cells, as an immune checkpoint, restrict Aldo-salt-induced aortic aneurysms, which is consistent with the role of PD-1^+^ T cells in many other diseases, including giant cell arteritis, cancer, and atherosclerosis (30). However, it should be pointed out that this finding completely contrasts with a recent study that the intraperitoneal injection of a humanized PD-1 antibody inhibited rather than aggravated CaCl_2_-induced AAA progression (44). The discrepancy between the current study (Figure 8) and the previous study (44) may be attributed to using different aortic aneurysm animal models (CaCl_2_ *vs*. Aldo-salt), anti-PD-1 antibodies(anti-human PD-1 *vs*. an anti-mouse PD-1), and mice (intact *vs*. orchiectomized). Further studies are needed to clarify this issue.

PD-1 functions as an immune checkpoint to inhibit T-cell activation via ligation with its ligands PD-L1 (30). PD-1 is exclusively expressed in activated immune cells, most importantly in T cells, whereas PD-L1 is broadly expressed in antigen-presenting cells (i.e., macrophages), cancer cells, and endothelial cells (30). It has long been recognized that sex hormones regulate PD-1 and PD-L1 expression. In particular, PD-1 is upregulated by estrogen in regulatory T cells (Tregs), macrophages, B cells, and dendritic cells in mice (45), while PD-L1 is downregulated by androgen via suppressing INFγ in cancer cells (46). However, whether androgen can modulate PD-1 expression has not been reported in any immune cells. As a result, the mechanism by which androgen suppresses PD-1 expression is completely unknown. In this respect, the current study discovered that androgen suppresses PD-1 expression via its receptor, AR, binding to the ARE in the PD-1 promoter in the spleen and inhibiting its transcriptional activity. The AR is well known to bind to a 15-bp bipartite palindromic ARE (AGAACAnnnTGTACC) to control AR target gene expression (34). However, a recent study reported that more than 50% of AR-binding sites do not contain the established 15 bp ARE and only contain a 6-bp (also known as “half-site” ARE), which is sufficient to serve as a cis-regulatory element in the PD-1 target genes (34). Consistent with this report, we found 12 putative 6-bp half-site AREs, but not 15-bp ARE, in the 5-kb mouse PD-1 promoter (Figure 7J). By ChIP assay, we demonstrate that AR can bind to the mouse PD-1 promoter via ARE6 in the spleen (Figures 7J−7M). Moreover, co-transfection of AR with a 500-bp PD-1 promoter containing ARE6 inhibits PD-1 promoter activity in HEK293 cells (Figures 7N−7O). However, it is worth pointing out that the fold of orchiectomy-induced PD-1 protein upregulation is much larger than its effect on PD-1 mRNA upregulation in the spleen in mice administered Aldo-salt (Figure 7E *vs*. 7I). Thus, additional mechanisms by which androgen suppresses PD-1 expression (i.e., posttranslational modifications (47)) cannot be ruled out.

Since androgen, a steroidal hormone, has a pleiotropic effect on many organs and systems via genomic and non-genomic mechanisms (39), androgen likely aggravates Aldo-salt-induced aortic aneurysms via different mechanisms rather than suppressing PD-1^+^ T cells only. Consistent with this concept, it has been shown that androgen promotes Ang II-induced AAA by modulating Ang II type 1A receptor (AT1aR), IL-1α, and TGFβ1 expression in the aorta in mice (9, 10). In line with these findings, the current study demonstrates that IL-6, a pleiotropic cytokine, responds to Aldo-salt, is regulated by androgen, and is implicated in Aldo-salt-induced aortic aneurysms (Supplemental Figures 9K, 10, and 11). Although both PD-1 and IL-6 are involved in Aldo-salt-induced and androgen-mediated aortic aneurysms, the mechanism by which PD-1 and IL-6 respond to androgen and mediate Aldo-salt-induced aortic aneurysm appear quite different as PD-1 is mostly expressed in T cells (Figures 6F−6K), whereas IL-6 is predominantly expressed in smooth muscle cells in the tunica media of the abdominal aorta (Supplemental Figure 10). Since it has been shown that STAT3 and STAT4, the downstream effects of IL-6 signaling (25), regulate PD-1 expression in T cells (48), IL-6 may also likely respond to androgen and mediate Aldo-salt-salt-induced aortic aneurysms via regulating PD-1 expression in T cells in the spleen. Further studies are needed to explore this possibility.

There are some limitations in the current studies that need to be addressed in future studies. First, most of the experiments in the current studies were conducted in mice administered Aldo-salt, excluding mice not administered Aldo-salt. Thus, it is unclear how the Aldo/MR signaling coordinates with the androgen/AR signaling to regulate PD-1 expression in T cells since the role of the Aldo/MR signaling in regulating immune cells, including T cells, is well recognized (49). Second, the current study has not identified a subtype of PD-1^+^-T cells (i.e., CD4^+^, CD8^+^, or Treg) that respond to androgen and are implicated in Aldo-salt-induced aortic aneurysms. Third, it is unclear whether intraperitoneal injection of anti-PD-1 antibodies also exacerbates Aldo-salt-induced aortic aneurysms in intact mice, as in orchiectomized mice.

Given that the immune checkpoint inhibitors (i.e., anti-PD-1 antibody) have been successful in cancer treatment and have revolutionized the cancer research field (15), there have been an increasing number of patients with cancer applied to immune checkpoint therapy (50). However, as with other cancer therapies, immune checkpoint therapy has suffered from serious immune-related cardiovascular adverse events, including autoimmune myocarditis, pericarditis, and vasculitis (50). In line with these clinical observations, the current study demonstrates for the first time that the treatment of mice with anti-PD-1 antibody can reinstate Aldo-salt-induced aortopathies (Figure 8), indicating an increased risk of developing aortic aneurysms for patients with immune checkpoint therapy. Consistent with this possibility, a recent case report shows that a 57-year-old man with lung adenocarcinoma treated with chemotherapy and immune checkpoint blockade developed inflammatory TAA (51). Hence, the current study suggests that cancer patients predisposed to the risk factors of aortic aneurysms (i.e., male, aging, and smoking) may have a chance to develop aortic aneurysms during immune checkpoint therapy and should be advised to screen for aortic aneurysms (i.e., subjected to ultrasound to screening aortic aneurysms) to increase life-saving power of cancer immunotherapy.

## Methods

Detailed materials and methods are provided in Supplemental Material.

### Statistical analysis

All data were expressed as mean ± standard error of the mean (SEM). For a comparison of one parameter between the two groups, normality and lognormally tests were performed first. A parametric, unpaired, and two-tailed t-test was performed if the data passed the normality test. A nonparametric, unpaired, and two-tailed test was performed if the data did not pass the normality test. For multiple comparisons of two parameters among multiple groups, a two-way ANOVA with the correction for multiple comparisons by controlling the false discovery rate was performed. For multiple comparisons of three parameters among multiple groups, a three-way ANOVA with the correction for multiple comparisons by controlling the false discovery rate was performed. A two-sided Chi-Square test was performed to compare the incidence of aortic aneurysms between the two groups. A simple linear regression was used to analyze the relationship between two quantitative variables. Significant outliers identified by the outlier calculator (GraphPad) were excluded from the statistical analysis. All statistical analysis was performed by Prism 9 software (GraphPad). A *p*-value or adjusted *p*-value < 0.05 was considered significant unless specified somewhere. A *p*-value of > 0.05 was considered nonsignificant (ns).

### Study approval

All animal procedures were approved by the Institutional Animal Care and Use Committee of the University of Kentucky. All procedures to use human aortic aneurysm specimens for the current study were approved by the Institutional Review Board of the University of Kentucky.

## Author contributions

X.M., S.L., Z.W., and K.J.: contributed to designing research studies, conducting experiments, acquiring data, and analyzing data; T.M, A.S, and A. T. contributed to analyzing RNA-seq data; E.L. and J.C. contributed to providing human aortic aneurysm specimens; M.G. and Z.G. contributed to the conceptualization, supervision, writing, project administration, and funding acquisition.

## Acknowledgments

This work is supported in part by the US NIH Grants HL125228, HL141103, HL142973, and HL164398 (to M.G. and Z.G.), VA Merit Award CX001683 (to E.L. and Z.G.), R01 DC014468 and K18 DC014050 (to T.M.), the Institutional Development Award from the US National Institute of General Medical Sciences of NIH P30GM127211 (to the University of Kentucky Center of Research in Obesity & Cardiovascular Disease), and the Office of the Vice President for Research and an NIH NCI Center Core Support Grant P30 CA177558 (to the University of Kentucky Markey Cancer Center and Flow Cytometry & Immune Monitoring Core Facility).

