## Supplementary Material for "Androgen aggravates aortic aneurysms via suppressing PD-1 in mice"

**This PDF file includes**

Supplementary text (Material and Methods)

Supplemental Figures 1 to 21

Supplemental Tables 1 to 6

Supplemental References

### Material and Methods

#### Animals

Ten-month-old C57BL/6 male and female mice were purchased from the National Institute on Aging (Charles River) or generated by in-house breeding. Mice were housed in the facilities of the Division of Laboratory Animal Resources, University of Kentucky. Mice were fed a chow diet with free access to drinking water. Only 10-13 month-old mice were used in the current study.

#### Administration of mice with aldosterone and high salt (Aldo-salt) to induce aortic aneurysm

Aortic aneurysms were induced by subcutaneous implantation of osmotic minipumps (Alzet model 2004; DURECT) containing aldosterone (200 μg/kg/day) in 10-12-month-old male and female C57BL/6 mice with free access to high salt drinking water (0.9% NaCl plus 0.2% KCl) for four or eight weeks to induce aortic aneurysms as described previously (1-3).

#### *In vivo* aortic ultrasound imaging

The maximal intraluminal diameters of suprarenal abdominal aortas and aortic arches were measured by a high-resolution ultrasound imaging system (Vevo 2100, VisualSonics) in mice one week before (to set up a baseline) and weekly four or eight weeks after Aldo-salt administration. The procedures for measuring the maximal intraluminal diameters of the suprarenal aortas by ultrasound were described previously (1-3). The procedures for measuring the maximal intraluminal diameters of the aortic arch by ultrasound were adapted from Sawada et al. (4). The growth rate of the suprarenal aortic diameters in response to Aldo-salt was calculated from the ultrasound data: growth rate (mm/week) = (the internal diameters of the suprarenal aorta four weeks after Aldo-salt administration - baseline of the suprarenal aorta) / 4 week.

#### *Ex vivo* morphometric analysis of the aorta

The aortas were isolated from mice four or eight weeks after Aldo-salt administration. After dissection to remove fat tissues, the aortas were photographed by the Nikon SMZ2800 stereo microscope with a digital camera. The maximal diameters of the ascending aortas, aortic arch, descending aortas, and suprarenal abdominal aortas were analyzed by Nikon NIS-Element software as described previously (1-3).

**Definition, classification, and quantification of aortic aneurysms**

Aldo-salt-induced AAA and TAA were defined as having at least a 50% increase in the maximal internal (intraluminal) or external diameters in mice administered Aldo-salt compared with the same region of the aorta in control mice or the appearance of evident aortopathy as described previously (1-3). Aldo-salt-induced aortic aneurysms were classified as Type I (the gross appearance of aortic dilation), Type II (at least a 2-time increase in external diameter of the aorta), Type III (a pronounced bulbous form of Type II with a thrombus), and Type IV (aortic rupture) as adapted from Daugherty et al. (5) and described in Supplemental Figure 3. Aldo-salt-induced aortic aneurysms were quantified as the percentage of the incidence of total aortic aneurysms, including abdominal aortic aneurysms (AAA), thoracic aortic aneurysms (TAA), and aortic aneurysm rupture, as described previously (1-3).

**Blood pressure measurement**

Mouse blood pressures were measured by a non-invasive tail-cuff method (Coda 8; Kent Scientific) one week before (to set up a baseline) and three or seven weeks after Aldo-salt administration, as described previously (1-3).

**Orchiectomy**

The procedures for orchiectomy and sham operation were adapted from Henriques et al. (6). Briefly, orchiectomy was performed on mice two weeks before Aldo-salt administration. After anesthetization with 2-3% isoflurane, a single midline incision was made at the caudal abdominal aorta. The skin was separated from the muscle layer, and a similar longitudinal incision was made through the muscle. Both testes can be reached through the same incision. A knot was tied under the testes to reduce bleeding. After the testes and epididymis were excised with scissors, the cut end was cauterized and put back under the muscle. The incision was closed in two layers with a suture. Sham operations were subject to a similar surgical procedure without removing testes.

**Serum testosterone measurement**

Serum was collected by cardiac puncture from orchiectomized and sham-operated mice four weeks after Aldo-salt administration. Serum testosterone levels were measured by a testosterone ELISA kit (Enzo Life Sciences) according to the manufacturer's instructions.

**Urinary and serum sodium measurement and salt retention analysis**

Mice were individually housed in metabolic cages (Tecniplast) to measure 24-h food and water intake and collect 24-h urine samples one week before (to set up a baseline) and three weeks after Aldo-salt administration. Serum was collected four weeks after Aldo-salt administration. The urinary and serum sodium concentrations [Na] were measured by a dual-channel flame photometer (Cole-Parmer) according to the manufacturer's instructions. The 24-h sodium retention was calculated as described by Lee et al. (7) as 24-h sodium intake – 24-h urinary sodium excretion, where 24-h sodium intake = (24 h food weight food x [Na]) + (24 h drinking water x drinking water [Na]), and 24-h urinary sodium excretion = 24 h urine volume x urine [Na].

**Treatment of mice with dihydrotestosterone (DHT), ASC-J9, and anti-PD-1 antibody**

The procedures for administering slow-release DHT pellets to orchiectomized mice were adapted from Henriques et al. (8). DHT pellets (10 mg, 60-day release) were purchased from Innovative Research of America. Mice were orchiectomized for two weeks and then administered with Aldo-salt with or without subcutaneous DHT pellet implantation for four weeks.

The procedures for administering ASC-J9 to mice were adapted from Huang et al. (9). Mice were administered Aldo-salt with or without ASC-J9 (Advanced ChemBlocks) for four weeks. ASC-J9 (25 mg) was dissolved in 120 μl dimethyl sulfoxide (Sigma-Aldrich) and mixed completely until clear yellow color without any uneven color distribution. The mixed solution was then diluted in a pre-warmed (55°C) sesame oil (Sigma-Aldrich) to 1,200 μl, mixed completely, and injected into mice (50 mg ASC-J9/kg mouse body weight, IP, and once a day) as soon as possible.

The procedures for administering anti-mouse PD-1 antibodies to mice were adapted from Koga et al. (10). The rat monoclonal anti-mouse PD-1 antibody (clone 29F.1A12) and rat IgG2a isotype control antibody (clone 1-1) were purchase from Leinco Technologies. Mice were first subjected to orchiectomy for two weeks and then administered Aldo-salt with the anti-PD1 or control antibody (200 μg/mice, IP, and twice a week) for eight weeks.

**Immunocytochemistry (IHC)**

The aorta, spleen, and lymph nodes were isolated from mice ten days, four weeks, and eight weeks after Aldo-salt administration, as indicated in the manuscript and figure legend, respectively. The human aortas with and without aortic aneurysms were obtained from Drs. Eugene Lee (Sacramento Veterans Affairs Medical Center) and John Curci (Vanderbilt University). All tissues were embedded in paraffin and then cut into 5-μm thickness sections.

Aortic elastin breaks were detected by Verhoeff-Van Gieson staining in longitudinal sections of the thoracic aortas and crosssections of the abdominal aorta with an Elastic Stain Kit (Fisher Scientific) according to the manufacturer's instructions. Aortic elastin breaks were quantified by the manual count of elastin breaks under a microscope and expressed as total elastin breaks/section/mice as described previously(2, 3, 11).

The procedure for histological and immunohistological staining was described previously (2, 3, 11). The information on the primary antibodies used in immunohistological staining was described in Supplemental Table 5. A pilot study with different dilutions was conducted to test all primary antibodies to obtain an optimal concentration of antibodies for immunostaining. All antibodies were verified using tissue sections from knockout mice, if available, or a nonspecific IgG to ensure antibody specificity. Immunostaining was quantified by ImageJ (https://imagej.nih.gov/ij/) and expressed as an average percentage of area fractions from five random images per section per mouse as described (12).

**Quantitative analysis of mRNA expression**

The aorta and spleen were isolated from orchiectomized and sham-operated mice ten days after Aldo-salt administration. The procedures for RNA purification, reverse transcription PCR, and real-time PCR were described previously (13-18). The information for real-time PCR primers was described in Supplemental Table 6. All PCR primers were designed to cross introns of chromosomes to eliminate potential genomic DNA contamination and verified by dissociation curve analysis to ensure the specificity of the PCR primers. All mRNA expressions were normalized to the housekeeping gene 36B4 (also known as Rplp0) and quantified by a delta-delta Ct method as described previously (13-18).

**RNA sequence (RNA-seq) and bioinformatics analysis**

The aortas were isolated from three groups of mice (5 mice/group): 1) administered Aldo-salt for one week; 2) orchiectomy followed by one-week Aldo-salt administration; 3) orchiectomy followed by one-week Aldo-salt with DHT administration. After storage and dissection in RNAlater solution, total RNA was purified using an RNeasy mini-kit and then sent to the Genomics Core Laboratory at the University of Kentucky to measure the integrity and purity of purified RNA by Agilent Bioanalyzer

2100 using an RNA nano Chip kit (Agilent Technologies). Purified total RNA samples with RNA integrity number (RIN) > 9 were sent to Novogene for Illumina RNA-seq with a sequencing depth of 20 million reads per sample.

RNA-Seq data in the form of compressed Fastq files from Novogene were uploaded to the Illumina BaseSpace Sequencing Hub (Illumina) and were aligned to the mouse genome (build mm10) using the STAR Aligner v2.5.2a. After alignment, differentially abundant mRNAs were identified using DESeq2 (19). The mRNAs with $p < 0.01$ were considered differentially abundant. Log2 (fold change) and -log10 ($p$-value) were used for the visualization of differences in transcript abundance in the form of volcano plots using Prism 9 (GraphPad). Venn diagrams and heatmaps were generated by TBtools, a Toolkit for Biologists (20). The 180 mRNAs upregulated by orchiectomy but downregulated DHT and 150 mRNAs downregulated by orchiectomy but upregulated by DHT were subjected to comprehensive pathway enrichment analyses via Enrichr using the Bioplanet 2019 database (https://maayanlab.cloud/Enrichr/) (21). The pathways with adjusted $p < 0.01$ (Benjamini-Hochberg method) were considered significant.

**Flow cytometry analysis of immune cells**

The procedures for flow cytometry analysis of immune cells in the mouse aorta, spleen, and blood were adapted from Galkina et al. (22) and Melak et al. (23). After 4-week Aldo-salt administration, the aortas were harvested from splenectomized or sham-operated male mice perfused with PBS containing 2% heparin. Isolated aortas were dissected to remove fatty tissue while keeping the adventitia intact. The aortas were cut into small pieces and then digested with 450 U/mL collagenase type I, 125 U/mL collagenase type XI, 60 U/mL hyaluronidase type I-s, and 60 U/mL DNase-I in DMEM media. All enzymes except for collagenase type I, which was purchased from Worthington Biochemicals, were obtained from Sigma-Aldrich and prepated in Dulbecco′s Modified Eagle′s Medium (DMEM; Fisher Scientific) at 37 $^0$C for 1 hour, with gentle shaking and vortex. After 10-day

Aldo-salt administration, the spleen, blood, and lateral aortic lymph nodes were harvested from orchiectomized and sham-operated mice perfused with PBS containing 2% heparin. A small piece of the spleen was removed from the whole spleen for flow cytometry. The small piece of the spleen and the whole spleen were weighed to calculate the total immune cells in the spleen. A single cell suspension was obtained by using syringe plunges to mash the small piece of the spleen, digested aortas, and pooled lymph nodes through a 70-μm strainer (Fisher Scientific). Blood and splenic cells were incubated with an RBC lysis buffer (eBioscience) to remove erythrocytes.

Cells were first mixed with an anti-mouse CD16/32 antibody (Biolegend) to block the nonspecific binding of the immunoglobulin to Fc receptors and then incubated with fluorescence-conjugated primary antibodies (Supplemental Table 5) specific for leukocytes ($CD45^+$), T cells ($CD3^+$), B cells ($CD19^+$), macrophages ($F4/80^+$), neutrophils ($Ly6G^+$), and PD-$1^+$ immune cells (i.e., $CD45^+CD3^+$PD-$1^+$). After washing, cells were fixed by a stabilizing fixative solution (BD Biosciences), mixed with Precision Count Beads (Biolegend; to obtain absolute counts of cells by flow cytometry), and then submitted to the University of Kentucky Flow Cytometry and Immune Monitoring Core Facility for flow cytometry analysis with BD FADSymphony 2.0. Cells from the spleen, lymph nodes, and blood were also stained with all the fluorophores minus one of them (FMO) to identify gating boundaries and ensure the specificity of antibodies. Data were analyzed using FlowJo$^{TM}$ v10.

**Western blot**

The spleens were isolated from orchiectomized and sham-operated mice ten days after Aldo-salt administration. The procedures for using the trichloroacetic acid (TCA) method to prepare protein samples from the spleen for Western blot analysis were described previously (15-17). The anti-mouse PD-1, CD3ε, CD19, GAPDH, and anti-human AR antibodies were described in Supplemental Table 5. Western blots were quantified by ImageJ as described (15-17).

**AR ChIP assay**

A splenic cell suspension was prepared by mashing one freshly-isolated mouse spleen through a 70-μm strainer for the AR ChIP assay. Approximately 4 X $10^6$ splenic cells were used for one ChIP assay. The AR ChIP assay was performed using SimpleChIP® Enzymatic Chromatin IP Kit (Magnetic Beads; Cell Signaling) according to the manufacturer's instructions. Two ChIP-grade anti-mouse AR antibodies with different epitopes (Supplemental Table 5) were used in the AR ChIP assay to pull down chromatins containing the PD-1 promoter. A normal rabbit IgG (Cell Signaling) was included in the AR ChIP assay as a control to ensure the specificity of AR immunoprecipitation. Two sets of ChIP-PCR primers were designed to amplify ARE4-5 and ARE 6 in the mouse PD-1 promoter (Figure 7J) and described in Supplemental Figure 6. The binding of AR to the mouse PD-1 promoter was quantified by ImageJ and expressed as the percentage of genomic DNA input as described (17, 18).

**AR promoter assay**

A 488-bp mouse PD-1 promoter (-4,444 to -3,956 bp relative to TSS) containing putative ARE6-10 (Figure 7J) was synthesized by Integrated DNA Technology (IDT) and then subcloned into the pGL3-basic firefly luciferase reporter vectors (Promega) in Kpn I and Hind III restriction enzyme sites. The pGL3-basic-PD-1 promoter construct (0.75 μg) was co-transfected with a pcDNA Flag-M4-human AR construct (3 μg), provided by Dr. Steven Balk (Addgene plasmid # 171240) (24), with 1:4 molecular ratio of PD-1 promoter to AR, into AD-HEK 293 cells (Stratagene) by LipoFexin (Lamda Biotech) according to the manufacturer's instructions. A pcDNA3.1 vector (Invitrogen; 2.25 μg) was added to the cells with the pGL3-basic vector or the PD-1 promoter only to maintain the same ratio of DNA to LipoFexin. A Renilla luciferase control reporter vector (pRL-TK control vector; 10 ng; Promega) was included in all samples to control transfection efficiency. After 4-hour transfection, the cell culture medium was changed to a DMEM supplemented with 10% charcoal dextran-stripped fetal bovine serum (VWR) to minimize potential endogenous hormone effects. DHT (100 nM) was added into the cells with the pcDNA Flag-M4-AR construct to activate AR after 24-h transfection. Cells were harvested after 48-h transfection. The PD-1 promoter activity was analyzed by a modified dual luciferase enzyme assay as described previously (16-18).

**Statistical analysis**

All data were expressed as mean ± standard error of the mean (SEM). For a comparison of one parameter between the two groups, normality and lognormally tests were performed first. A parametric, unpaired, and two-tailed t-test was performed if the data passed the normality test. A nonparametric, unpaired, and two-tailed test was performed if the data did not pass the normality test. For multiple comparisons of two parameters among multiple groups, a two-way ANOVA with the correction for multiple comparisons by controlling the false discovery rate was performed. For multiple comparisons of three parameters among multiple groups, a three-way ANOVA with the correction for multiple comparisons by controlling the false discovery rate was performed. A two-sided Chi-Square test was performed to compare the incidence of aortic aneurysms between the two groups. A simple linear regression was used to analyze the relationship between two quantitative variables. A *p*-value or adjusted *p*-value $< 0.05$ was considered significant unless specified somewhere. A *p*-value of $> 0.05$ was considered nonsignificant (ns). All statistical analysis was performed by Prism 9 software (GraphPad).

**Study approval**

All procedures to use mice for the current study were approved by the Institutional Animal Care and Use Committee of the University of Kentucky. All procedures to use human aortic aneurysm specimens for the current study were approved by the Institutional Review Board of the University of Kentucky.

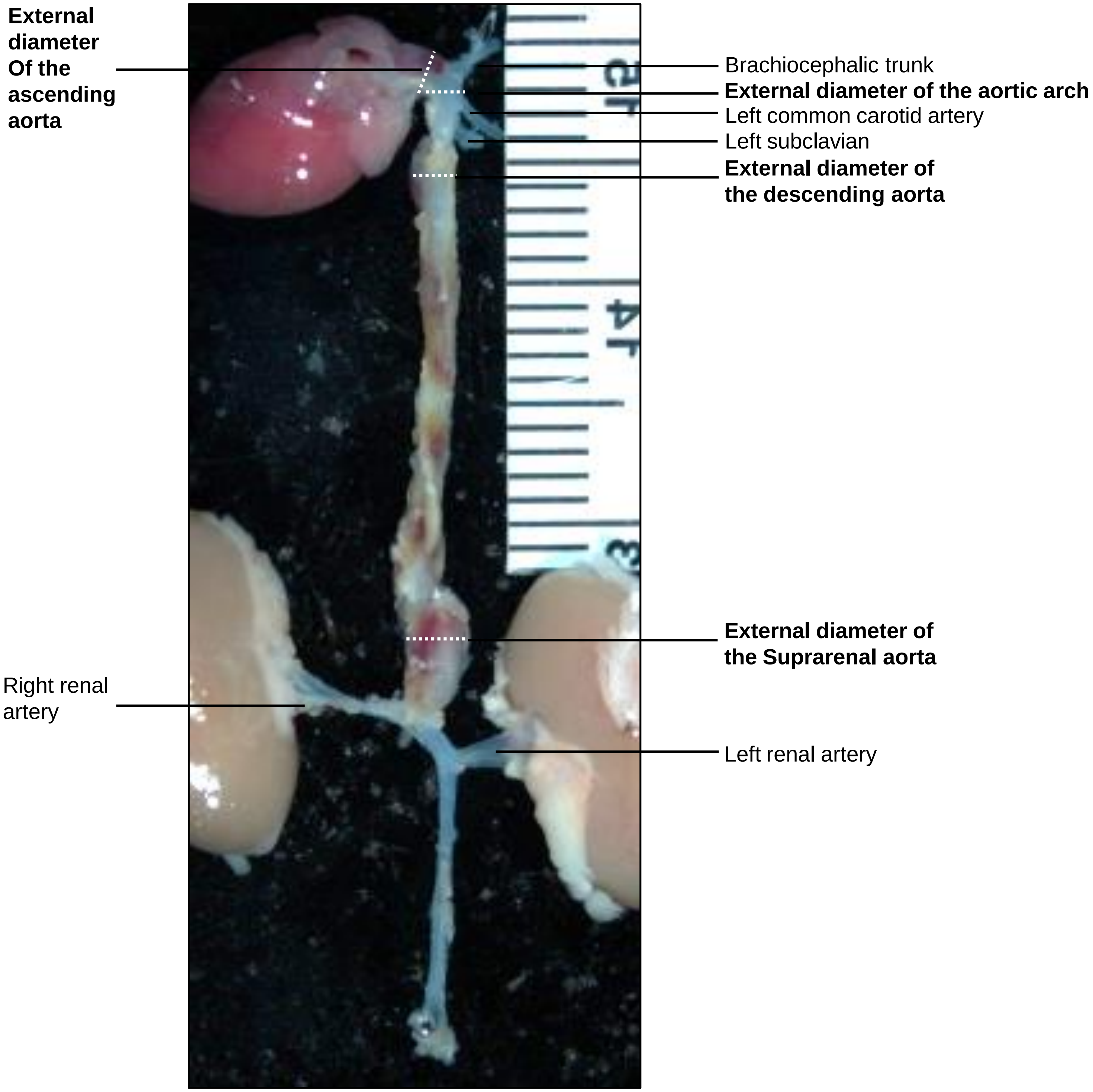


**Supplemental Figure 1. *Ex Vivo* measurement of maximal external diameters of the ascending aorta, aortic arch, descending aorta, and suprarenal aorta by microscopy.** The brachiocephalic trunk, left common carotid artery, left subclavian, right renal artery, and left renal artery are indicated. The dotted dash lines labeled in the white show the maximal external diameters of ascending aorta, aortic arch, descending aorta, and suprarenal aorta, measured in the current study.

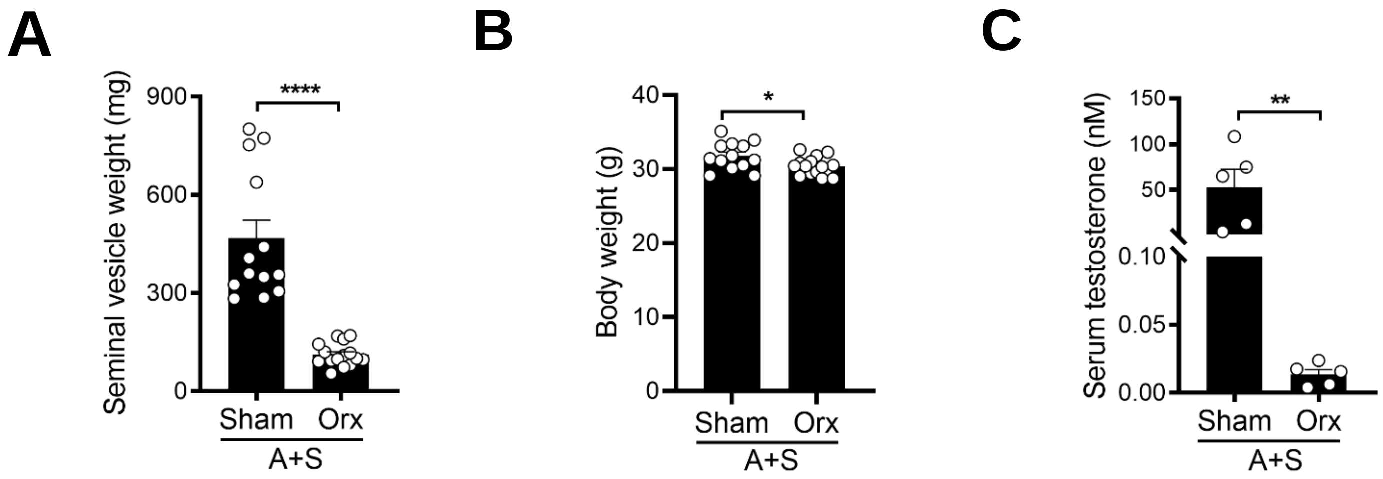


**Supplemental Figure 2. Gonadal androgen deprivation decreases seminal vesicle weight, body weight, and serum testosterone in mice administered aldosterone and high salt (Aldo-salt).** The seminal vesicle weight (**A**), body weight (**B**), and serum testosterone level (**C**) were measured in 10-month-old male C57BL/6 mice with orchiectomy (orx) or sham operation four weeks after Aldo-sat (A+S) administration. The data were expressed as mean ± standard error of the mean (SEM) and analyzed by a two-tailed unpaired T-test. **, $p < 0.05$; **, $p < 0.01$; ****, $P < 0.0001$.*

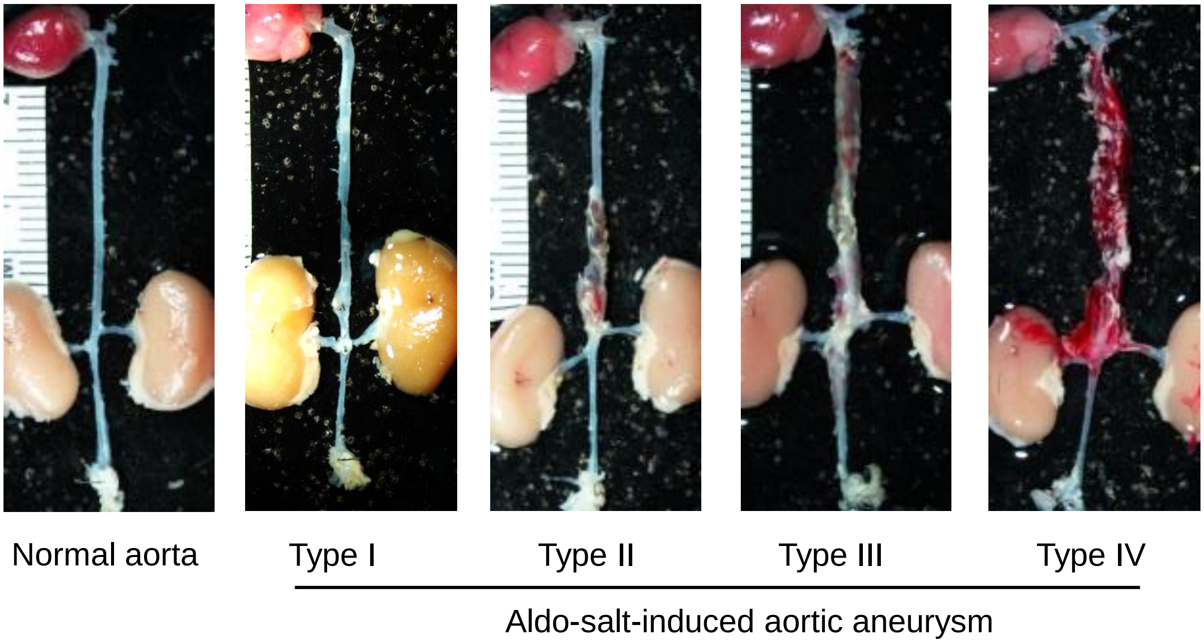


**Supplemental Figure 3. Classification of Aldo-salt-induced aortic aneurysms in 10-month-old or older C57BL/6 mice.** Aldo-salt-induced aortic aneurysms were classed as Type I (a gross appearance of the abdominal or thoracic aortic dilation compared with the normal aorta), Type II (at least two times the normal external diameter of the abdominal or thoracic aorta, frequently containing a thrombus), Type III (a pronounced bulbous form of type II that includes a thrombus in both the abdominal and thoracic aorta), and Type IV (aortic aneurysms with rupture).

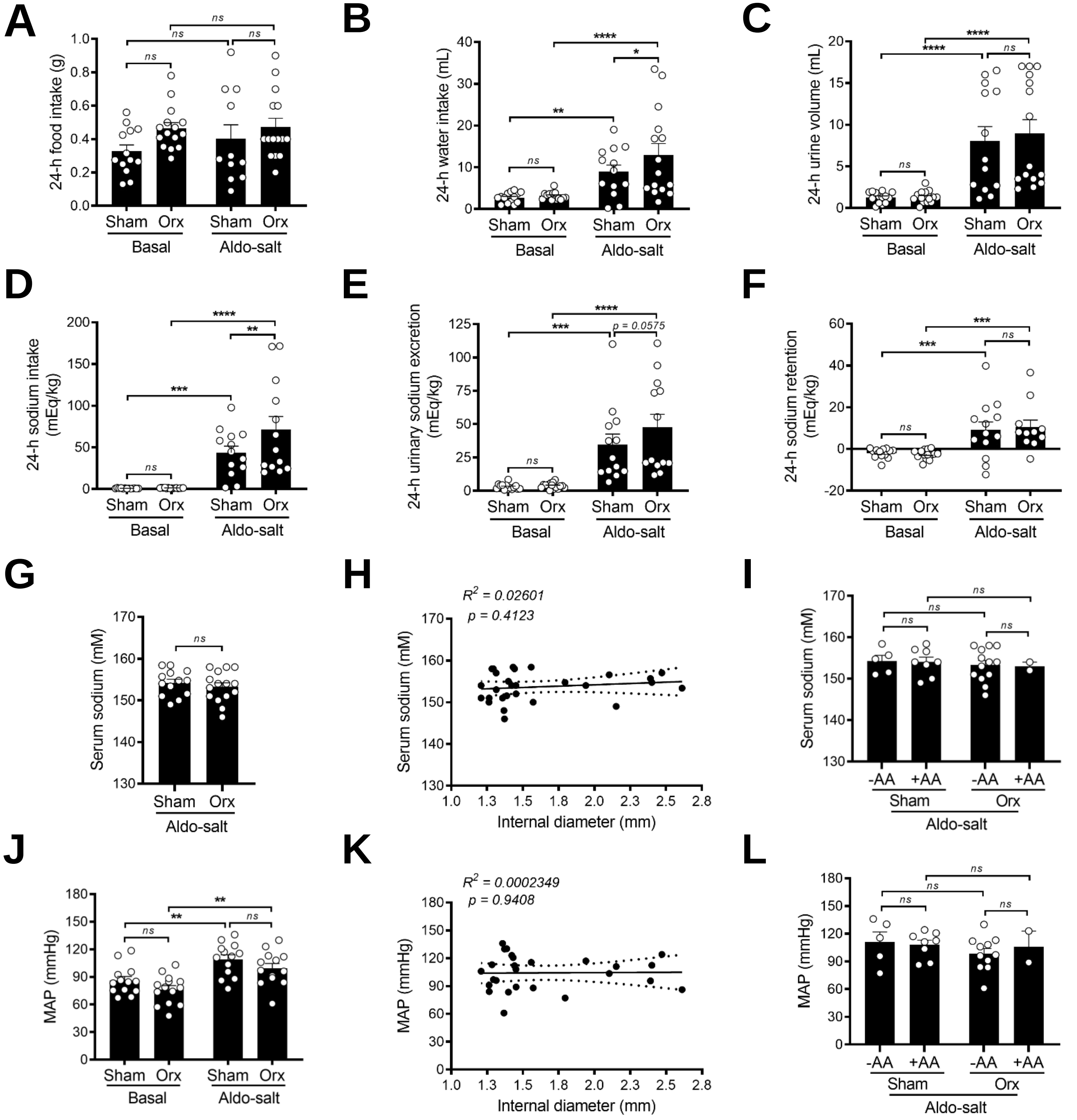


**Supplemental Figure 4. Gonadal androgen deprivation does not affect Aldo-salt-induced sodium retention and hypertension.** (**A**–**F**) 24-h food intake (A), water intake (B), urine volume (C), sodium intake (D), urinary sodium excretion (E), and sodium retention (F) were measured in 10-month-old male C57BL/6J mice with orchiectomy (orx) or sham operation one week before (basal) and three weeks after Aldo-salt administration. (**G**) Serum sodium levels between mice with orx and sham-operation four weeks after Aldo-salt administration. (**H**) Correlation analysis of the internal diameter of the suprarenal aorta and serum sodium levels in orx and sham-operated mice four weeks after Aldo-salt administration. (**I**) Serum sodium levels in orx or sham-operated mice with (+) and without (-) aortic aneurysms (AA). (**J**) Mean arterial pressure (MAP) in mice one week before and three weeks after Aldo-salt administration. (**K**) Correlation analysis of the internal diameter of the suprarenal aorta and MAP in mice with orx or sham-operation three weeks after Aldo-salt administration. (**L**) MAP in orx and sham-operated mice with (+) and without (-) aortic aneurysm (AA). Data were expressed as mean ± SEM and analyzed by two-way ANOVA with multiple comparison tests (A–F, I, J, and L), two-tailed unpaired *t*-test (G), and simple linear regression analysis (H and K). **, p < 0.05; **, p < 0.01; ***, P < 0.001; ****, P < 0.0001; ns, not significant.*

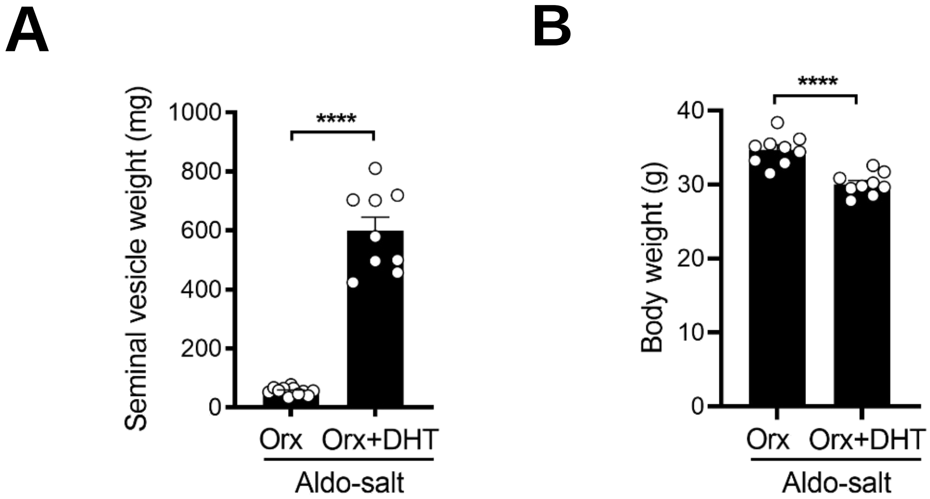


**Supplemental Figure 5. Effects of exogenous dihydrotestosterone (DHT) on seminal vesicle weight and body weight in orchiectomized mice administered Aldo-salt.** The seminal vesicle weight (**A**) and body weight (**B**) were measured in 10-month-old male C57BL/6 mice with orchiectomy (orx) four weeks after Aldo-salt with or without dihydrotestosterone (DHT) administration. The data were expressed as mean ± SEM and analyzed by a two-tailed unpaired *t*-test. ****, $p < 0.0001$.

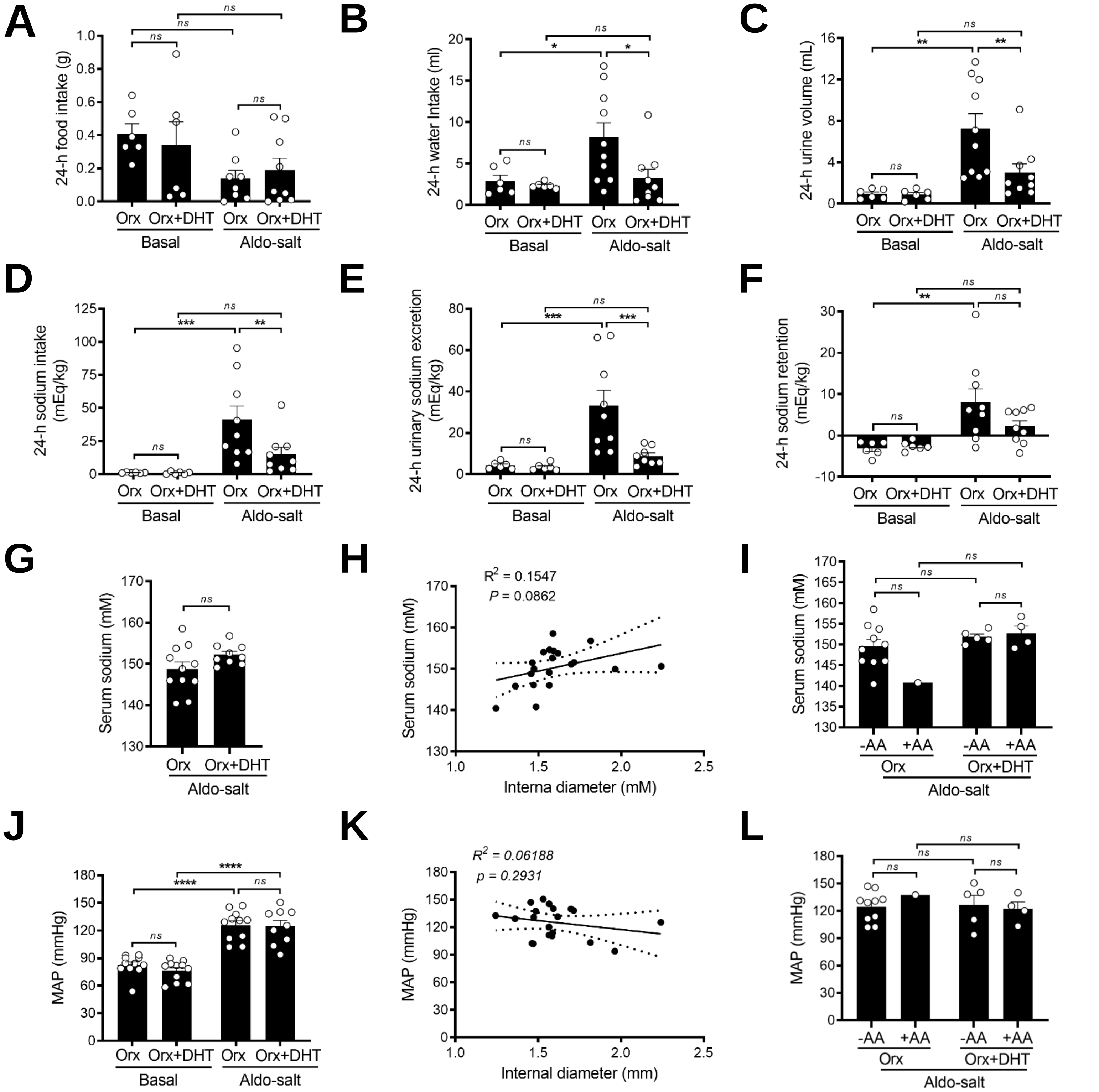


**Supplemental Figure 6. Exogenous dihydrotestosterone administration to orchiectomized mice does not affect Aldo-salt-induced sodium retention and hypertension.** (**A**–**F**) 24-h food intake (A), water intake (B), urine volume (C), sodium intake (D), urinary sodium excretion (E), and sodium retention (F) were determined in 10-month-old C57BL/6J mice with orchiectomy (orx) one week before (basal) and three weeks after Aldo-alt with or without DHT administration. (**G**) Serum sodium levels in orchiectomized mice four weeks after Aldo-salt with or without DHT administration. (**H**) Correlation analysis of the internal diameter of the suprarenal aorta and serum sodium levels in orchiectomized mice four weeks after Aldo-salt with or without DHT administration. (**I**) Serum sodium levels between orx or orx+DHT mice with (+) or without (-) aortic aneurysms (AA). (**J**) MAP in orx mice one week before and three weeks after Aldo-salt with or without DHT administration. (**K**) Correlation analysis of the internal diameter of the suprarenal aorta and MAP in orchiectomized mice three weeks after Aldo-salt with or without DHT administration. (**L**) MAP in orx and orx+DHT mice with and without AA. Data were expressed as mean ± SEM and analyzed by two-way ANOVA for multiple comparisons (A–F, **I**, J, and L), two-tailed unpaired T-test (G), and simple linear regression analysis (H and K). **, p < 0.05; **, p < 0.01; ***, p < 0.001; ****, P < 0.0001; ns, not significant.*

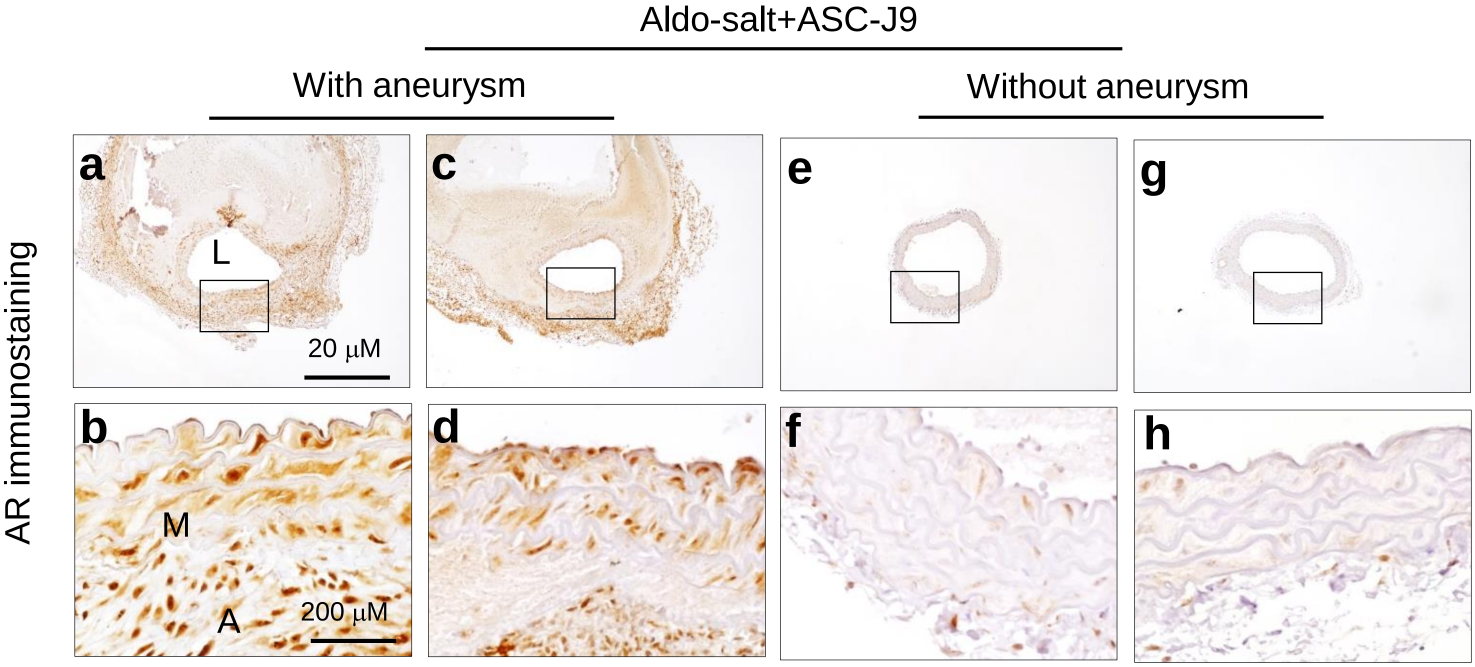


**Supplemental Figure 7. Representative immunostaining of androgen receptors (AR) in the suprarenal aortas in mice administered Aldo-salt.** The suprarenal aortas were isolated from 10-month-old male C57BL/6J mice four weeks after Aldo-salt with ASC-J9 administration and subjected to AR immunostaining. (**a−d**) AR immunostaining of the suprarenal aortas in mice administered Aldo-salt and ASC-J9 but developed AAA. (**e−h**) AR immunostaining of the suprarenal aortas in mice administered Aldo-salt and ASC-J9 but did not develop AAA. The area of rectangles in photographs a, c, e, and g indicated the magnified areas in photographs b, d, f, and h, respectively. L, lumen. M, media. A, adventitia.

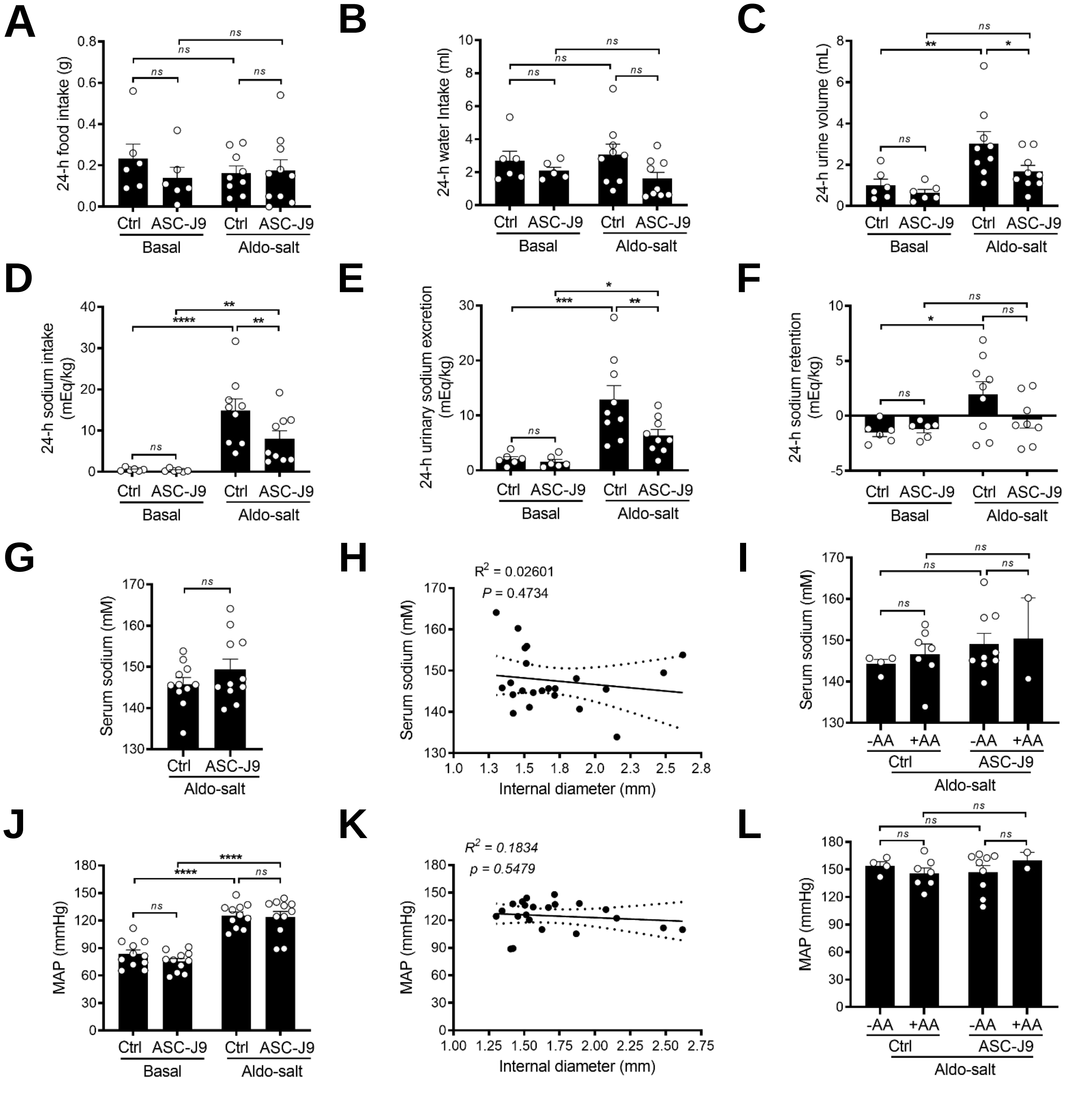


**Supplemental Figure 8. Downregulation of androgen receptors with ASC-J9 has little effect on Aldo-salt-induced sodium retention and hypertension.** (**A**–**F**) 24-h food intake (A), water intake (B), urine volume (C), sodium intake (D), urinary sodium excretion (E), and sodium retention (F) were determined in 10-month-old male C57BL/6J mice one week before (basal) and three weeks after Aldo-salt with or without (Ctrl) ASC-J9 administration. (**G**) Serum sodium levels in mice four weeks after Aldo-salt with or without ASC-J9 administration. (**H**) Correlation analysis of the internal diameter of the suprarenal aorta and serum sodium levels in mice four weeks after Aldo-salt with or without ASC-J9 administration. (**I**) serum sodium levels in mice administered Aldo-salt with or without ASC-J9 with (+) and without (-) aortic aneurysms (AA). (**J**) MAP was measured one week before and three weeks after Aldo-salt with or without ASC-J9 administration. (**K**) Correlation analysis of the internal diameter of the suprarenal aorta and MAP in mice three weeks after Aldo-salt with or without ASC-J9 administration. (**L**) MAP in mice administered Aldo-salt with or without ASC-J9 and with (+) or without (-) AA. Data were expressed as mean ± SEM and analyzed by two-way ANOVA for multiple comparisons (A–F, I, J, and L), two-tailed unpaired *t*-test (G), and simple linear regression analysis (H and K). ***, $p < 0.01$; ****, $P < 0.0001$; ns, not significant.*

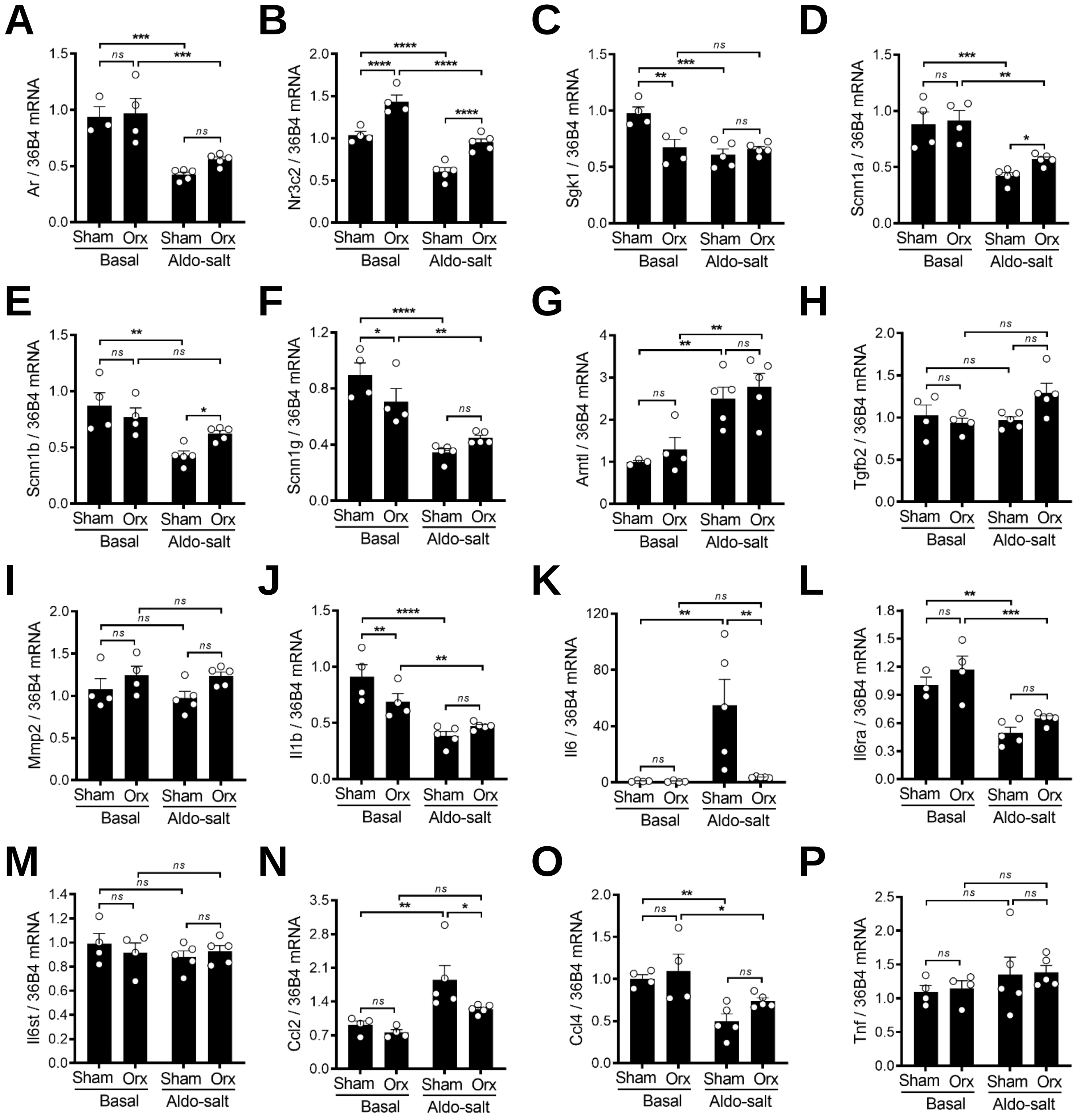


**Supplemental Figure 9. Effects of gonadal androgen deprivation on mRNA expression in the aorta in mice administered Aldo-salt.** The mRNA expressions in the whole aortas were determined by real-time PCR and normalized to 36B4 in 10-month-old male C57BL/6 mice with orchiectomy (orx) or sham operation ten days after Aldo-salt administration. (**A**) Androgen receptor (Ar). (**B**) Mineralocorticoid receptor (Nr3c2; also known as MR). (**C**) Serum and glucocorticoid-regulated Kinase 1 (Sgk1). (**D**) Sodium channel epithelial 1 subunit alpha (Scnn1a; also known as ENaCα). (**E**) Scnn1b (also known as Enacβ). (**F**) Scnn1g (also known as ENaCγ). (**G**) Aryl hydrocarbon receptor nuclear translocator-like (Arntl; also known as Bmal1). (**H**) Transforming growth factor beta 2 (Tgfb2; also known as TGFβ2). (**I**) Matrix metallopeptidase 2 (Mmp2). (**J**) Interleukin 1 beta (Il1; also known as IL-1β). (**K**) Interleukin 6 (Il6; also known as IL-6). (**L**) Interleukin 6 receptor alpha (Il6ra; also known as IL-6Rα). (**M**) Interleukin 6 signal transducer (Il6st; also known as IL-6Rβ or gp130). (**N**) C-C motif chemokine ligand 2 (Ccl2; also known as MCP-1). (**O**) C-C motif chemokine ligand 4 (Ccl4). (**P**) Tumor necrosis factor (Tnf; also known as TNF-α). Data were expressed as mean ± SEM and analyzed by two-way ANOVA for multiple comparisons. **, p < 0.05; **, p < 0.01; ***, p < 0.001; ****, P < 0.0001; ns, not significant.*

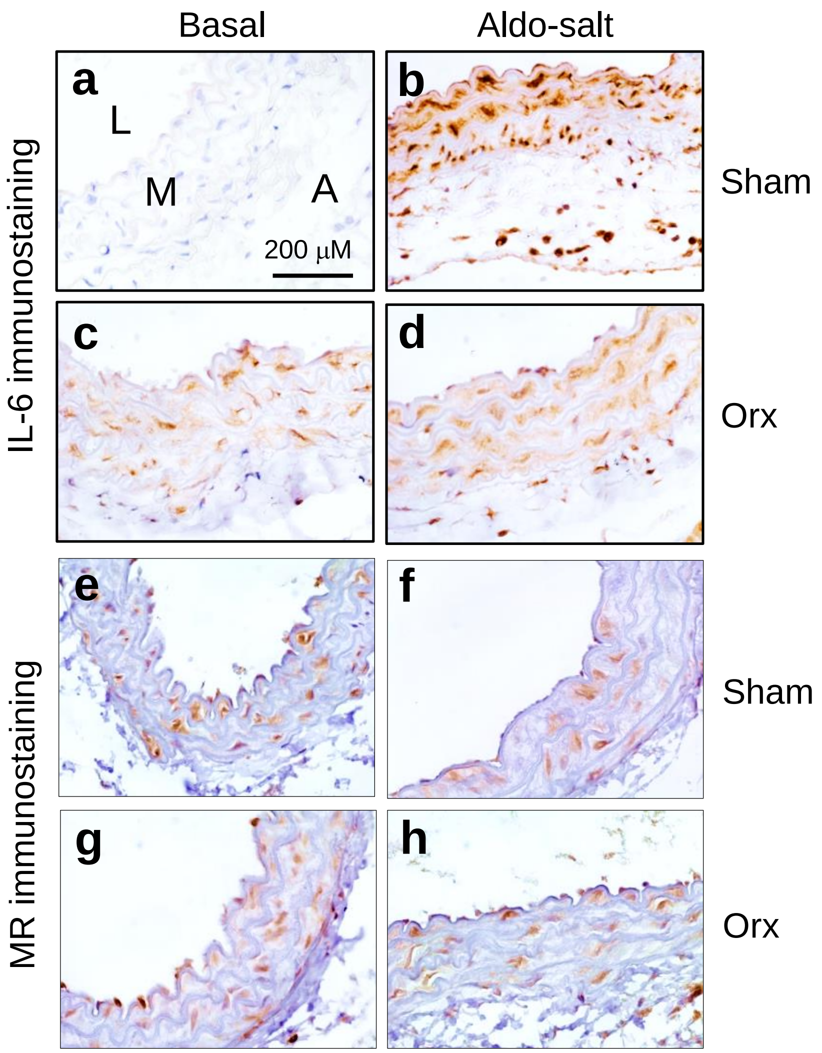


**Supplemental Figure 10. Gonadal androgen deprivation affects the IL-6 but not MR protein expression in the suprarenal aorta before and after Aldo-salt administration.** Representative **i**mmunostaining of interleukin 6 (IL-6; a−d) and mineralocorticoid receptor (MR; e−h) in the cross-sections of the suprarenal aortas in 10-month-old C57BL/6J mice with orchiectomy or sham operation seven days before (basal) and ten days after Aldo-salt administration (3 mice/group). L, lumen. M, media. A, adventitia.

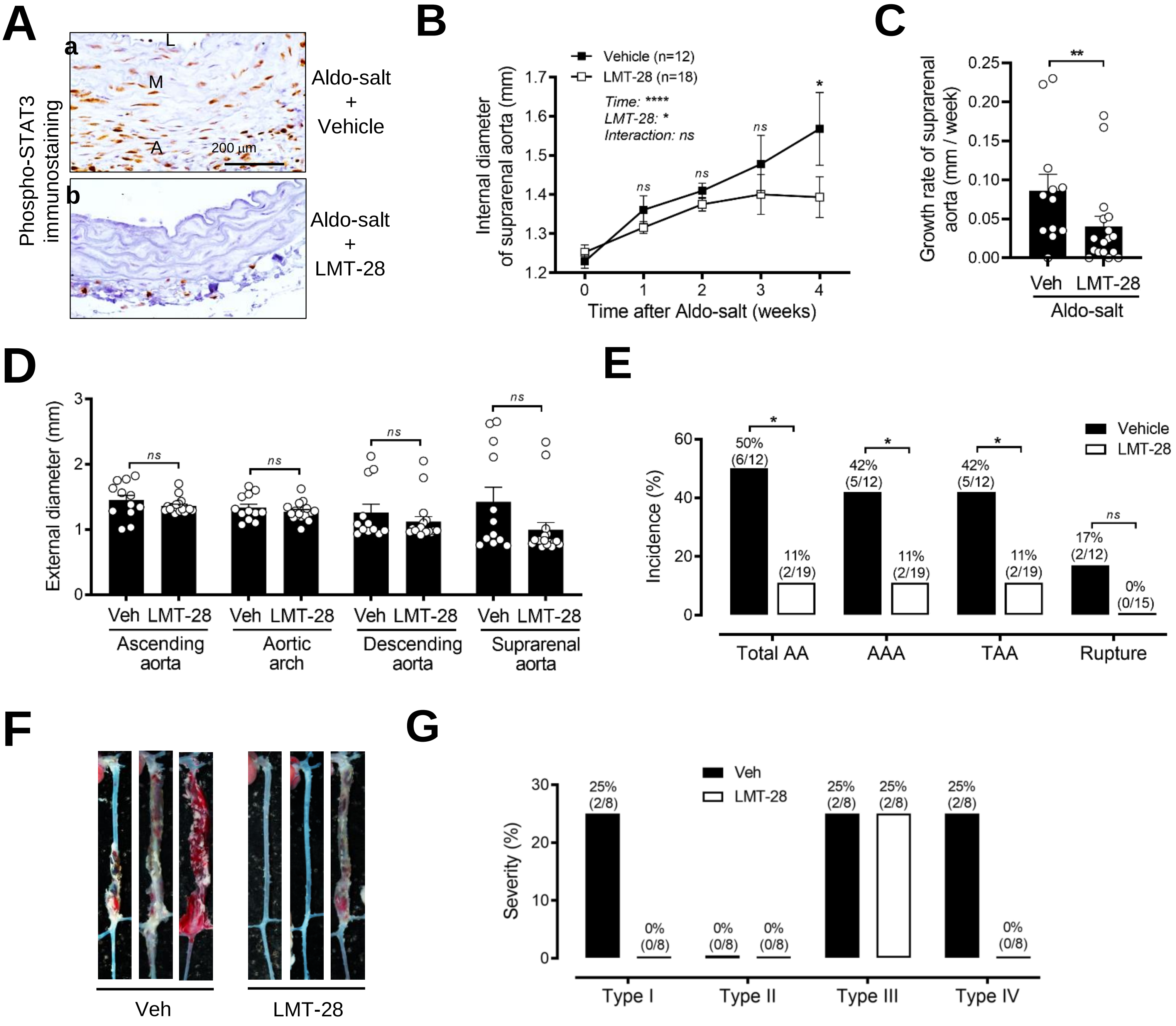


**Supplemental Figure 11. Inhibition of IL-6 signaling by LMT-28 ameliorates Aldo-salt-induced aortic aneurysms.** (**A**) Representative immunostaining of STAT3 phosphorylation at Tyr705 in the abdominal aorta in 10-month-old male C57BL/6J mice four weeks after Aldo-salt with LTM-28 or vehicle (carboxy methyl cellulose) administration. (**B** and **C**) *In vivo* quantification of the maximal intraluminal diameters (B) and growth rate (C) of the suprarenal aortas by ultrasound one week before (time 0) and weekly after Aldo-salt with LMT-28 or vehicle administration. (**D**) ) *Ex vivo* measurement of the maximal external diameters of the ascending aorta, aortic arch, descending aorta, and suprarenal aorta in isolated aortas four weeks after Aldo-salt with LMT-28 or vehicle administration. (**E**) The incidence of total aortic aneurysms (AA), AAA, TAA, and aortic aneurysm rupture. The percentages of the incidences (number of mice with total AA, AAA, TAA, and rupture / total number of mice) were indicated on the top of each bar. (**F**) Representative photographs of the aortas in mice with or without aortic aneurysms. (**G**) The percentages of severity (number of mice with Type I, II, III, and IV aortic aneurysms / total number of mice with aortic aneurysms) were indicated on the top of each bar. The data were expressed as mean ± SEM and analyzed by two-way ANOVA with multiple comparisons test (B), two-tailed unpaired *t*-test (C and D), and two-sided Chi-square test (E). **, p < 0.05; **, p < 0.01; ****, P < 0.0001; ns, not significant.*

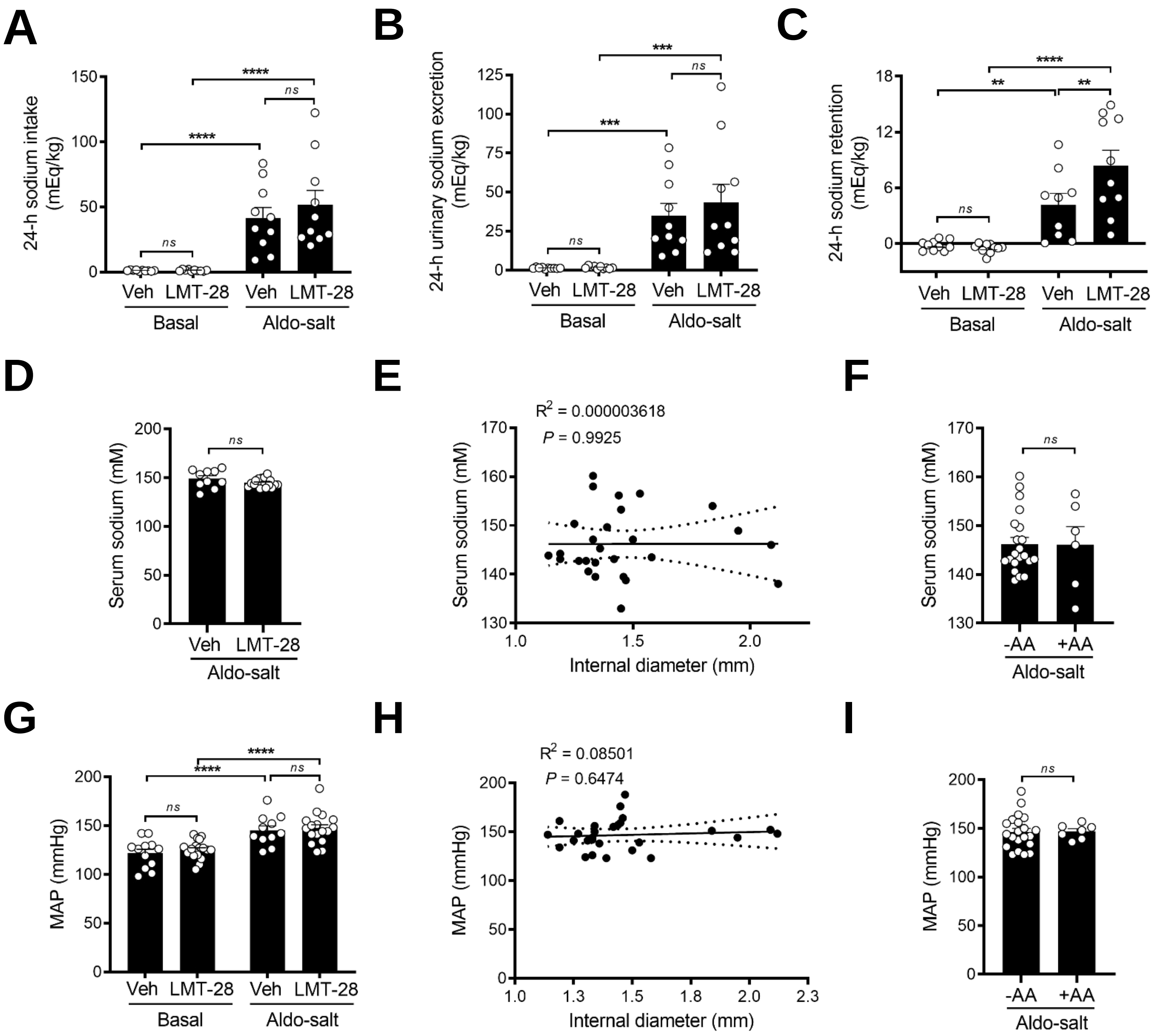


**Supplemental Figure 12. LMT-28 has no effect on Aldo-salt-induced renal sodium retention and hypertension.** (**A**–**C**) 24-h sodium intake (A), urinary sodium excretion (B), and sodium retention (C) were determined in 10-month-old male C57BL/6J mice one week before (basal) and three weeks after Aldo-salt with LMT-28 or vehicle (Veh) administration. (**D**) Serum sodium levels in mice four weeks after Aldo-salt with LMT-28 or vehicle administration. (**E**) ) Correlation analysis of the internal diameter of the suprarenal aorta and serum sodium levels in mice four weeks after Aldo-salt with LMT-28 or vehicle administration. (**F**) Serum sodium levels in mice administered with LMT-28 or vehicle with and without aortic aneurysms (AA). (**G**) Mean arterial pressure (MAP) was measured one week before (basal) and three weeks after Aldo-salt with LMT-28 or vehicle administration. (**H**) Correlation analysis of the MAP and internal diameter of the suprarenal aorta four weeks after Aldo-salt with LMT-28 or vehicle administration. (**I**) MAP in mice administered Aldo-salt with LMT-28 or vehicle with (+) and without (-) aortic aneurysm (AA). Data were expressed as mean ± SEM and analyzed by two-way ANOVA for multiple comparisons (A-C and G), two-tailed unpaired *t*-test (D, F, and I), and simple linear regression analysis (E and H). **, p < 0.05; **, p < 0.01; ***, p < 0.001; ****, P < 0.0001; ns, not significant.*

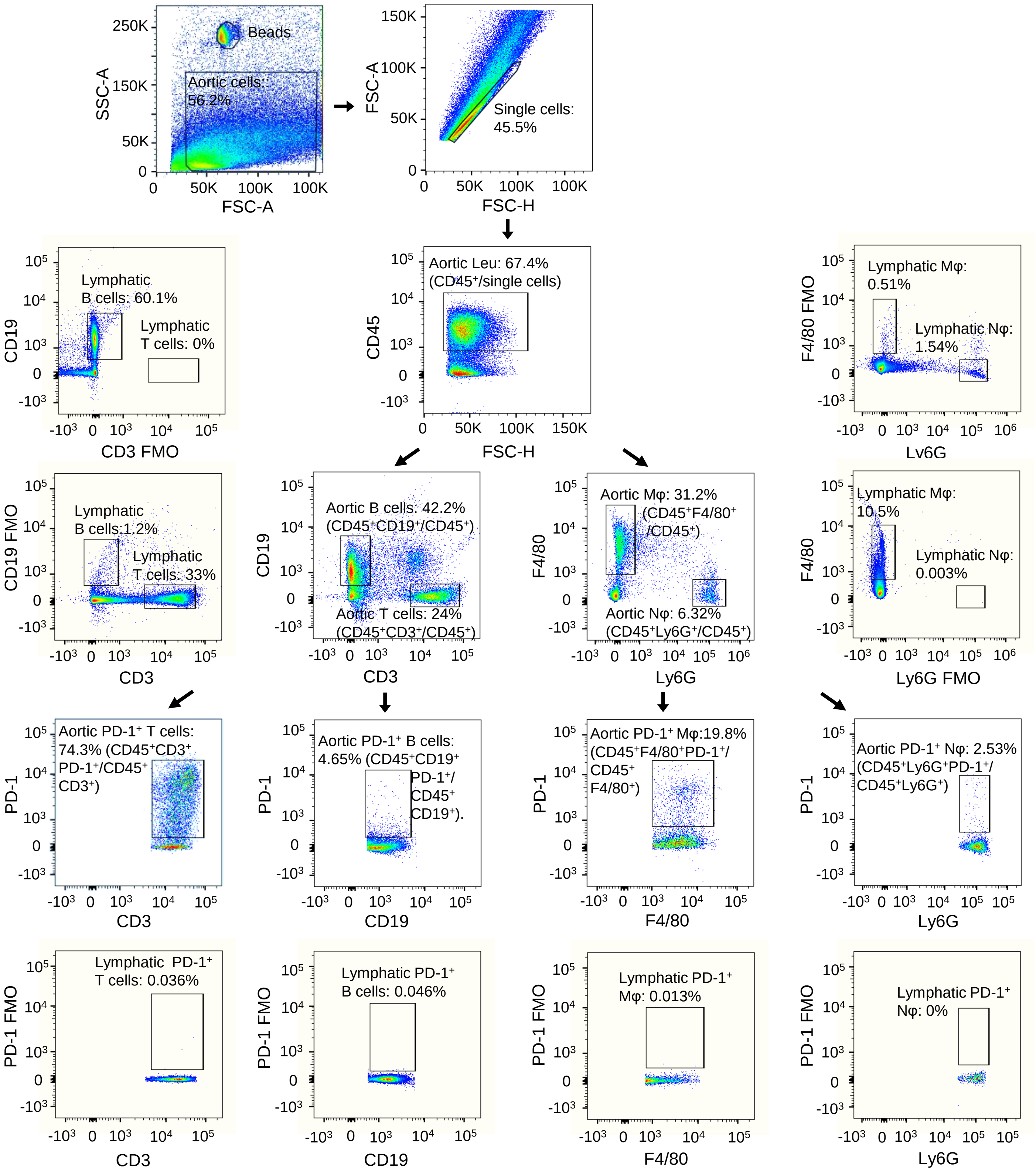


**Supplemental Figure 13. Gating strategy to identify PD-1-positive T cells, B cells, macrophages, and neutrophils in the mouse aorta.** Single cells were prepared from the mouse aortas and lymph notes and gated in by a forward scatter height (FSC-H) *vs.* forward scatter area (FSC-A) pseudocolor plot. Single cells were gated on CD45$^+$ to select leukocytes (Leu) and then gated on CD3$^+$, CD19$^+$, F4/80$^+$, and Ly6G$^+$ to identify T cells (CD45$^+$CD3$^+$), B cells (CD45$^+$CD19$^+$), macrophages (Mφ; CD45$^+$F4/80$^+$), and neutrophils (Nφ; CD45$^+$Ly6G$^+$), respectively. The cells were then gated on PD-1$^+$ to identify PD-1$^+$ T cells (CD45$^+$CD3$^+$PD-1$^+$), B cells (CD45$^+$CD19$^+$PD-1$^+$), macrophages (CD45$^+$F4/80$^+$PD-1$^+$), and neutrophils (CD45$^+$Ly6G$^+$PD-1$^+$), respectively. The single cells from lymph nodes were also analyzed by flow cytometry with fluorescence minus one (FMO) (labeled with a yellow background in pseudocolor plots) to identify gating boundaries and ensure the specificity of antibodies.

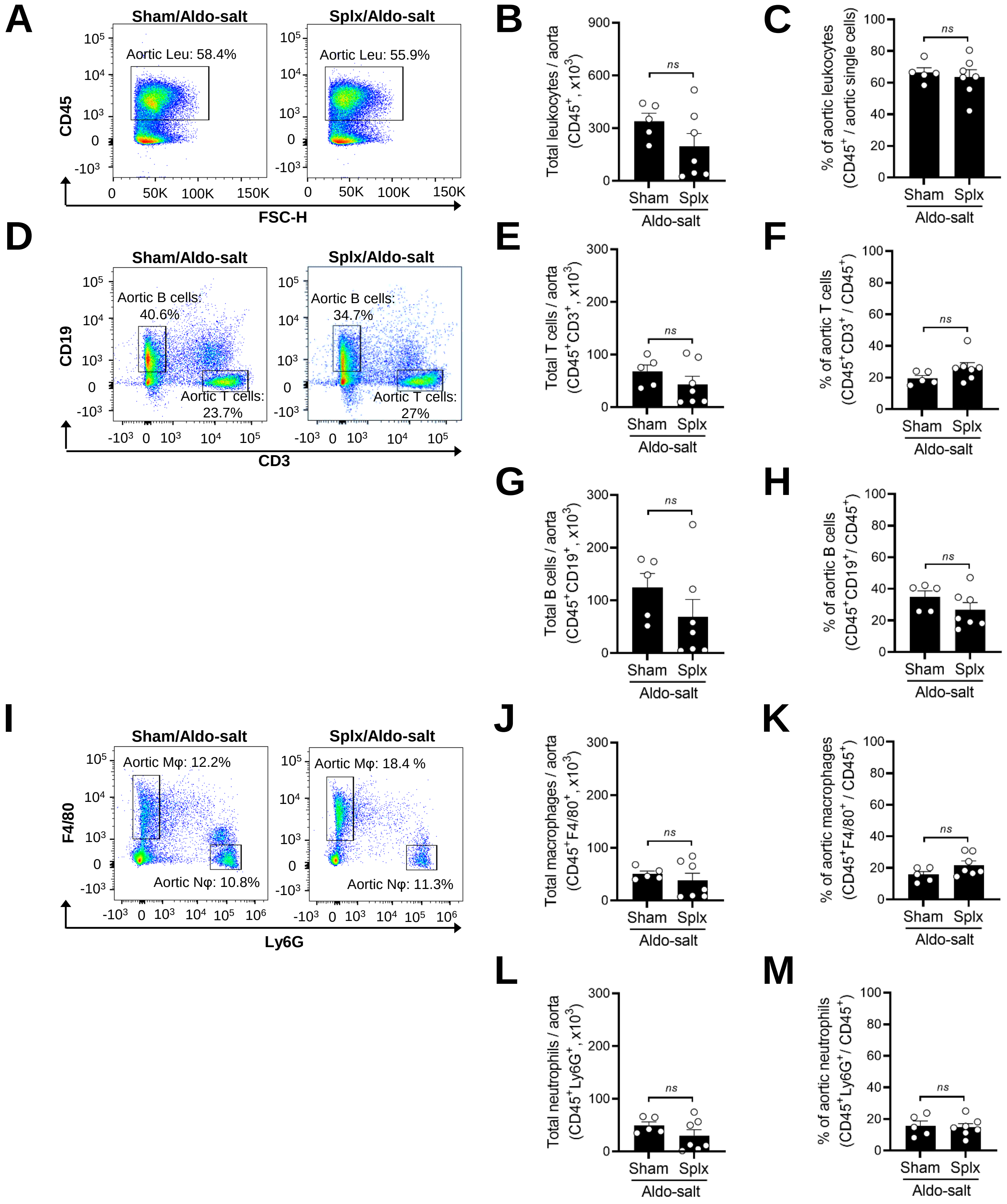


**Supplemental Figure 14. Splenectomy does not alter leukocytes, T cells, B cells, macrophages, and neutrophils in the aorta in mice administered Aldo-salt.** Representative pseudocolor plots and quantitative data of flow cytometry analysis of the total and percentage of leukocytes (Leu; $CD45^+$; **A−C**), T cells ($CD45^+CD3^+$; **D−F**), B cells ($CD45^+CD19^+$; **D**, **G**, and **H**), macrophages (Mφ; $CD45^+F4/80^+$; **I−K**), and neutrophils (Nφ; $CD45^+Ly6G^+$; **I**, **L**, and **M**) in the whole aortas in 11-13-month-old male C57BL/6J mice with splenectomy (splx) or sham operation four weeks after Aldo-salt administration. One symbol in the figures represents the cells of one mouse aorta. The data were expressed as mean ± SEM and analyzed by a two-tailed unpaired *t*-test. *Ns, not significant.*

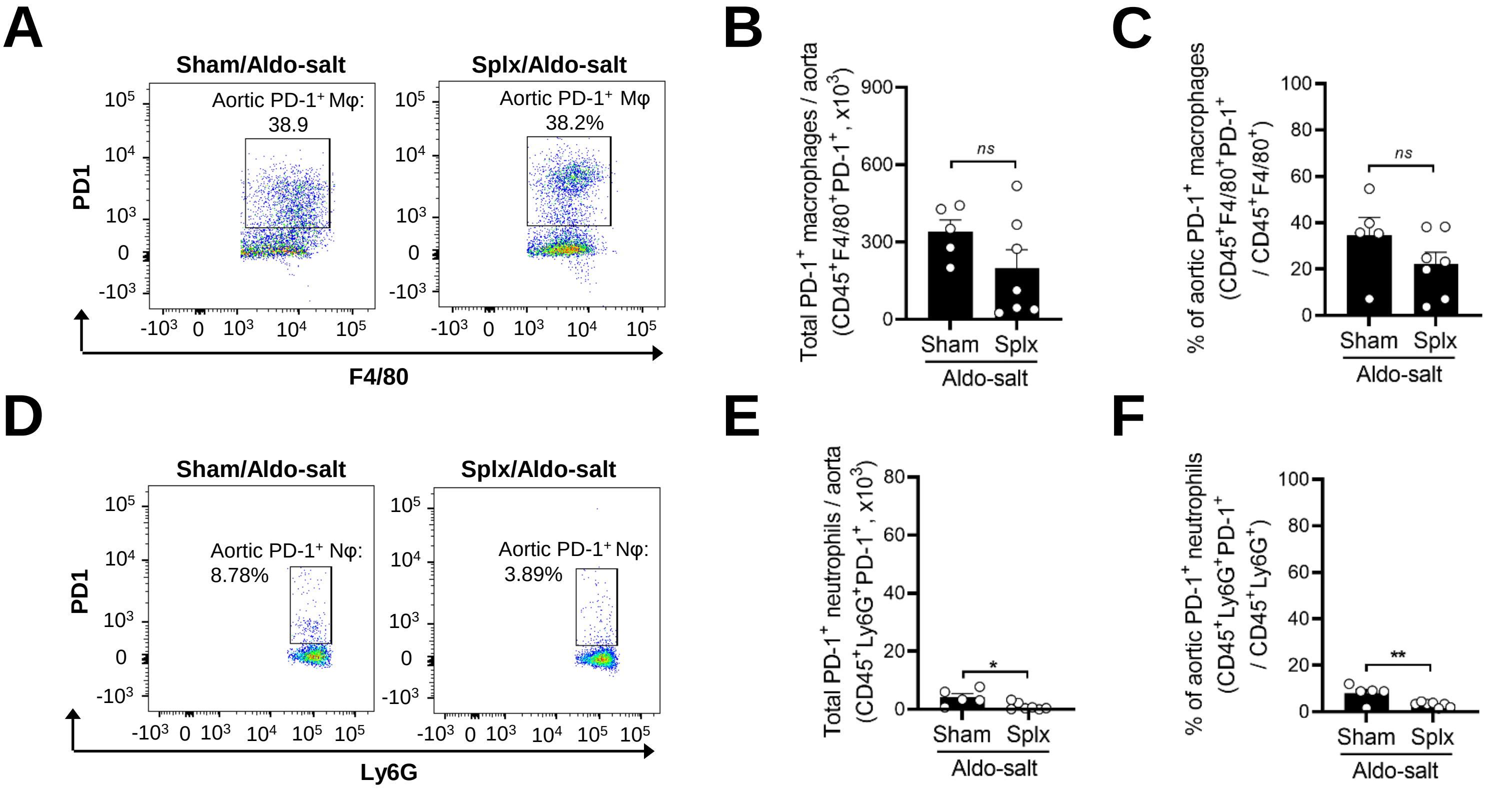


**Supplemental Figure 15. Effects of splenectomy on PD-1 positive macrophages and neutrophils in the aorta of mice administered Aldo-salt.** Representative pseudocolor plots and quantitative data of flow cytometry analysis of the total numbers and percentages of PD-1$^+$ macrophages (Mφ; CD45$^+$F4/80$^+$PD-1$^+$) and neutrophils (CD45$^+$Ly6G$^+$PD-1$^+$) in the whole aortas of 11-13-month-old male C57BL/6J mice with splenectomy (splx) or sham-operation four weeks after Aldo-salt administration. One symbol shown in the figures represents cells in one whole mouse aorta. The data were expressed as mean ± SEM and analyzed by a two-tailed unpaired *t*-test. *, *p<0.05; **, p<0.01. ns, not significant.*

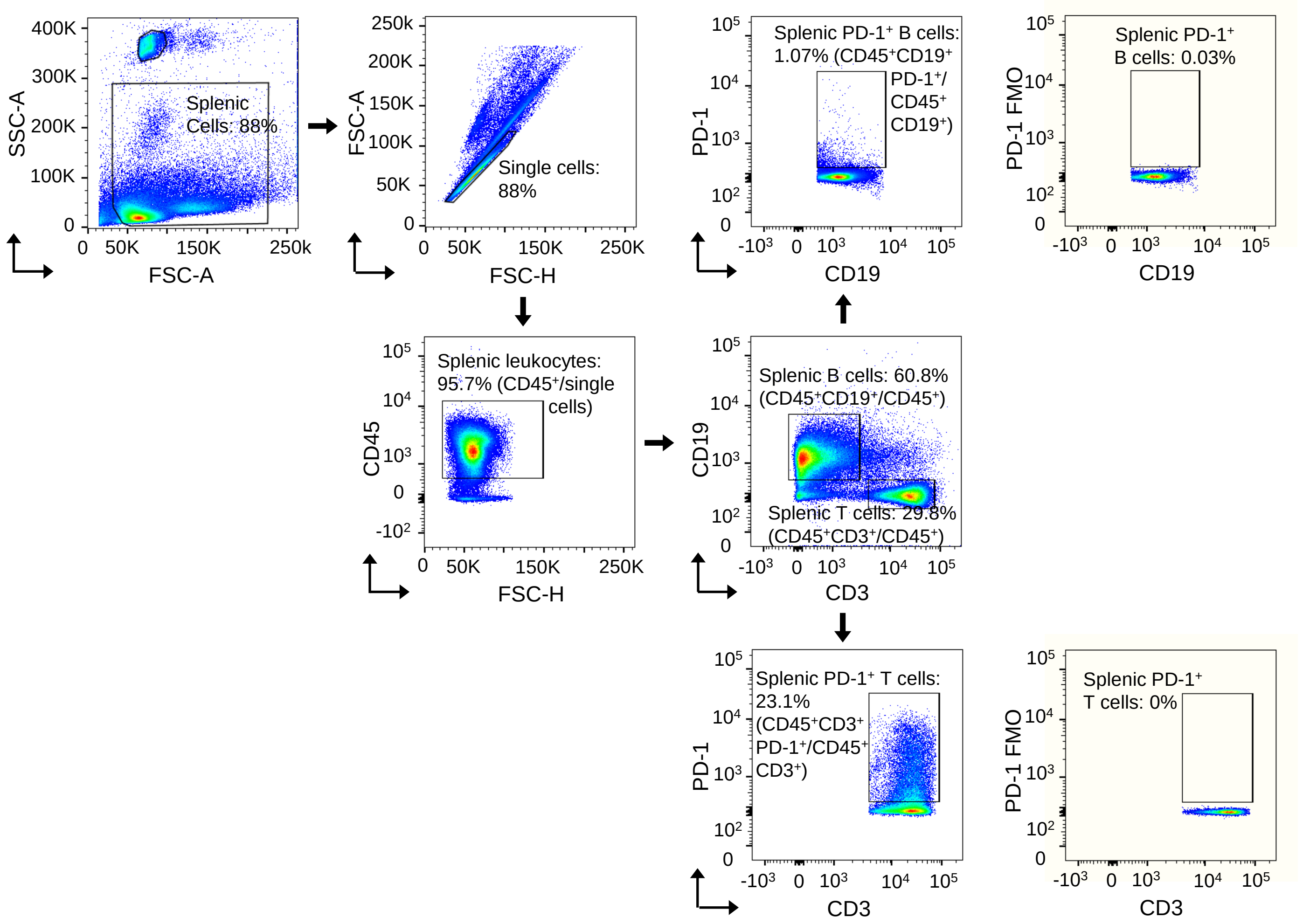


**Supplemental Figure 16. Gating strategy to identify PD1⁺-T cells and PD1⁺- B cells in the mouse spleen.** Single cells were prepared from the mouse spleen and gated in by a forward scatter height (FSC-H) *vs.* forward scatter area (FSC-A) pseudocolor plot. Single cells were gated on CD45⁺ to select leukocytes (CD45⁺) and then gated on CD3⁺ and CD19⁺ to identify T cells (CD45⁺CD3⁺) and B cells (CD45⁺CD19⁺), respectively. Single splenic cells were then gated on PD-1⁺ to identify PD-1⁺ T cells (CD45⁺CD3⁺PD-1⁺) and B cells (CD45⁺CD19⁺PD-1⁺), respectively. Single splenic cells were also analyzed by fluorescence minus one (FMO) (labeled with a yellow color background in pseudocolor plots) to identify gating boundaries and ensure the specificity of antibodies.

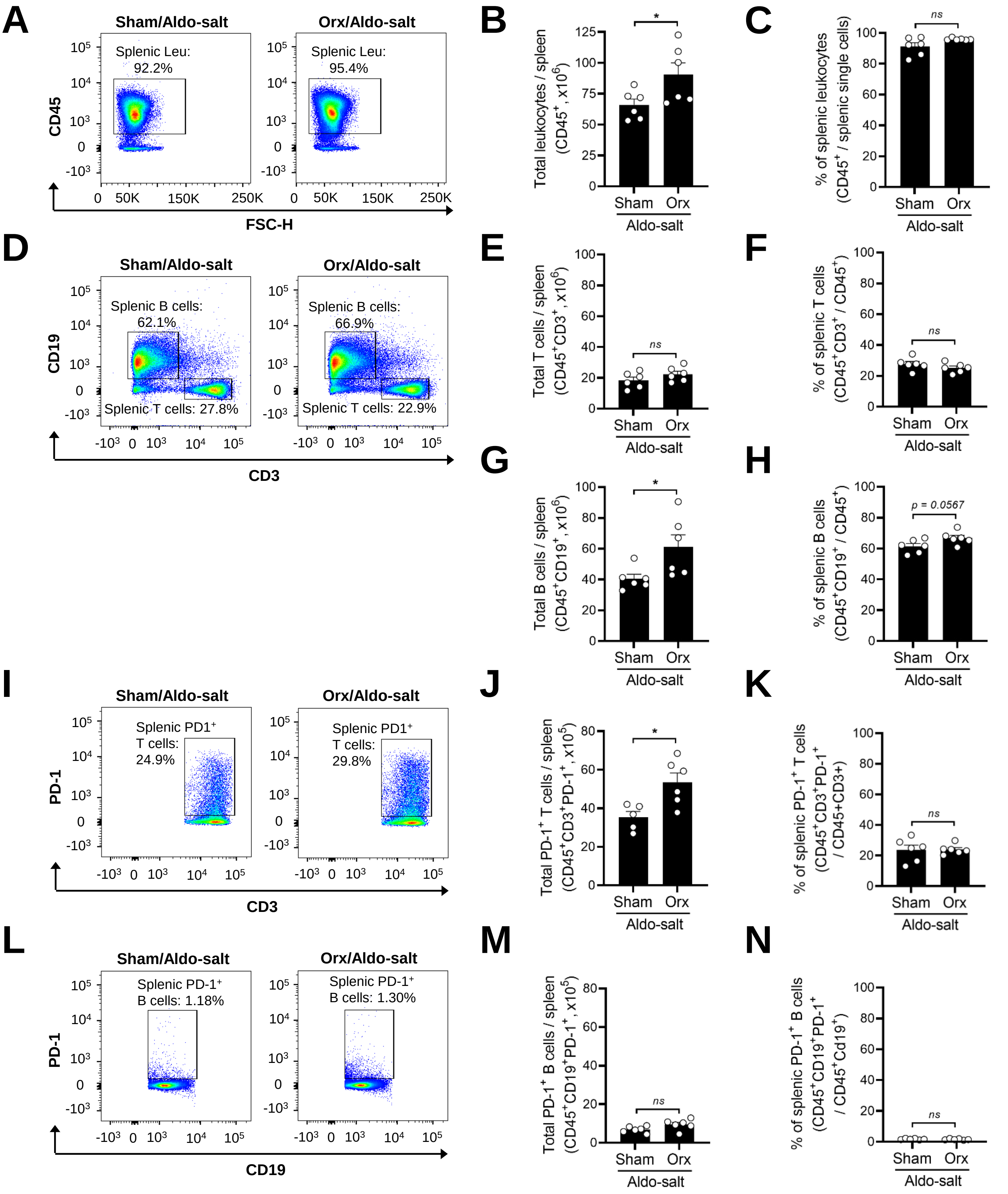


**Supplemental Figure 17. Gonadal androgen deprivation increases the PD-1⁺ T cells, but not PD-1⁺ B cells, in the spleen in mice administered Aldo-salt.** Representative pseudocolor plots and quantitative data of flow cytometry analysis of the total numbers and percentage of leukocytes (Leu; $CD45^+$; **A−C**), T cells ($CD45^+CD3^+$; **D−**F), B cells ($CD45^+CD19^+$; **D**, **G**, and **H**), $PD-1^+$ T cells ($CD45^+CD3^+PD-1^+$; **I−K**), and $PD-1^+$ B cells ($CD45^+CD19^+PD-1^+$; **L−N**) in the spleen from 10-month-old male C57BL/6J mice with orchiectomy (orx) or sham-operation ten days after Aldo-salt administration. The data were expressed as mean ± SEM and analyzed by a two-tailed unpaired *t*-test. **, p<0.05; ns, not significant.*

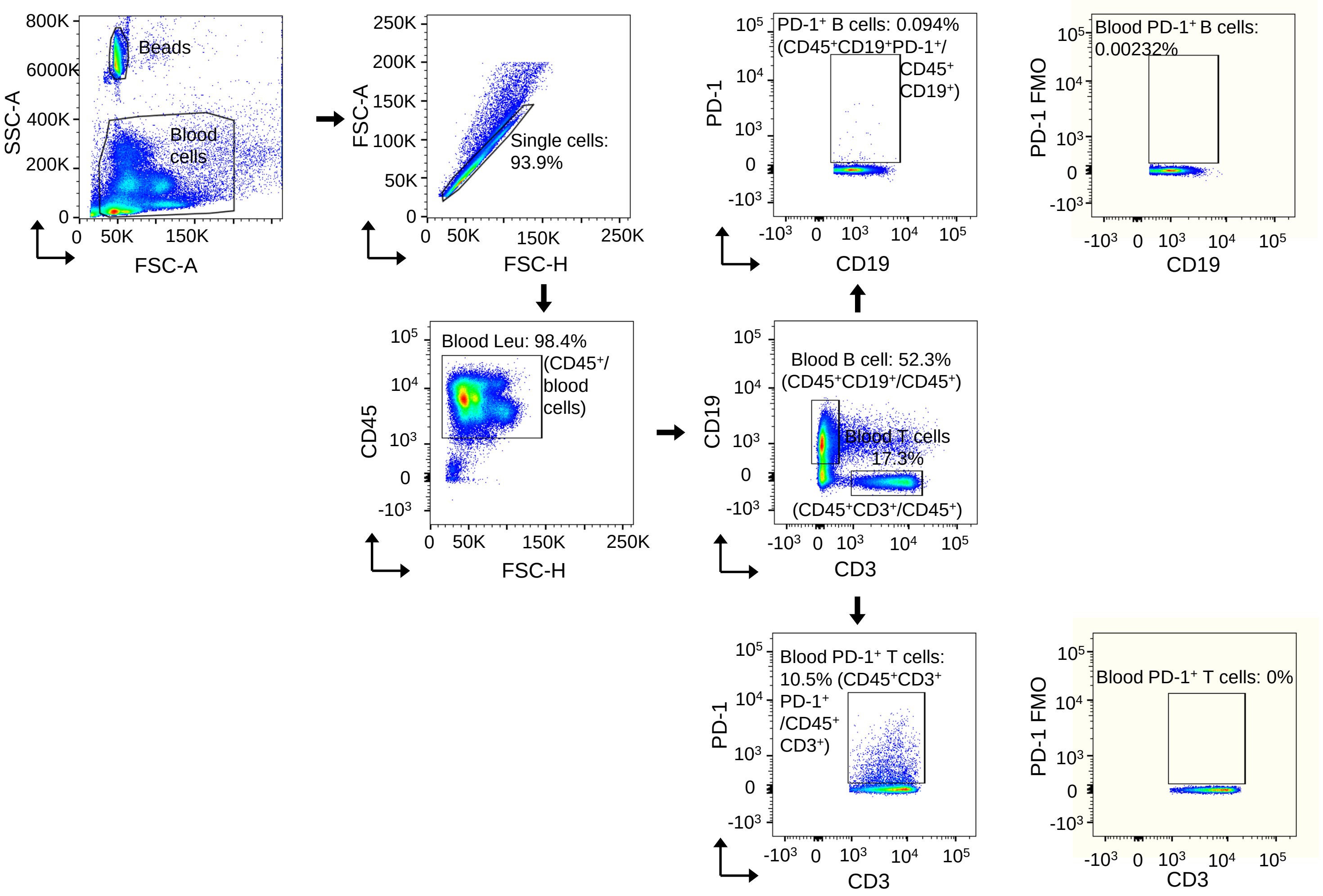


**Supplemental Figure 18. Gating strategy to identify PD-1 positive T cells and PD1$^{+}$- B cells in the mouse blood.** Single cells were prepared from the mouse blood and gated in by a forward scatter height (FSC-H) *vs.* forward scatter area (FSC-A) pseudocolor plot. Single cells were gated on CD45$^{+}$ to select leukocytes (Leu; CD45$^{+}$) and then gated on CD3$^{+}$ and CD19 to identify T cells (CD45$^{+}$CD3$^{+}$) and B cells (CD45$^{+}$CD19$^{+}$), respectively. Single aortic cells were then gated on PD-1$^{+}$ to identify PD-1$^{+}$ T cells (CD45$^{+}$CD3$^{+}$PD-1$^{+}$) and B cells (CD45$^{+}$CD19$^{+}$PD-1$^{+}$), respectively. Single cells from the blood were also analyzed by flow cytometry with fluorescence minus one (FMO) (labeled with a yellow color background in pseudocolor plots) to identify gating boundaries and ensure the specificity of antibodies.

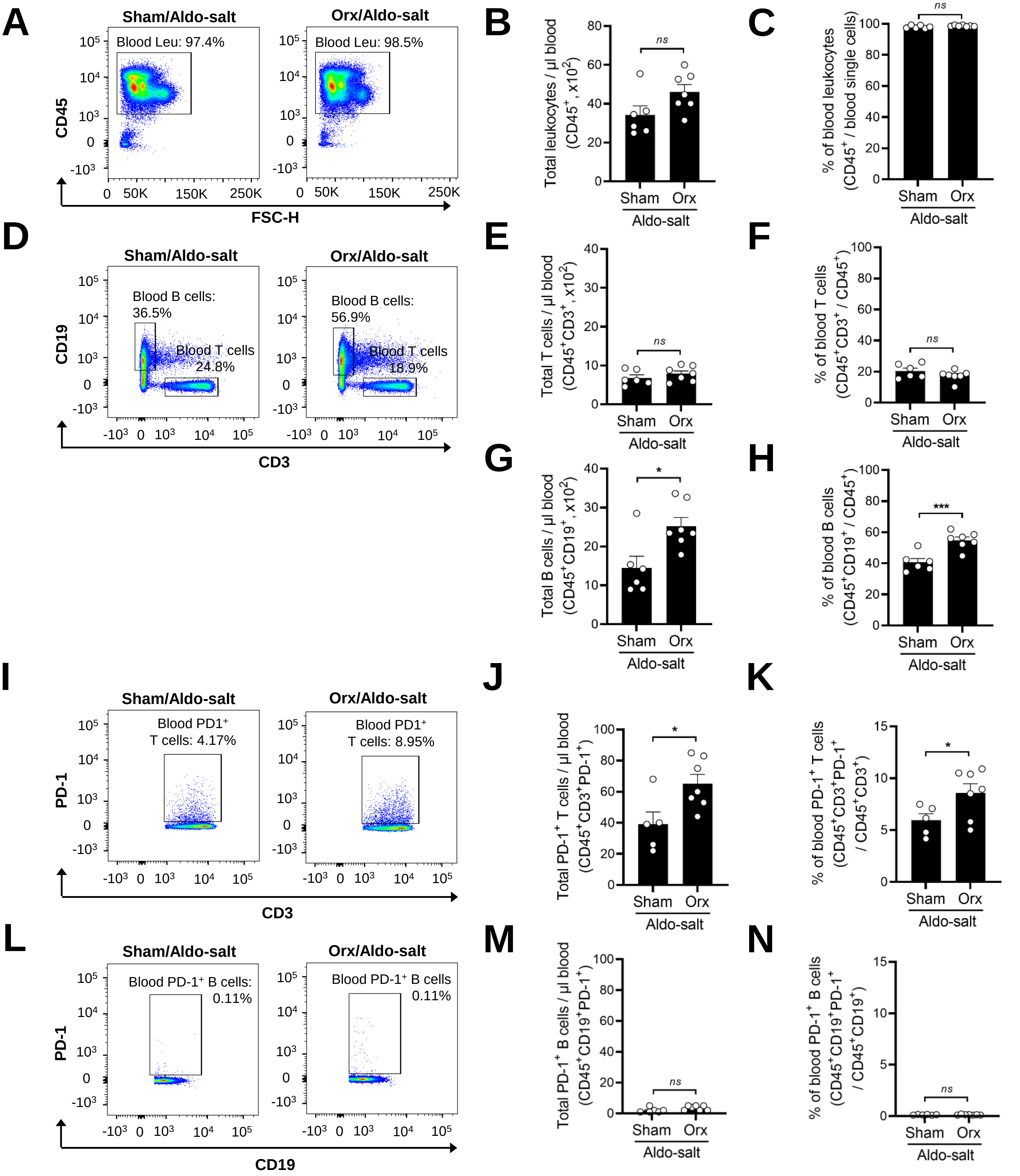


**Supplemental Figure 19. Gonadal androgen deprivation augments PD-1 positive T cells but not PD-1$^+$ B cells in the blood in mice administered Aldo-salt.** Representative pseudocolor plots and quantitative data of flow cytometry analysis of the total numbers and percentage of leukocytes (Leu; CED45$^+$; **A−C**), T cells (CD45$^+$CD3$^+$; **D−**F), B cells (CD45$^+$CD19$^+$; **D**, **G**, and **H**), PD-1$^+$ T cells (CD45$^+$CD3$^+$PD-1$^+$; **I−K**) and PD-1$^+$ B cells (CD45$^+$CD19$^+$PD-1$^+$; **L−N**) in the blood from 10-month-old male C57BL/6J mice with orchiectomy (orx) or sham-operation ten days after Aldo-salt administration. The data were expressed as mean ± SEM and analyzed by a two-tailed unpaired *t*-test. **, p<0.05; ns, not significant.*

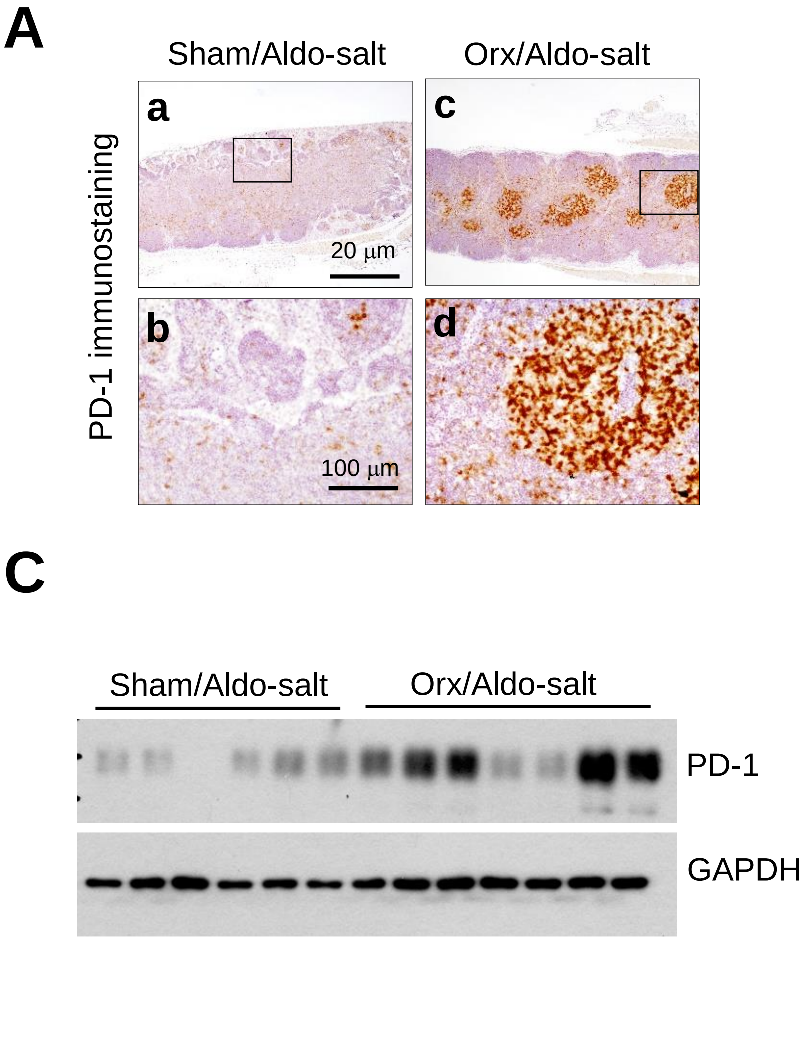


**Supplemental Figure 20. Androgen inhibits PD-1 protein expression in the periaortic lymph nodes in mice administered with Aldo-salt. (A** and **B)** Representative immunostaining (A) and quantitative data (B) of PD-1 protein expression in the periaortic lymph nodes from 10-month-old male C57BL/6J mice with orchiectomy (orx) and sham operation ten days after Aldo-salt administration. The percentage of areas fraction = (the PD-1 positive area / the area of fields of view) x 100%. The data were calculated from five fields of view randomly photographed per lymph node section per mouse. (**C** and **D**) Representative Western blots (C) and quantitative data (D) of PD-1 protein expression in the periaortic lymph nodes from 10-month-old C57BL/6J mice with orx and sham-operated ten days after Aldo-salt administration. PD-1 protein expressions were normalized to GAPDH. The data were expressed as mean ± SEM and analyzed by a two-tailed unpaired *t*-test. *, $p<0.05$.

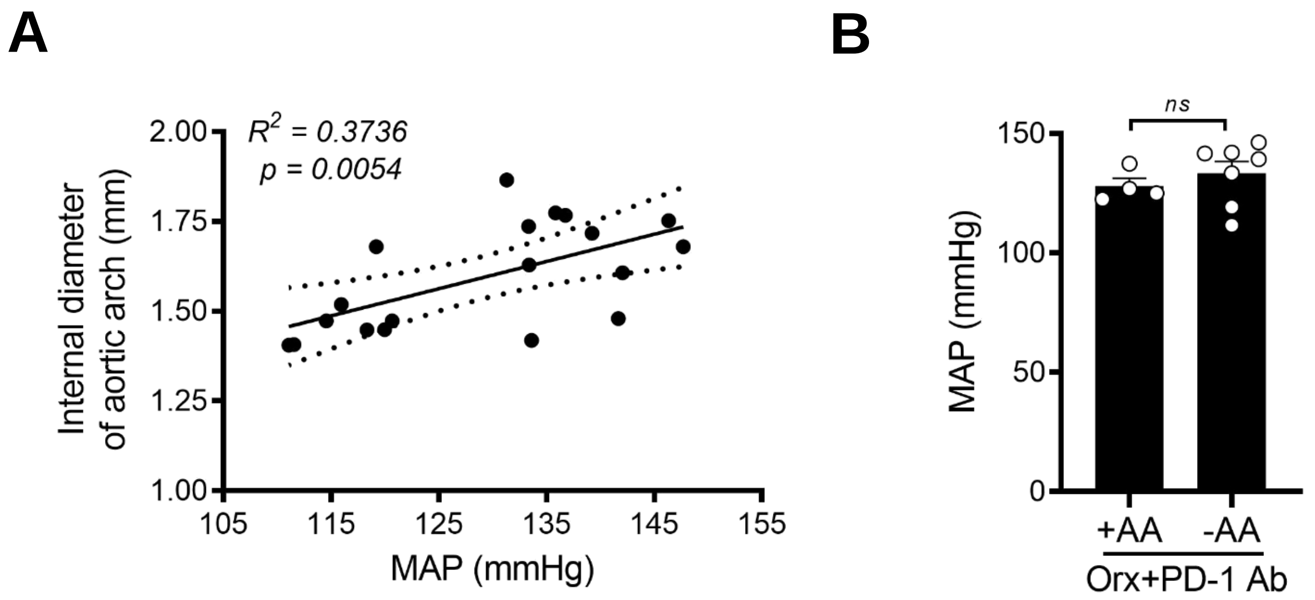


**Supplemental Figure 21. MAP is associated with the internal diameters of the aortic arch but not aortic aneurysms in orchiectomized mice administered Aldo-salt and anti-PD-1 antibody.** (**A**) Correlation analysis of the internal diameter of the aortic arch and MAP in orchiectomized (orx) mice seven weeks in after Aldo-salt and anti-PD-1 antibody (Ab) administration. (**B**) MAP in orchiectomized mice administered Aldo-salt with anti-PD-1 Ab with (+) or without (-) aortic aneurysms (AA). The data were expressed as mean ± SEM and analyzed by simple linear regression analysis (A) and a two-tailed unpaired *t*-test (B). *ns, not significant*

**Supplemental Table 1. A complete list of 180 genes whose mRNAs are upregulated by orchiectomy (orx *vs.* sham ctrl) but downregulated by exogenous dihydrotestosterone (DHT) administration to orchiectomized mice (Orx+DHT *vs.* orx) in the aorta in mice administered Aldo-salt**

| # | Gene ID | Description | Location | Fold Change (orx vs. ctrl) | p-value | Fold Change (orx+DHT vs. orx) | p-value |
|---|---|---|---|---|---|---|---|
| 1 | Krt17 | keratin 17 | chr11 | 72.56 | 0.00293234 | -93.23 | 0.00515206 |
| 2 | Alb | albumin | chr5 | 66.33 | 0.00000639 | -11.24 | 0.00140744 |
| 3 | Tbata | thymus, brain and testes associated | chr10 | 53.87 | 0.00035045 | -28.29 | 0.00098214 |
| 4 | Hamp2 | hepcidin antimicrobial peptide 2 | chr7 | 39.67 | 0.00003697 | -23.43 | 0.00045217 |
| 5 | Camp | cathelicidin antimicrobial peptide | chr9 | 23.37 | 0.00009690 | -4.54 | 0.00397801 |
| 6 | Ngp | neutrophilic granule protein | chr9 | 20.00 | 0.00000009 | -6.09 | 0.00006854 |
| 7 | S100a8 | S100 calcium binding protein A8 (calgranulin A) | chr3 | 13.36 | 0.00000001 | -3.40 | 0.00378026 |
| 8 | H2-Eb2 | histocompatibility 2, class II antigen E beta2 | chr17 | 12.48 | 0.00020575 | -10.62 | 0.00301283 |
| 9 | Themis | thymocyte selection associated | chr10 | 12.16 | 0.00429165 | -18.45 | 0.00022233 |
| 10 | S100a9 | S100 calcium binding protein A9 (calgranulin B) | chr3 | 11.99 | 0.00000000 | -3.20 | 0.00339268 |
| 11 | Myb | myeloblastosis oncogene | chr10 | 10.97 | 0.00009978 | -17.17 | 0.00000386 |
| 12 | Cr2 | complement receptor 2 | chr1 | 10.59 | 0.00073704 | -14.30 | 0.00047904 |
| 13 | Cd79a | CD79A antigen | chr7 | 10.49 | 0.00000089 | -8.41 | 0.00005525 |
| 14 | Ms4a1 | membrane-spanning 4-domains A1 | chr19 | 10.47 | 0.00000850 | -18.94 | 0.00000015 |
| 15 | Cd19 | CD19 antigen | chr7 | 10.22 | 0.00000678 | -10.85 | 0.00001361 |
| 16 | Sh2d1a | SH2 domain containing 1A | chrX | 9.42 | 0.00433350 | -17.30 | 0.00136215 |
| 17 | Tnfrsf13c | TNF receptor superfamily member 13c | chr15 | 9.40 | 0.00022008 | -8.35 | 0.00058788 |
| 18 | Mzb1 | marginal zone B and B1 cell-specific protein 1 | chr18 | 9.10 | 0.00007934 | -10.04 | 0.00012146 |
| 19 | Fcmr | Fc fragment of IgM receptor | chr1 | 9.06 | 0.00055230 | -9.07 | 0.00033054 |
| 20 | Prg2 | proteoglycan 2, bone marrow | chr2 | 8.97 | 0.00009111 | -8.38 | 0.00008549 |
| 21 | Cd8b1 | CD8 antigen, beta chain 1 | chr6 | 8.79 | 0.00000930 | -11.39 | 0.00000187 |
| 22 | Bcl11a | B cell CLL/lymphoma 11A | chr11 | 8.78 | 0.00011468 | -12.34 | 0.00004661 |
| 23 | Gfi1 | growth factor independent 1 transcription repressor | chr5 | 8.54 | 0.00072452 | -4.90 | 0.00809827 |
| 24 | Sidt1 | SID1 transmembrane family member 1 | chr16 | 8.49 | 0.00140464 | -11.35 | 0.00184782 |
| 25 | Cacna1e | calcium voltage-gated channel subunit alpha1 E | chr1 | 8.38 | 0.00018835 | -5.63 | 0.00415678 |
| 26 | Cxcr5 | chemokine (C-X-C motif) receptor 5 | chr9 | 8.36 | 0.00008848 | -10.71 | 0.00009679 |
| 27 | Cd79b | CD79B antigen | chr11 | 8.28 | 0.00000775 | -7.62 | 0.00002434 |
| 28 | Spib | Spi-B transcription factor | chr7 | 8.21 | 0.00012726 | -17.88 | 0.00000381 |
| 29 | Cd8a | CD8 antigen, alpha chain | chr6 | 8.18 | 0.00007024 | -14.10 | 0.00000312 |
| 30 | Gimap7 | GTPase, IMAP family member 7 | chr6 | 7.92 | 0.00242371 | -6.81 | 0.00635858 |
| 31 | Pax5 | paired box 5 | chr4 | 7.79 | 0.00042815 | -7.91 | 0.00004445 |
| 32 | Siglecg | sialic acid binding Ig-like lectin G | chr7 | 7.76 | 0.00059477 | -9.95 | 0.00010210 |
| 33 | Ly6d | lymphocyte antigen 6 complex, locus D | chr15 | 7.74 | 0.00002771 | -8.43 | 0.00001140 |
| 34 | Cd22 | CD22 antigen | chr7 | 7.66 | 0.00036876 | -7.85 | 0.00021901 |
| 35 | Txk | TXK tyrosine kinase | chr5 | 7.58 | 0.00406763 | -15.63 | 0.00131832 |
| 36 | Fcrla | Fc receptor-like A | chr1 | 7.31 | 0.00058840 | -9.12 | 0.00026021 |
| 37 | Pou2af1 | POU class 2 homeobox associating factor 1 | chr9 | 6.47 | 0.00070169 | -9.62 | 0.00001364 |
| 38 | Ltb | lymphotoxin B | chr17 | 6.32 | 0.00005346 | -5.68 | 0.00022554 |
| 39 | Fcrl1 | Fc receptor-like 1 | chr3 | 6.30 | 0.00640415 | -8.28 | 0.00467074 |
| 40 | Ms4a4b | membrane-spanning 4-domains, subfamily A, 4B | chr19 | 6.25 | 0.00054053 | -8.63 | 0.00030896 |
| 41 | Bcl11b | B cell leukemia/lymphoma 11B | chr12 | 6.20 | 0.00022008 | -17.37 | 0.00000158 |
| 42 | H2-Oa | histocompatibility 2, O region alpha locus | chr17 | 6.18 | 0.00530697 | -15.90 | 0.00057125 |
| 43 | Ly6c2 | lymphocyte antigen 6 complex, locus C2 | chr15 | 6.00 | 0.00048563 | -5.97 | 0.00008967 |
| 44 | Slamf6 | SLAM family member 6 | chr1 | 5.99 | 0.00844321 | -7.29 | 0.00744750 |
| 45 | Cd27 | CD27 antigen | chr6 | 5.94 | 0.00088574 | -6.77 | 0.00219890 |
| 46 | Cd5 | CD5 antigen | chr19 | 5.83 | 0.00000845 | -2.99 | 0.00188554 |
| 47 | Gjb2 | gap junction protein, beta 2 | chr14 | 5.62 | 0.00822289 | -8.75 | 0.00898610 |
| 48 | H2-Ob | histocompatibility 2, O region beta locus | chr17 | 5.52 | 0.00051984 | -5.94 | 0.00053131 |
| 49 | Lef1 | lymphoid enhancer binding factor 1 | chr3 | 5.47 | 0.00024262 | -5.34 | 0.00027613 |
| 50 | Ptprcap | protein tyrosine phosphatase receptor type C associated protein | chr19 | 5.44 | 0.00016530 | -7.24 | 0.00002414 |
| 51 | H2-DMb2 | histocompatibility 2, class II, locus Mb2 | chr17 | 5.36 | 0.00006367 | -6.08 | 0.00003221 |
| 52 | Cd69 | CD69 antigen | chr6 | 5.32 | 0.00309230 | -4.77 | 0.00370211 |
| 53 | Cd3g | CD3 antigen, gamma polypeptide | chr9 | 5.30 | 0.00000798 | -4.02 | 0.00040940 |
| 54 | Bcl2a1d | B cell leukemia/lymphoma 2 related protein A1d | chr9 | 5.25 | 0.00053813 | -8.40 | 0.00013498 |
| 55 | Sell | selectin, lymphocyte | chr1 | 5.20 | 0.00121179 | -6.00 | 0.00015825 |
| 56 | Lck | lymphocyte protein tyrosine kinase | chr1 | 5.11 | 0.00053414 | -6.87 | 0.00003226 |
| 57 | Tcf7 | transcription factor 7, T cell specific | chr11 | 4.90 | 0.00015294 | -8.26 | 0.00000127 |
| 58 | Treml4 | triggering receptor expressed on myeloid cells-like 4 | chr17 | 4.84 | 0.00605111 | -4.78 | 0.00891611 |
| 59 | Card11 | caspase recruitment domain family, member 11 | chr5 | 4.77 | 0.00022641 | -9.47 | 0.00001543 |
| 60 | Itk | IL2 inducible T cell kinase | chr11 | 4.66 | 0.00077488 | -10.58 | 0.00000563 |
| 61 | Azgp1 | alpha-2-glycoprotein 1, zinc | chr5 | 4.58 | 0.00226445 | -13.71 | 0.00006137 |
| 62 | Traf3ip3 | TRAF3 interacting protein 3 | chr1 | 4.57 | 0.00122313 | -9.23 | 0.00001125 |
| 63 | Col11a1 | collagen, type XI, alpha 1 | chr3 | 4.53 | 0.00386393 | -4.11 | 0.00926047 |
| 64 | Prss35 | protease, serine 35 | chr9 | 4.47 | 0.00552033 | -4.64 | 0.00871472 |
| 65 | Il18r1 | interleukin 18 receptor 1 | chr1 | 4.47 | 0.00633410 | -13.41 | 0.00002101 |
| 66 | Btla | B and T lymphocyte associated | chr16 | 4.46 | 0.00219236 | -5.82 | 0.00069288 |
| 67 | Ikzf3 | IKAROS family zinc finger 3 | chr11 | 4.42 | 0.00042893 | -11.40 | 0.00000050 |
| 68 | Cd3d | CD3 antigen, delta polypeptide | chr9 | 4.34 | 0.00091830 | -5.79 | 0.00041619 |
| 69 | Cytip | cytohesin 1 interacting protein | chr2 | 4.27 | 0.00125618 | -6.67 | 0.00003467 |
| 70 | Blk | B lymphoid kinase | vhr14 | 4.27 | 0.00543365 | -5.22 | 0.00050444 |

**Supplemental Table 1 (continued). A complete list of 180 genes whose mRNAs are upregulated by orchiectomy (orx *vs.* sham ctrl) but downregulated by exogenous dihydrotestosterone (DHT) administration to orchiectomized mice (Orx+DHT *vs.* orx) in the aorta in mice administered Aldo-salt**

| # | Gene ID | Description | Location | Fold Change (orx vs. ctrl) | p-value | Fold Change (orx+DHT vs. orx) | p-value |
|---|---|---|---|---|---|---|---|
| 71 | Rnase6 | ribonuclease, RNase A family, 6 | chr14 | 4.25 | 0.00062204 | -3.62 | 0.00697734 |
| 72 | Rasal3 | RAS protein activator like 3 | chr17 | 4.18 | 0.00026567 | -5.66 | 0.00009229 |
| 73 | Tcf21 | transcription factor 21 | chr10 | 4.15 | 0.00359346 | -6.98 | 0.00028077 |
| 74 | Cd28 | CD28 antigen | chr1 | 4.10 | 0.00487485 | -11.30 | 0.00015504 |
| 75 | Acap1 | ArfGAP with coiled-coil, ankyrin repeat and PH domains 1 | Chr11 | 4.04 | 0.00049392 | -3.99 | 0.00106126 |
| 76 | Itgax | integrin alpha X | Chr7 | 3.91 | 0.00021341 | -7.78 | 0.00000025 |
| 77 | Fcho1 | FCH domain only 1 | chr8 | 3.77 | 0.00028312 | -5.48 | 0.00004304 |
| 78 | Tbc1d10c | TBC1 domain family, member 10c | chr19 | 3.72 | 0.00335201 | -6.03 | 0.00014369 |
| 79 | Ipcef1 | interaction protein for cytohesin exchange factors 1 | chr10 | 3.72 | 0.00063026 | -4.62 | 0.00033393 |
| 80 | Cdca7 | cell division cycle associated 7 | chr2 | 3.64 | 0.00219618 | -4.19 | 0.00117219 |
| 81 | Rhbg | Rhesus blood group-associated B glycoprotein | chr3 | 3.61 | 0.00461591 | -3.67 | 0.00831695 |
| 82 | Grap2 | GRB2-related adaptor protein 2 | chr15 | 3.59 | 0.00042164 | -4.61 | 0.00008908 |
| 83 | P2ry10 | purinergic receptor P2Y, G-protein coupled 10 | chrX | 3.59 | 0.00518994 | -8.35 | 0.00006476 |
| 84 | Cd3e | CD3 antigen, epsilon polypeptide | chr9 | 3.57 | 0.00107340 | -5.39 | 0.00080476 |
| 85 | Map4k1 | mitogen-activated protein kinase kinase kinase kinase 1 | chr7 | 3.46 | 0.00065314 | -3.55 | 0.00033196 |
| 86 | Gimap3 | GTPase, IMAP family member 3 | chr6 | 3.46 | 0.00052659 | -3.27 | 0.00106921 |
| 87 | Itgal | integrin alpha L | chr7 | 3.39 | 0.00065878 | -5.42 | 0.00000829 |
| 88 | Parvg | parvin, gamma | chr15 | 3.30 | 0.00275671 | -3.46 | 0.00086344 |
| 89 | Rasgrp1 | RAS guanyl releasing protein 1 | chr2 | 3.26 | 0.00013955 | -6.18 | 0.00000003 |
| 90 | Lat | linker for activation of T cells | chr7 | 3.25 | 0.00028462 | -4.26 | 0.00003633 |
| 91 | Ptpn22 | protein tyrosine phosphatase, non-receptor type 22 | chr3 | 3.25 | 0.00017971 | -5.64 | 0.00000163 |
| 92 | Gpr141b | G protein-coupled receptor 141B | chr13 | 3.25 | 0.00714189 | -4.58 | 0.00272678 |
| 93 | Cd2 | CD2 antigen | chr3 | 3.21 | 0.00547508 | -3.89 | 0.00071948 |
| 94 | Skap1 | src family associated phosphoprotein 1 | chr11 | 3.15 | 0.00086272 | -5.52 | 0.00005163 |
| 95 | Unc13d | unc-13 homolog D | chr11 | 3.12 | 0.00115720 | -2.97 | 0.00141585 |
| 96 | Cd52 | CD52 antigen | chr4 | 3.06 | 0.00053838 | -3.10 | 0.00054816 |
| 97 | Il7r | interleukin 7 receptor | chr15 | 3.03 | 0.00465978 | -5.88 | 0.00004590 |
| 98 | Cxcr6 | chemokine (C-X-C motif) receptor 6 | chr9 | 3.00 | 0.00662081 | -7.99 | 0.00000974 |
| 99 | Sla | src-like adaptor | chr15 | 3.00 | 0.00031091 | -4.40 | 0.00000104 |
| 100 | Itgb7 | integrin beta 7 | chr15 | 2.90 | 0.00292785 | -4.49 | 0.00001454 |
| 101 | Stat4 | signal transducer and activator of transcription 4 | chr1 | 2.86 | 0.00813945 | -7.63 | 0.00002618 |
| 102 | Bcl2a1b | B cell leukemia/lymphoma 2 related protein A1b | chr9 | 2.84 | 0.00505359 | -4.53 | 0.00043336 |
| 103 | Gbp8 | guanylate-binding protein 8 | chr5 | 2.84 | 0.00241229 | -6.61 | 0.00000543 |
| 104 | Nup210 | nucleoporin 210 | chr6 | 2.80 | 0.00257918 | -2.49 | 0.00690455 |
| 105 | Cd37 | CD37 antigen | chr7 | 2.80 | 0.00119792 | -3.14 | 0.00034960 |
| 106 | Rhoh | ras homolog family member H | chr5 | 2.78 | 0.00255259 | -3.77 | 0.00029389 |
| 107 | Pmaip1 | phorbol-12-myristate-13-acetate-induced protein 1 | chr18 | 2.77 | 0.00940216 | -4.14 | 0.00081350 |
| 108 | Il2rb | interleukin 2 receptor, beta chain | chr15 | 2.56 | 0.00166477 | -3.37 | 0.00009137 |
| 109 | Wdfy4 | WD repeat and FYVE domain containing 4 | chr14 | 2.54 | 0.00147824 | -4.00 | 0.00000055 |
| 110 | Il21r | interleukin 21 receptor | chr7 | 2.52 | 0.00218437 | -3.53 | 0.00000894 |
| 111 | Cyp4f18 | cytochrome P450, family 4, subfamily f, polypeptide 18 | chr8 | 2.52 | 0.00593724 | -3.69 | 0.00057901 |
| 112 | Ezr | ezrin | chr17 | 2.52 | 0.00921918 | -4.72 | 0.00001711 |
| 113 | Coro1a | coronin, actin binding protein 1A | chr7 | 2.49 | 0.00009215 | -2.88 | 0.00001024 |
| 114 | Cd247 | CD247 antigen | chr1 | 2.49 | 0.00011772 | -3.17 | 0.00006881 |
| 115 | Mamdc2 | MAM domain containing 2 | chr19 | 2.46 | 0.00309219 | -2.72 | 0.00103849 |
| 116 | Blnk | B cell linker | chr19 | 2.43 | 0.00651873 | -3.26 | 0.00042273 |
| 117 | Arl5c | ADP-ribosylation factor-like 5C | chr11 | 2.42 | 0.00540941 | -2.40 | 0.00739382 |
| 118 | Sash3 | SAM and SH3 domain containing 3 | chrX | 2.38 | 0.00115611 | -3.04 | 0.00005892 |
| 119 | Rgs14 | regulator of G-protein signaling 14 | chr13 | 2.38 | 0.00720522 | -4.80 | 0.00003052 |
| 120 | Rac2 | Rac family small GTPase 2 | chr15 | 2.31 | 0.00109290 | -2.70 | 0.00034669 |
| 121 | Irf8 | interferon regulatory factor 8 | chr8 | 2.31 | 0.00992903 | -3.21 | 0.00020599 |
| 122 | H2-DMa | histocompatibility 2, class II, locus DMa | chr17 | 2.30 | 0.00256607 | -2.50 | 0.00221721 |
| 123 | Ptprc | protein tyrosine phosphatase receptor type C | chr1 | 2.29 | 0.00466891 | -4.49 | 0.00000005 |
| 124 | H2-Ab1 | histocompatibility 2, class II antigen A, beta 1 | chr17 | 2.28 | 0.00279696 | -2.65 | 0.00038114 |
| 125 | Shisa2 | shisa family member 2 | chr14 | 2.28 | 0.00544805 | -2.51 | 0.00417194 |
| 126 | Cd74 | CD74 antigen | chr18 | 2.24 | 0.00281218 | -2.53 | 0.00042047 |
| 127 | Hvcn1 | hydrogen voltage-gated channel 1 | chr5 | 2.24 | 0.00437503 | -2.28 | 0.00306926 |
| 128 | Ikzf1 | IKAROS family zinc finger 1 | chr11 | 2.22 | 0.00263754 | -3.87 | 0.00000016 |
| 129 | Psd4 | pleckstrin and Sec7 domain containing 4 | chr2 | 2.18 | 0.00345950 | -2.98 | 0.00019628 |
| 130 | Spn | sialophorin | chr7 | 2.14 | 0.00363685 | -3.06 | 0.00000287 |
| 131 | Hmgb2 | high mobility group box 2 | chr8 | 2.14 | 0.00003441 | -2.26 | 0.00000462 |
| 132 | Selplg | selectin, platelet (p-selectin) ligand | chr5 | 2.13 | 0.00314154 | -2.85 | 0.00012451 |
| 133 | Dlgap5 | DLG associated protein 5 | chr14 | 2.12 | 0.00762526 | -4.69 | 0.00001236 |
| 134 | H2-Aa | histocompatibility 2, class II antigen A, alpha | chr17 | 2.09 | 0.00643965 | -2.23 | 0.00426046 |
| 135 | Sema4d | semaphorin 4D | chr13 | 2.07 | 0.00119369 | -2.21 | 0.00083843 |
| 136 | Mpeg1 | macrophage expressed gene 1 | chr19 | 2.07 | 0.00739749 | -2.95 | 0.00001279 |
| 137 | Myo1g | myosin IG | chr11 | 2.05 | 0.00819989 | -3.85 | 0.00000032 |
| 138 | Pclaf | PCNA clamp associated factor | chr9 | 2.00 | 0.00922412 | -3.65 | 0.00000918 |
| 139 | Ptpn6 | protein tyrosine phosphatase, non-receptor type 6 | chr6 | 2.00 | 0.00882316 | -2.98 | 0.00010761 |
| 140 | Kif11 | kinesin family member 11 | chr19 | 1.97 | 0.00320371 | -4.96 | 0.00000001 |

**Supplemental Table 1 (continued). A complete list of 180 genes whose mRNAs are upregulated by orchiectomy (orx *vs.* sham ctrl) but downregulated by exogenous dihydrotestosterone (DHT) administration to orchiectomized mice (Orx+DHT *vs.* orx) in the aorta in mice administered Aldo-salt**

| # | Gene ID | Description | Location | Fold Change (orx vs. ctrl) | p-value | Fold Change (orx+DHT vs. orx) | p-value |
|---|---|---|---|---|---|---|---|
| 141 | St3gal6 | ST3 beta-galactoside alpha-2,3-sialyltransferase 6 | chr16 | 1.95 | 0.00015243 | -2.99 | 0.00000000 |
| 142 | Slc7a10 | solute carrier family 7 member 10 | chr7 | 1.91 | 0.00003684 | -2.91 | 0.00000657 |
| 143 | Endou | endonuclease, polyU-specific | chr15 | 1.90 | 0.00958322 | -2.30 | 0.00131035 |
| 144 | Top2a | topoisomerase (DNA) II alpha | chr11 | 1.88 | 0.00650663 | -3.78 | 0.00000001 |
| 145 | Mthfd1l | methylenetetrahydrofolate dehydrogenase 1-like | chr10 | 1.85 | 0.00002071 | -1.48 | 0.00306873 |
| 146 | Arhgap45 | Rho GTPase activating protein 45 | chr10 | 1.83 | 0.00825779 | -2.20 | 0.00039083 |
| 147 | Nusap1 | nucleolar and spindle associated protein 1 | chr2 | 1.83 | 0.00103524 | -2.44 | 0.00004059 |
| 148 | Ccr2 | chemokine (C-C motif) receptor 2 | chr9 | 1.81 | 0.00949725 | -3.16 | 0.00000078 |
| 149 | Tbx20 | T-box 20 | chr9 | 1.81 | 0.00592786 | -2.08 | 0.00196389 |
| 150 | Lat2 | linker for activation of T cells family, member 2 | chr5 | 1.79 | 0.00805564 | -2.53 | 0.00004088 |
| 151 | Bmp3 | bone morphogenetic protein 3 | chr5 | 1.79 | 0.00004372 | -1.45 | 0.00473755 |
| 152 | Rrm2 | ribonucleotide reductase M2 | chr12 | 1.77 | 0.00727620 | -2.80 | 0.00006412 |
| 153 | Galnt6 | polypeptide N-acetylgalactosaminyltransferase 6 | chr15 | 1.76 | 0.00672652 | -2.33 | 0.00018578 |
| 154 | Alcam | activated leukocyte cell adhesion molecule | chr16 | 1.72 | 0.00193548 | -1.84 | 0.00039358 |
| 155 | Cdca8 | cell division cycle associated 8 | chr4 | 1.71 | 0.00379456 | -2.31 | 0.00075954 |
| 156 | Cpxm1 | carboxypeptidase X 1 (M14 family) | vhr2 | 1.68 | 0.00002026 | -2.04 | 0.00001212 |
| 157 | Irf5 | interferon regulatory factor 5 | chr6 | 1.65 | 0.00422174 | -2.51 | 0.00000055 |
| 158 | Cpz | carboxypeptidase Z | chr5 | 1.62 | 0.00211843 | -2.33 | 0.00009330 |
| 159 | Kifc1 | kinesin family member C1 | chr17 | 1.61 | 0.00700034 | -2.47 | 0.00008529 |
| 160 | Sox4 | SRY (sex determining region Y)-box 4 | chr13 | 1.58 | 0.00208812 | -1.74 | 0.00020357 |
| 161 | Gpc3 | glypican 3 | chrX | 1.52 | 0.00073456 | -3.09 | 0.00000000 |
| 162 | Arl4c | ADP-ribosylation factor-like 4C | chr1 | 1.50 | 0.00835630 | -1.84 | 0.00005559 |
| 163 | Ctsk | cathepsin K | chr3 | 1.49 | 0.00252571 | -1.84 | 0.00014613 |
| 164 | St3gal4 | ST3 beta-galactoside alpha-2,3-sialyltransferase 4 | chr9 | 1.48 | 0.00249456 | -1.57 | 0.00090671 |
| 165 | Efemp1 | EGF-containing fibulin-like extracellular matrix protein 1 | chr11 | 1.48 | 0.00960942 | -2.46 | 0.00000004 |
| 166 | Chst2 | carbohydrate sulfotransferase 2 | chr9 | 1.43 | 0.00706828 | -1.56 | 0.00174383 |
| 167 | Mcm4 | minichromosome maintenance complex component 4 | chr16 | 1.42 | 0.00303205 | -1.45 | 0.00020502 |
| 168 | Mcm3 | minichromosome maintenance complex component 3 | chr1 | 1.40 | 0.00005370 | -1.63 | 0.00000000 |
| 169 | Mmp2 | matrix metallopeptidase 2 | chr19 | 1.35 | 0.00322527 | -1.90 | 0.00000004 |
| 170 | Prcp | prolylcarboxypeptidase (angiotensinase C) | chr7 | 1.33 | 0.00586741 | -1.58 | 0.00000028 |
| 171 | Sparcl1 | SPARC-like 1 | chr5 | 1.31 | 0.00162648 | -1.39 | 0.00000000 |
| 172 | Rcn1 | reticulocalbin 1 | chr2 | 1.31 | 0.00273570 | -1.57 | 0.00011684 |
| 173 | Cmtm3 | CKLF-like MARVEL transmembrane domain containing 3 | chr8 | 1.29 | 0.00967474 | -1.47 | 0.00487152 |
| 174 | C1ra | complement component 1, r subcomponent A | chr6 | 1.28 | 0.00999725 | -1.71 | 0.00000593 |
| 175 | Pdcd4 | programmed cell death 4 | chr19 | 1.26 | 0.00654881 | -1.40 | 0.00017664 |
| 176 | Pcna | proliferating cell nuclear antigen | chr2 | 1.24 | 0.00890499 | -1.42 | 0.00001990 |
| 177 | Hmgb1 | high mobility group box 1 | chr5 | 1.24 | 0.00172916 | -1.30 | 0.00004352 |
| 178 | Pon2 | paraoxonase 2 | chr6 | 1.23 | 0.00892161 | -1.34 | 0.00006575 |
| 179 | Alyref | Aly/REF export factor | chr11 | 1.23 | 0.00625671 | -1.32 | 0.00066921 |
| 180 | Aqp1 | aquaporin 1 | chr6 | 1.17 | 0.00964582 | -1.34 | 0.00000000 |

#### Supplemental Table 2. A complete list of 150 genes whose mRNAs are downregulated by orchiectomy (orx *vs.* sham ctrl) but upregulated by exogenous dihydrotestosterone (DHT) administration to orchiectomized mice (Orx+DHT *vs.* orx) in the aorta in mice administered Aldo-salt

| # | Gene ID | Description | Location | Fold Change (orx vs. ctrl) | p-value | Fold Change (orx+DHT vs. orx) | p-value |
|---|---|---|---|---|---|---|---|
| 1 | Myh4 | myosin heavy chain 4, skeletal muscle | chr11 | -26.73655093 | 0.00006572 | 13.92182629 | 0.00655175 |
| 2 | Cacng6 | calcium channel, voltage-dependent, gamma subunit 6 | chr7 | -24.8409316 | 0.00276263 | 34.25279623 | 0.00031960 |
| 3 | Slco1a1 | solute carrier organic anion transporter family, member 1a1 | chr6 | -17.09886947 | 0.00233895 | 19.12046232 | 0.00069800 |
| 4 | Scd3 | stearoyl-coenzyme A desaturase 3 | chr19 | -15.297534 | 0.00034029 | 10.74260432 | 0.00185519 |
| 5 | Gm45915 | immune system associated SCR family kinase partner | chr14 | -12.55500926 | 0.00721071 | 18.11818619 | 0.00484563 |
| 6 | Atp2a1 | ATPase, Ca++ transporting, cardiac muscle, fast twitch 1 | chr7 | -9.529129404 | 0.00221234 | 11.66179158 | 0.00002494 |
| 7 | Bmp8b | bone morphogenetic protein 8b | chr4 | -8.942783046 | 0.00154569 | 6.639997161 | 0.00049010 |
| 8 | Pvalb | parvalbumin | chr15 | -8.822837329 | 0.00479196 | 6.289348289 | 0.00563687 |
| 9 | Hephl1 | hephaestin-like 1 | chr9 | -8.50672212 | 0.00000000 | 7.173434607 | 0.00000007 |
| 10 | Mybpc1 | myosin binding protein C, slow-type | chr10 | -7.475004077 | 0.00826108 | 24.30187667 | 0.00000293 |
| 11 | Rspo4 | R-spondin 4 | chr2 | -5.295639174 | 0.00762548 | 6.232415488 | 0.00003637 |
| 12 | Tnnc2 | troponin C2, fast | chr2 | -5.279964009 | 0.00430892 | 8.774819367 | 0.00000080 |
| 13 | Actn3 | actinin alpha 3 | chr19 | -5.149314201 | 0.00176955 | 4.103126291 | 0.00081762 |
| 14 | Mylpf | myosin light chain, phosphorylatable, fast skeletal muscle | chr7 | -5.030485019 | 0.00066253 | 7.205542241 | 0.00000006 |
| 15 | Adcy10 | adenylate cyclase 10 | chr1 | -4.824251965 | 0.00009234 | 3.767798349 | 0.00021866 |
| 16 | 6430571L13Rik | RIKEN cDNA 6430571L13 gene | chr9 | -4.779242313 | 0.00000351 | 3.82626693 | 0.00000183 |
| 17 | Slc15a5 | solute carrier family 15, member 5 | chr6 | -4.731133154 | 0.00120820 | 9.931236506 | 0.00000019 |
| 18 | Myoz1 | myozenin 1 | chr14 | -4.198139313 | 0.00400330 | 4.115450402 | 0.00023289 |
| 19 | Neb | nebulin | chr2 | -3.97642231 | 0.00010067 | 6.946361658 | 0.00000000 |
| 20 | Slc2a5 | solute carrier family 2 member 5 | chr4 | -3.772616571 | 0.00130678 | 4.822316769 | 0.00000321 |
| 21 | Marc1 | mitochondrial amidoxime reducing component 1 (Mtarc1) | chr1 | -3.70937506 | 0.00011831 | 3.852321405 | 0.00000015 |
| 22 | Tfr2 | transferrin receptor 2 | chr5 | -3.702884638 | 0.00008953 | 3.178095483 | 0.00036694 |
| 23 | 2310069B03Rik | RIKEN cDNA 2310069B03 gene | chr6 | -3.430601918 | 0.00442447 | 2.879139611 | 0.00067184 |
| 24 | Mlxipl | MLX interacting protein-like | chr5 | -3.304635818 | 0.00000362 | 2.771832582 | 0.00014300 |
| 25 | Hapln4 | hyaluronan and proteoglycan link protein 4 | chr8 | -3.295089637 | 0.00001340 | 4.638451257 | 0.00012873 |
| 26 | Pnpla3 | patatin-like phospholipase domain containing 3 | chr15 | -3.27063947 | 0.00000031 | 2.301166383 | 0.00013270 |
| 27 | Gm11520 | predicted gene 11520 | chr11 | -3.193152912 | 0.00517541 | 2.603489383 | 0.00575359 |
| 28 | Ttc25 | outer dynein arm complex subunit 4 (Odad4) | chr11 | -3.188548614 | 0.00282747 | 3.386327104 | 0.00003446 |
| 29 | Fam57b | TLC domain containing 3B (Tlcd3b) | chr7 | -3.051711678 | 0.00065974 | 2.534739256 | 0.00119985 |
| 30 | Elovl3 | ELOVL fatty acid elongase 3 | chr19 | -2.878781262 | 0.00170205 | 2.572605219 | 0.00055543 |
| 31 | Odf3l1 | outer dense fiber of sperm tails 3-like 1 | chr9 | -2.858605255 | 0.00210978 | 2.613562674 | 0.00334420 |
| 32 | Angptl8 | angiopoietin-like 8 | chr9 | -2.830001395 | 0.00044141 | 2.342879714 | 0.00000034 |
| 33 | Acaca | acetyl-Coenzyme A carboxylase alpha | chr11 | -2.825987526 | 0.00006213 | 2.328369192 | 0.00116391 |
| 34 | Ctcfl | CCCTC-binding factor (zinc finger protein)-like | chr2 | -2.822863407 | 0.00004196 | 3.147440361 | 0.00007848 |
| 35 | Scd1 | stearoyl-Coenzyme A desaturase 1 | chr19 | -2.79365452 | 0.00000214 | 2.024278423 | 0.00035163 |
| 36 | Acly | ATP citrate lyase | chr11 | -2.786751051 | 0.00024856 | 2.231941973 | 0.00211594 |
| 37 | Acacb | acetyl-Coenzyme A carboxylase beta | chr5 | -2.730347215 | 0.00006324 | 2.345124918 | 0.00069934 |
| 38 | Fasn | fatty acid synthase | chr11 | -2.711814031 | 0.00027666 | 2.360887753 | 0.00221614 |
| 39 | Acot11 | acyl-CoA thioesterase 11 | chr4 | -2.699798185 | 0.00033497 | 2.303976145 | 0.00054574 |
| 40 | Hk2 | hexokinase 2 | chr6 | -2.695666378 | 0.00000499 | 2.28503619 | 0.00003183 |
| 41 | Ncan | neurocan | chr8 | -2.670425807 | 0.00009096 | 13.10809959 | 0.00000233 |
| 42 | Kng2 | kininogen 2 | chr16 | -2.631043584 | 0.00190526 | 2.38354522 | 0.00392413 |
| 43 | Impa2 | inositol monophosphatase 2 | chr18 | -2.599388357 | 0.00033038 | 2.340899643 | 0.00138838 |
| 44 | Ppp1r3b | protein phosphatase 1, regulatory subunit 3B | chr8 | -2.585642632 | 0.00010491 | 1.868952864 | 0.00206128 |
| 45 | Gm32200 | predicted gene, 32200 | chr1 | -2.579560512 | 0.00059940 | 2.384251324 | 0.00031380 |
| 46 | Plin2 | perilipin 2 | chr4 | -2.57301374 | 0.00036310 | 2.56262567 | 0.00000827 |
| 47 | Elovl6 | ELOVL fatty acid elongase 6 | chr3 | -2.563104392 | 0.00178066 | 2.31783194 | 0.00214619 |
| 48 | B430212C06Rik | RIKEN cDNA B430212C06 gene | chr18 | -2.555155617 | 0.00465133 | 2.191188656 | 0.00387148 |
| 49 | Deptor | DEP domain containing MTOR-interacting protein | chr15 | -2.534030055 | 0.00010325 | 2.283292166 | 0.00060302 |
| 50 | Orm1 | orosomucoid 1 | chr4 | -2.487835722 | 0.00029616 | 1.930801808 | 0.00018227 |
| 51 | Ybx2 | Y box protein 2 | chr11 | -2.372547721 | 0.00068539 | 2.497398859 | 0.00016660 |
| 52 | C7 | complement component 7 | chr15 | -2.364556185 | 0.00299812 | 2.582352233 | 0.00371133 |
| 53 | Me1 | malic enzyme 1, NADP(+)-dependent, cytosolic | chr9 | -2.312458522 | 0.00147527 | 2.197144045 | 0.00007772 |
| 54 | Tmem79 | transmembrane protein 79 | chr3 | -2.308283014 | 0.00247910 | 2.308631183 | 0.00046048 |
| 55 | Mybpc2 | myosin binding protein C, fast-type | chr7 | -2.303774322 | 0.00788846 | 2.575690918 | 0.00017106 |
| 56 | Nat8l | N-acetyltransferase 8-like | chr5 | -2.303048643 | 0.00078200 | 2.36004515 | 0.00045022 |
| 57 | Agpat2 | 1-acylglycerol-3-phosphate O-acyltransferase 2 | chr2 | -2.298558276 | 0.00191509 | 2.177002225 | 0.00036141 |
| 58 | Lpin1 | lipin 1 | chr12 | -2.294161149 | 0.00010579 | 1.851453715 | 0.00105545 |
| 59 | Acss2 | acyl-CoA synthetase short-chain family member 2 | chr2 | -2.292827529 | 0.00169793 | 1.829010795 | 0.00664079 |
| 60 | Sbk1 | SH3-binding kinase 1 | chr7 | -2.290899165 | 0.00041209 | 2.155875982 | 0.00016710 |
| 61 | Pfkl | phosphofructokinase, liver, B-type | chr10 | -2.271973256 | 0.00054044 | 2.306888751 | 0.00004527 |
| 62 | Paqr9 | progestin and adipoQ receptor family member IX | chr9 | -2.265359905 | 0.00101696 | 1.911965951 | 0.00644233 |
| 63 | Slc16a1 | solute carrier family 16 member 1 | chr3 | -2.262846428 | 0.00097938 | 1.655724294 | 0.00883555 |
| 64 | Sfxn5 | sideroflexin 5 | chr6 | -2.252235361 | 0.00108356 | 1.833943093 | 0.00571332 |
| 65 | Ibsp | integrin binding sialoprotein | chr5 | -2.234168899 | 0.00231733 | 4.716266365 | 0.00000000 |
| 66 | Gpd1 | glycerol-3-phosphate dehydrogenase 1 | chr15 | -2.217866746 | 0.00302967 | 2.132792299 | 0.00334719 |
| 67 | Plin5 | perilipin 5 | chr17 | -2.191033023 | 0.00393885 | 1.996942247 | 0.00169445 |
| 68 | Hoxc5 | homeobox C5 | chr15 | -2.186362002 | 0.00085246 | 2.204316809 | 0.00391005 |
| 69 | Tusc5 | trafficking regulator of GLUT4 (SLC2A4) 1 (Trarg1) | chr11 | -2.157952745 | 0.00113431 | 2.115138551 | 0.00027939 |
| 70 | Acsf3 | acyl-CoA synthetase family member 3 | chr8 | -2.132420125 | 0.00240348 | 2.222633743 | 0.00035874 |

**Supplemental Table 2 (continued). A complete list of 150 genes whose mRNAs are downregulated by orchiectomy (orx *vs.* sham ctrl) but upregulated by exogenous dihydrotestosterone (DHT) administration to orchiectomized mice (Orx+DHT *vs.* orx) in the aorta in mice administered Aldo-salt**

| # | Gene ID | Description | Location | Fold Change (orx vs. ctrl) | p-value | Fold Change (orx+DHT vs. orx) | p-value |
|---|---|---|---|---|---|---|---|
| 71 | Pgd | phosphogluconate dehydrogenase | chr4 | -2.131726713 | 0.00205066 | 2.143337945 | 0.00010505 |
| 72 | 9330102E08Rik | RIKEN cDNA 9330102E08 gene | chr6 | -2.107627848 | 0.00486940 | 2.024567569 | 0.00211630 |
| 73 | Tkt | transketolase | chr14 | -2.091740646 | 0.00409522 | 2.067176939 | 0.00206869 |
| 74 | Acsl5 | acyl-CoA synthetase long-chain family member 5 | chr19 | -2.078736676 | 0.00200660 | 1.853073813 | 0.00221677 |
| 75 | Ddhd2 | DDHD domain containing 2 | chr8 | -2.068611456 | 0.00072610 | 1.865787973 | 0.00224640 |
| 76 | Slc25a1 | solute carrier family 25 member 1 | chr16 | -2.03651381 | 0.00864575 | 2.51048562 | 0.00001006 |
| 77 | Tmem120b | transmembrane protein 120B | chr5 | -2.02086396 | 0.00287219 | 1.743871312 | 0.00334209 |
| 78 | Comt | catechol-O-methyltransferase | chr16 | -1.963952441 | 0.00138519 | 2.121874011 | 0.00000058 |
| 79 | Aacs | acetoacetyl-CoA synthetase | chr5 | -1.939758149 | 0.00247771 | 1.781540914 | 0.00417571 |
| 80 | Tmem45b | transmembrane protein 45b | chr9 | -1.93866892 | 0.00754666 | 1.588854032 | 0.00752814 |
| 81 | Mecr | mitochondrial trans-2-enoyl-CoA reductase | chr4 | -1.928501137 | 0.00395059 | 1.72435937 | 0.00619958 |
| 82 | Hp | haptoglobin | chr8 | -1.886586334 | 0.00279771 | 2.248206394 | 0.00012413 |
| 83 | Hsd11b1 | hydroxysteroid 11-beta dehydrogenase 1 | chr1 | -1.881418688 | 0.00007937 | 1.60069324 | 0.00378482 |
| 84 | Hacd2 | 3-hydroxyacyl-CoA dehydratase 2 | chr16 | -1.867366825 | 0.00218642 | 1.617045546 | 0.00328187 |
| 85 | Mrap | melanocortin 2 receptor accessory protein | chr16 | -1.860112556 | 0.00563508 | 1.729569949 | 0.00198629 |
| 86 | Kcnb1 | potassium voltage-gated channel subfamily B member 1 | chr2 | -1.853407374 | 0.00063840 | 1.852136652 | 0.00426265 |
| 87 | Lss | lanosterol synthase | chr10 | -1.850836766 | 0.00013245 | 1.605997685 | 0.00385468 |
| 88 | Rmdn3 | regulator of microtubule dynamics 3 | chr2 | -1.848397365 | 0.00447196 | 1.572412075 | 0.00838924 |
| 89 | Tlcd1 | TLC domain containing 1 | chr11 | -1.840102931 | 0.00377380 | 1.710694622 | 0.00018632 |
| 90 | Cnnm2 | cyclin M2 | chr19 | -1.801393238 | 0.00000237 | 1.782632062 | 0.00004313 |
| 91 | G6pdx | glucose-6-phosphate dehydrogenase X-linked | chrX | -1.800836729 | 0.00272730 | 1.543627875 | 0.00353357 |
| 92 | Stradb | STE20-related kinase adaptor beta | chr1 | -1.798053004 | 0.00119327 | 1.599447743 | 0.00099766 |
| 93 | C1rl | complement component 1, r subcomponent-like | chr6 | -1.784992951 | 0.00033755 | 1.509456118 | 0.00999971 |
| 94 | Pcyt2 | phosphate cytidylyltransferase 2, ethanolamine | chr11 | -1.7697534 | 0.00657982 | 1.73544584 | 0.00268045 |
| 95 | Nmnat1 | nicotinamide nucleotide adenylyltransferase 1 | chr4 | -1.743290137 | 0.00188101 | 1.707795439 | 0.00008760 |
| 96 | Lrrc39 | eucine rich repeat containing 39 | chr3 | -1.740638829 | 0.00000018 | 1.386946913 | 0.00128862 |
| 97 | Phlda3 | pleckstrin homology like domain family A member 3 | chr1 | -1.716044185 | 0.00818734 | 1.743282126 | 0.00095372 |
| 98 | Mgll | monoglyceride lipase | chr6 | -1.683459704 | 0.00776891 | 1.576835916 | 0.00910642 |
| 99 | Nfil3 | nuclear factor, interleukin 3, regulated | chr13 | -1.667030556 | 0.00111362 | 1.441425339 | 0.00225324 |
| 100 | Mvd | mevalonate (diphospho) decarboxylase | chr8 | -1.65412722 | 0.00378824 | 1.673206738 | 0.00221631 |
| 101 | Cox19 | cytochrome c oxidase assembly protein 19 | chr5 | -1.650729314 | 0.00767108 | 1.774601459 | 0.00030905 |
| 102 | Slc22a3 | solute carrier family 22 member 3 | chr17 | -1.630520927 | 0.00003373 | 1.740924376 | 0.00000000 |
| 103 | Coa5 | cytochrome C oxidase assembly factor 5 | chr1 | -1.624452201 | 0.00810975 | 1.5297709 | 0.00265661 |
| 104 | Cars2 | cysteinyl-tRNA synthetase 2 (mitochondrial) | chr8 | -1.621565232 | 0.00555612 | 1.529314002 | 0.00289166 |
| 105 | Carmn | cardiac mesoderm enhancer-associated non-coding RNA | chr18 | -1.619787595 | 0.00197920 | 2.918660771 | 0.00000000 |
| 106 | Itpk1 | inositol 1,3,4-triphosphate 5/6 kinase | chr12 | -1.589336596 | 0.00256452 | 1.473029058 | 0.00358650 |
| 107 | Dpep1 | dipeptidase 1 | chr8 | -1.5757984 | 0.00097956 | 1.735499686 | 0.00043256 |
| 108 | Scd2 | stearoyl-Coenzyme A desaturase 2 | chr19 | -1.574427077 | 0.00108712 | 1.46864475 | 0.00098439 |
| 109 | Mmaa | metabolism of cobalamin associated A | chr8 | -1.569681131 | 0.00551116 | 1.454749422 | 0.00191185 |
| 110 | Ndufaf4 | NADH:ubiquinone oxidoreductase complex assembly factor 4 | chr4 | -1.559775966 | 0.00267275 | 1.462207822 | 0.00604102 |
| 111 | Mmd | monocyte to macrophage differentiation-associated | chr11 | -1.559386139 | 0.00039233 | 1.426513229 | 0.00044902 |
| 112 | Lurap1 | leucine rich adaptor protein 1 | chr4 | -1.558933711 | 0.00040246 | 1.582664412 | 0.00783060 |
| 113 | Cyb5b | cytochrome b5 type B | chr8 | -1.542238773 | 0.00036187 | 1.384604221 | 0.00049918 |
| 114 | Apol6 | apolipoprotein L 6 | chr15 | -1.532663891 | 0.00115312 | 1.570024671 | 0.00010926 |
| 115 | P2rx5 | purinergic receptor P2X 5 | chr11 | -1.490944297 | 0.00565247 | 1.557318024 | 0.00080185 |
| 116 | Pank3 | pantothenate kinase 3 | chr11 | -1.48337874 | 0.00133920 | 1.517074363 | 0.00134805 |
| 117 | Npr3 | natriuretic peptide receptor 3 | chr15 | -1.478076943 | 0.00502678 | 1.426737408 | 0.00148496 |
| 118 | Meg3 | maternally expressed 3 | chr12 | -1.448072809 | 0.00171498 | 1.483402855 | 0.00161525 |
| 119 | Ptger1 | prostaglandin E receptor 1 | chr8 | -1.443791434 | 0.00543476 | 1.446960466 | 0.00175566 |
| 120 | Srebf1 | sterol regulatory element binding transcription factor 1 | chr11 | -1.440616238 | 0.00244150 | 1.400334518 | 0.00335320 |
| 121 | Top1mt | DNA topoisomerase 1, mitochondrial | chr15 | -1.439810381 | 0.00089557 | 1.582438847 | 0.00000808 |
| 122 | Trp53inp2 | tumor protein p53 inducible nuclear protein 2 | chr2 | -1.4131865 | 0.00055115 | 1.438719591 | 0.00013957 |
| 123 | Agl | amylo-1,6-glucosidase, 4-alpha-glucanotransferase | chr3 | -1.406107676 | 0.00004941 | 1.313159241 | 0.00460060 |
| 124 | Kmt5a | lysine methyltransferase 5A | chr5 | -1.392868198 | 0.00501356 | 1.618484378 | 0.00001105 |
| 125 | H6pd | hexose-6-phosphate dehydrogenase (glucose 1-dehydrogenase) | chr4 | -1.388417752 | 0.00061754 | 1.345726239 | 0.00237135 |
| 126 | Slc26a6 | solute carrier family 26, member 6 | chr9 | -1.368193118 | 0.00959822 | 1.390056813 | 0.00881876 |
| 127 | Tmem94 | transmembrane protein 94 | chr11 | -1.346056855 | 0.00578169 | 1.410066316 | 0.00926384 |
| 128 | Zfp91 | zinc finger protein 91 | chr19 | -1.339287698 | 0.00145760 | 1.28390934 | 0.00118220 |
| 129 | Tesk1 | testis specific protein kinase 1 | chr4 | -1.324693259 | 0.00535743 | 1.542268919 | 0.00000282 |
| 130 | Dvl3 | Dishevelled segment polarity protein 3 | chr16 | -1.317613868 | 0.00035895 | 1.384705362 | 0.00071284 |
| 131 | Tubg1 | tubulin, gamma 1 | chr11 | -1.302315434 | 0.00855233 | 1.35846039 | 0.00019502 |
| 132 | Ankrd40 | ankyrin repeat domain 40 | chr11 | -1.291592189 | 0.00224489 | 1.276077607 | 0.00068951 |
| 133 | Ermp1 | endoplasmic reticulum metallopeptidase 1 | chr19 | -1.276341119 | 0.00372563 | 1.452032333 | 0.00000013 |
| 134 | Fam149b | family with sequence similarity 149, member B | chr14 | -1.276200989 | 0.00061755 | 1.236998333 | 0.00150033 |
| 135 | Usp10 | ubiquitin specific peptidase 10 | chr8 | -1.264677609 | 0.00039713 | 1.248040584 | 0.00120024 |
| 136 | Cipc | CLOCK interacting protein, circadian | chr12 | -1.255642343 | 0.00250727 | 1.186484603 | 0.00809228 |
| 137 | Tigar | Trp53 induced glycolysis regulatory phosphatase | chr6 | -1.246052953 | 0.00380992 | 1.328222343 | 0.00024874 |
| 138 | Med24 | mediator complex subunit 24 | chr11 | -1.245250438 | 0.00001704 | 1.213322503 | 0.00006806 |
| 139 | Mapk8ip3 | mitogen-activated protein kinase 8 interacting protein 3 | chr17 | -1.237510664 | 0.00746874 | 1.325517425 | 0.00028239 |
| 140 | Srr | serine racemase | chr11 | -1.237403941 | 0.00932449 | 1.248915057 | 0.00383917 |

**Supplemental Table 2 (continued). A complete list of 150 genes whose mRNAs are downregulated by orchiectomy (orx *vs.* sham ctrl) but upregulated by exogenous dihydrotestosterone (DHT) administration to orchiectomized mice (Orx+DHT *vs.* orx) in the aorta in mice administered Aldo-salt**

| # | Gene ID | Description | Location | Fold Change (orx vs. ctrl) | p-value | Fold Change (orx+DHT vs. orx) | p-value |
|---|---|---|---|---|---|---|---|
| 141 | Neurl4 | neuralized E3 ubiquitin protein ligase 4 | chr11 | -1.227717106 | 0.00109247 | 1.298341332 | 0.00120645 |
| 142 | Rnf10 | ring finger protein 10 [ | chr5 | -1.2086912 | 0.00703013 | 1.325846168 | 0.00061523 |
| 143 | Scaf1 | SR-related CTD-associated factor | chr7 | -1.207545164 | 0.00331215 | 1.290980285 | 0.00004449 |
| 144 | Arfip2 | ADP-ribosylation factor interacting protein 2 | chr7 | -1.202275336 | 0.00061851 | 1.254769016 | 0.00001189 |
| 145 | Dvl1 | dishevelled segment polarity protein | chr4 | -1.190127928 | 0.00946186 | 1.271239888 | 0.00058910 |
| 146 | Tab2 | TGF-beta activated kinase 1/MAP3K7 binding protein 2 | chr10 | -1.189513717 | 0.00049606 | 1.152624365 | 0.00630599 |
| 147 | Smg5 | SMG5 nonsense mediated mRNA decay factor | chr3 | -1.181871224 | 0.00798154 | 1.191993936 | 0.00409333 |
| 148 | Mbd6 | methyl-CpG binding domain protein 6 | chr10 | -1.175619455 | 0.00707640 | 1.360700607 | 0.00029690 |
| 149 | Crebzf | CREB/ATF bZIP transcription factor | chr7 | -1.173629123 | 0.00559876 | 1.185244594 | 0.00155486 |
| 150 | R3hdm2 | R3H domain containing 2 | chr10 | -1.140454302 | 0.00334356 | 1.222527753 | 0.00160605 |

**Supplemental Table 3. A complete list of 65 signaling pathways upregulated by orchiectomy (orx *vs.* sham ctrl) but downregulated by exogenous dihydrotestosterone (DHT) administration to orchiectomized mice (Orx+DHT *vs.* orx) in the aorta in mice administered Aldo-salt**

| # | Pathway | P-value | Genes |
|---|---|---|---|
| 1 | Interleukin-2 signaling pathway | 1.37E-16 | Itk;Pcna;Txk;Tcf7;Cxcr5;Ptpn22;Ikzf3;Rasgrp1;Spn;Ctsk;Myb;Itgax;Gpc3;Il21r;Stat4;Pmaip1;Itgb7;Cytip;Ccr2;Cd52;Cr2;Sh2d1a;Cd2;Ptprc;Sell;Lck;Cd5;Il2rb;Pdcd4;Cd27;Irf5;Ptpn6;Ltb;Cd247;Cd69;Il7r;Lat;Il18r1 |
| 2 | T helper cell surface molecules | 2.19E-16 | Cd2;Ptprc;Cd8a;Cd28;Cd3g;Cd247;Cd3e;Itgal;Cd3d |
| 3 | Adaptive immune system | 5.13E-16 | Blk;Itk;Cd3g;Kif11;Itgal;Cd3e;Rasgrp1;Cd3d;Cd79b;Cd79a;H2-DMa;Grap2;Cd19;Ctsk;Btla;Blnk;Itgb7;H2-Oa;H2-Ob;H2-Ab1;Cd74;H2-Eb2;Ptprc;Sell;Cd8a;Lck;Cd28;Ptpn6;Cd247;Card11;Lat;H2-Aa |
| 4 | T cell receptor signaling in naive CD4+ T cells | 8.86E-16 | Map4k1;Itk;Cd3g;Cd3e;Cd3d;Rasgrp1;Ptprc;Lck;Grap2;Cd28;Ptpn6;Cd247;Card11;Lat |
| 5 | T cell receptor signaling in naive CD8+ T cells | 1.71E-15 | Cd3g;Cd3e;Cd3d;Rasgrp1;Ptprc;Cd8a;Lck;Grap2;Cd28;Ptpn6;Cd247;Card11;Lat |
| 6 | Cell adhesion molecules (CAMs) | 3.86E-15 | H2-Eb2;Selplg;Itgal;Spn;Cd2;H2-DMa;Alcam;Ptprc;Sell;Cd8a;Cd28;Itgb7;H2-Oa;H2-Ob;H2-Ab1;Cd22;H2-Aa |
| 7 | Immune system | 4.89E-15 | Blk;Itk;Cd3g;Hmgb1;Kif11;Itgal;Cd3e;Rasgrp1;Cd3d;Cd79b;Cd79a;H2-DMa;Grap2;Cd19;Ctsk;Btla;Blnk;Itgb7;H2-Oa;H2-Ob;H2-Ab1;Ccr2;Cd74;H2-Eb2;Nup210;Ptprc;Sell;Cd8a;Lck;Il2rb;Cd28;Irf8;Irf5;Ptpn6;Cd247;Il7r;Card11;Lat;H2-Aa |
| 8 | Generation of second messenger molecules | 1.14E-14 | Itk;H2-Eb2;Lck;Grap2;Cd3g;Cd247;Cd3e;Cd3d;H2-Ab1;Lat;H2-Aa |
| 9 | PD-1 signaling | 1.02E-13 | H2-Eb2;Ptprc;Lck;Cd3g;Ptpn6;Cd247;Cd3e;Cd3d;H2-Ab1;H2-Aa |
| 10 | Primary immunodeficiency | 4.10E-13 | Cd79a;Ptprc;Cd8a;Lck;Cd19;Blnk;Tnfrsf13c;Cd3e;Il7r;Cd3d |
| 11 | Hematopoietic cell lineage | 1.10E-12 | Cr2;H2-Eb2;Cd3g;Cd3e;Cd3d;Cd2;Cd8a;Cd5;Cd19;Cd37;Il7r;Ms4a1;Cd22 |
| 12 | CostimuLation by the Cd28 family | 1.87E-12 | H2-Eb2;Lck;Grap2;Btla;Cd28;Cd3g;Ptpn6;Cd247;Cd3e;Cd3d;H2-Ab1;H2-Aa |
| 13 | Leptin influence on immune response | 2.04E-11 | Mmp2;Cd3e;Cd3d;Cd79b;Cd79a;Azgp1;Alcam;Sell;Itgax;Irf8;Ltb;Il7r;Ccr2 |
| 14 | T cell receptor signaling pathway | 3.03E-11 | Itk;Cd3g;Cd3e;Rasgrp1;Cd3d;Ptprc;Cd8a;Lck;Grap2;Cd28;Ptpn6;Cd247;Card11;Lat |
| 15 | T cell receptor reguLation of apoptosis | 2.00E-10 | Top2a;Lef1;Cd3g;Hmgb1;Itgal;Cd3e;Rasgrp1;Spn;Myb;Rac2;Il21r;Pmaip1;Ccr2;Map4k1;Cr2;Sema4d;Mmp2;Rhoh;Cd2;Ptprc;Lck;Cd28;Ltb;Il7r;H2-Aa |
| 16 | T cell signal transduction | 2.12E-10 | Lat2;Itk;Ptprc;Lck;Grap2;Cd28;Cd3g;Cd3e;Cd3d;Rasgrp1;Lat |
| 17 | Intestinal immune network for IgA production | 3.93E-10 | H2-DMa;H2-Eb2;Cd28;Itgb7;Tnfrsf13c;H2-Oa;H2-Ob;H2-Ab1;H2-Aa |
| 18 | T cell activation co-stimuLatory signal | 4.45E-10 | Itk;Lck;Cd28;Cd3g;Cd247;Cd3e;Cd3d |
| 19 | Lck and Fyn tyrosine kinases in initiation of T cell receptor activation | 7.96E-10 | Ptprc;Lck;Cd3g;Cd247;Cd3e;Cd3d |
| 20 | Interleukin-17 signaling pathway | 2.29E-09 | Cd2;Cd8a;Cd3g;Cd247;Cd3e;Cd3d |
| 21 | NO2-dependent IL-12 pathway in NK cells | 5.57E-09 | Cd2;Stat4;Cd3g;Cd247;Cd3e;Cd3d |
| 22 | Interleukin-12-mediated signaling events | 6.57E-09 | Cd8a;Lck;Il2rb;Stat4;Cd3g;Cd247;Cd3e;Cd3d;Il18r1 |
| 23 | HIV-induced T cell apoptosis | 6.83E-09 | Cd28;Cd3g;Cd247;Cd3e;Cd3d |
| 24 | Asthma | 7.27E-09 | H2-DMa;H2-Eb2;Prg2;H2-Oa;H2-Ob;H2-Ab1;H2-Aa |
| 25 | Antigen-activated B-cell receptor generation of second messengers | 7.55E-09 | Blk;Map4k1;Cr2;Cd79b;Lat2;Cd79a;Ptprc;Cd19;Cd28;Rac2;Blnk;Ptpn6;Card11;Cd22 |
| 26 | Inhibition of T cell receptor signaling by activated Csk | 1.20E-08 | Ptprc;Lck;Cd3g;Cd247;Cd3e;Cd3d |
| 27 | Viral myocarditis | 1.46E-08 | H2-DMa;H2-Eb2;Rac2;Cd28;Itgal;H2-Oa;H2-Ob;H2-Ab1;H2-Aa |
| 28 | MEF2D role in T cell apoptosis | 2.31E-08 | Ptprc;Lck;Cd3g;Cd247;Cd3e;Cd3d;Lat |
| 29 | Allograft rejection | 3.49E-08 | H2-DMa;H2-Eb2;Cd28;H2-Oa;H2-Ob;H2-Ab1;H2-Aa |
| 30 | CTL mediated immune response against target cells | 6.78E-08 | Cd3g;Cd247;Cd3e;Itgal;Cd3d |
| 31 | Graft-versus-host disease | 7.39E-08 | H2-DMa;H2-Eb2;Cd28;H2-Oa;H2-Ob;H2-Ab1;H2-Aa |
| 32 | Type 1 diabetes mellitus | 1.04E-07 | H2-DMa;H2-Eb2;Cd28;H2-Oa;H2-Ob;H2-Ab1;H2-Aa |
| 33 | Interleukin-12/Stat4 pathway | 1.45E-07 | Stat4;Cd28;Cd3g;Cd247;Cd3e;Cd3d;Il18r1 |
| 34 | Cell surface interactions at the vascular wall | 1.75E-07 | Spn;Cd2;Selplg;Sell;Lck;Itgax;Ptpn6;Itgal;Slc7a10 |
| 35 | MHC class II antigen presentation | 3.85E-07 | Cd74;H2-Eb2;H2-DMa;Ctsk;Kif11;H2-Oa;H2-Ob;H2-Ab1;H2-Aa |
| 36 | Autoimmune thyroid disease | 4.05E-07 | H2-DMa;H2-Eb2;Cd28;H2-Oa;H2-Ob;H2-Ab1;H2-Aa |
| 37 | Tob role in T-cell activation | 5.86E-07 | Cd28;Cd3g;Cd247;Cd3e;Cd3d |
| 38 | Antigen processing and presentation | 6.77E-07 | Cd74;H2-DMa;H2-Eb2;Cd8a;H2-Oa;H2-Ob;H2-Ab1;H2-Aa |
| 39 | Hemostasis pathway | 1.20E-06 | Selplg;Prcp;Kif11;Slc7a10;Itgal;Rasgrp1;Spn;Cd2;Sell;Kifc1;Lck;Myb;Alb;Rac2;Itgax;Ptpn6;Lat |
| 40 | ImmunoreguLatory interactions between a lymphoid and a non-lymphoid cell | 1.31E-06 | Sell;Cd8a;Cd19;Cd3g;Itgb7;Cd247;Cd3e;Itgal;Cd3d |
| 41 | CD8/T cell receptor downstream pathway | 2.60E-06 | Cd8a;Il2rb;Stat4;Cd3g;Cd247;Cd3e;Cd3d |
| 42 | Leishmaniasis | 3.83E-06 | H2-DMa;H2-Eb2;Ptpn6;H2-Oa;H2-Ob;H2-Ab1;H2-Aa |
| 43 | Interleukin-4 reguLation of apoptosis | 5.71E-06 | Top2a;Arl4c;Pou2af1;Tcf7;Mcm3;Cd27;Mcm4;Ltb;Il7r;Rasgrp1;Chst2;H2-Aa |
| 44 | Stathmin and breast cancer resistance to antimicrotubule agents | 7.85E-06 | Cd2;Cd3g;Cd247;Cd3e;Cd3d |
| 45 | CXCR4 signaling pathway | 1.02E-05 | Blk;Ptprc;Lck;Cd3g;Ptpn6;Cd247;Cd3e;Cd3d |
| 46 | Alpha-M beta-2 integrin signaling | 3.22E-05 | Blk;Selplg;Lck;Mmp2;Hmgb1 |
| 47 | Immune system signaling by interferons, interleukins, prolactin, and growth hormones | 4.86E-05 | H2-Eb2;Nup210;Lck;Il2rb;Blnk;Irf8;Irf5;Ptpn6;Il7r;H2-Ab1;H2-Aa |
| 48 | Ras-independent pathway in NK cell-mediated cytotoxicity | 5.86E-05 | Cd2;Cd28;Ptpn6;Lat |
| 49 | Apoptotic DNA fragmentation and tissue homeostasis | 8.21E-05 | Top2a;Hmgb2;Hmgb1 |
| 50 | Leukocyte transendothelial migration | 9.18E-05 | Itk;Txk;Mmp2;Rac2;Rhoh;Ezr;Itgal |

**Supplemental Table 3 (continued). A complete list of 65 signaling pathways upregulated by orchiectomy (orx *vs.* sham ctrl) but downregulated by exogenous dihydrotestosterone (DHT) administration to orchiectomized mice (Orx+DHT *vs.* orx) in the aorta in mice administered Aldo-salt**

| # | Pathway | P-value | Genes |
|---|---|---|---|
| 51 | B lymphocyte cell surface molecules | 1.12E-04 | Cr2;Ptprc;Itgal |
| 52 | Cytokine-cytokine receptor interaction | 1.49E-04 | Il2rb;Cxcr5;Il21r;Cd27;Tnfrsf13c;Ltb;Cxcr6;Il7r;Il18r1;Ccr2 |
| 53 | Natural killer cell receptor signaling pathway | 2.04E-04 | Lat2;Lck;Rac2;Ptpn6;Lat |
| 54 | T cell receptor/JNK pathway | 2.43E-04 | Map4k1;Grap2;Lat |
| 55 | Natural killer cell-mediated cytotoxicity | 2.45E-04 | Lck;Sh2d1a;Rac2;Ptpn6;Cd247;Itgal;Lat |
| 56 | Interferon-gamma signaling pathway | 2.45E-04 | H2-Eb2;Irf8;Irf5;Ptpn6;H2-Ab1;H2-Aa |
| 57 | Systemic lupus erythematosus | 2.68E-04 | H2-Eb2;H2-DMa;Cd28;H2-Oa;H2-Ob;H2-Ab1;H2-Aa |
| 58 | Adhesion and diapedesis of granulocytes | 3.01E-04 | Selplg;Sell;Itgal |
| 59 | Signaling by the B cell receptor (BCR) | 4.42E-04 | Blk;Cd79b;Cd79a;Cd19;Blnk;Rasgrp1;Card11 |
| 60 | Dendritic cells in reguLating TH1 and TH2 development | 4.44E-04 | Cd2;Cd5;Itgax |
| 61 | Phagosome | 4.98E-04 | H2-Eb2;H2-DMa;H2-Oa;H2-Ob;Coro1a;H2-Ab1;H2-Aa |
| 62 | Interferon signaling | 8.33E-04 | H2-Eb2;Nup210;Irf8;Irf5;Ptpn6;H2-Ab1;H2-Aa |
| 63 | Interleukin-2 receptor beta chain in T cell activation | 1.23E-03 | Pcna;Il2rb;Ptpn6;Ikzf3 |
| 64 | Endogenous Toll-like receptor signaling | 1.26E-03 | Hmgb1;S100a9;S100a8 |
| 65 | Focal adhesion | 1.26E-03 | Blk;Parvg;Txk;Col11a1;Rac2;Itgax;Itgb7;Itgal |

**Supplemental Table 4. A complete list of 19 signaling pathways downregulated by orchiectomy (orx *vs.* sham ctrl) but upregulated by exogenous dihydrotestosterone (DHT) administration to orchiectomized mice (Orx+DHT *vs.* orx) in the aorta in mice administered Aldo-salt**

| # | Pathway | P-value | Genes |
|---|---|---|---|
| 1 | Triglyceride biosynthesis | 1.64E-13 | Slc25a1;Acly;Fasn;Elovl3;Gpd1;Acsl5;Elovl6;Lpin1;Agpat2;Acaca |
| 2 | Fatty acid, triacylglycerol, and ketone body metabolism | 3.63E-12 | Slc25a1;Srebf1;Elovl3;Acsl5;Elovl6;Acacb;Agpat2;Acaca;Acly;Med24;Fasn;Gpd1;Me1;Plin2;Lpin1 |
| 3 | Fatty acyl-CoA biosynthesis | 3.44E-11 | Slc25a1;Acly;Fasn;Elovl3;Acsl5;Elovl6;Acaca |
| 4 | Lipid and lipoprotein metabolism | 1.13E-10 | Slc25a1;Srebf1;Pcyt2;Elovl3;Acsl5;Elovl6;Acacb;Lss;Agpat2;Acaca;Hsd11b1;Acly;Med24;Fasn;Gpd1;Me1;Pnpla3;Plin2;Mvd;Lpin1;Mgll |
| 5 | Fatty acid biosynthesis | 4.98E-10 | Acly;Acss2;Mecr;Fasn;Acsl5;Acacb;Acaca |
| 6 | ChREBP activates metabolic gene expression | 1.22E-09 | Acly;Mlxipl;Fasn;Acacb;Acaca |
| 7 | Metabolism | 2.72E-08 | Pank3;H6pd;Acss2;Ncan;Comt;Acacb;Agpat2;Acaca;Hk2;Hsd11b1;Impa2;Me1;Mecr;Pcyt2;Kcnb1;Agl;Elovl3;Itpk1;Acsl5;Elovl6;Pgd;Lss;Acly;Mlxipl;Pfkl;Med24;Nmnat1;Fasn;Gpd1;Pnpla3;Mvd;Tkt;Lpin1;Mgll |
| 8 | Striated muscle contraction | 9.12E-06 | Mybpc1;Mybpc2;Actn3;Tnnc2;Neb |
| 9 | Shuttle for transfer of acetyl groups from mitochondria to the cytosol | 2.25E-05 | Slc25a1;Acly;Me1 |
| 10 | Muscle contraction | 3.24E-05 | Mybpc1;Mybpc2;Actn3;Tnnc2;Neb |
| 11 | Ghrelin pathway | 3.40E-05 | Srebf1;Fasn;Plin2;Acaca |
| 12 | Pyruvate metabolism | 4.34E-05 | Acss2;Slc16a1;Me1;Acacb;Acaca |
| 13 | Integration of energy metabolism | 4.42E-05 | Acly;Mlxipl;Kcnb1;Fasn;Tkt;Acacb;Acaca |
| 14 | Glycerophospholipid biosynthesis | 4.62E-05 | Pcyt2;Gpd1;Pnpla3;Lpin1;Agpat2;Mgll |
| 15 | Pentose phosphate pathway | 4.66E-05 | Pfkl;H6pd;Pgd;Tkt |
| 16 | AMPK signaling | 1.58E-04 | Srebf1;Stradb;Fasn;Acacb;Acaca |
| 17 | Carbohydrate metabolism | 4.05E-04 | Slc25a1;Pfkl;Ncan;Agl;Pgd;Slc2a5;Tkt;Hk2 |
| 18 | Adipogenesis | 5.05E-04 | Srebf1;Dvl1;Pnpla3;Plin2;Lpin1;Agpat2 |
| 19 | Acyl chain remodeling of diacylglycerol and triacylglycerol | 5.51E-04 | Pnpla3;Mgll |

**Supplemental Table 5. A complete list of antibodies used in the current study**

| Antibody | Application | Source/ Reactivity | Manufacture | Catalog | Dilution/amount | Fluorophore | Reference |
|---|---|---|---|---|---|---|---|
| AR | IHC | Rabbit anti-mouse | Abcam | Ab74272 | 1:300 | | Dufour et al., Cell Reports. 2022;38, 110534 |
| AR | WB | Rabbit anti-mouse | Cell Signaling | 5153 | 1:2000 | | Li et al. Maturitas.2009; 63, 142-148. |
| IL-6 | IHC | Rabbit anti-mouse | Bioss | Bs-0782R | 1:2000 | | Sun et al., Biomed Res Int. 2015;919401 |
| MR | IHC | Mouse anti-mouse | DSHB | rMR1-18 1D5 | 1.1 mg/ml | | Prager et al., PLoS One. 2010;5(12):e14344 |
| p-STAT3 | IHC | Rabbit anti-mouse | Cell Signaling | 9145 | 1:400 | | Cheng et al., Nat Commun. 2022;29;13(1):4418 |
| F4/80 | IHC | Rabbit anti-mouse | Cell Signaling | 70076 | 1:1000 | | Wang et al., Front Immunol. 2022;13:901209 |
| Ly6G | IHC | Rat anti-mouse | BD Pharmingen | 551459 | 1:1500 | | Fleming, et. al. J Immunol. 1993; 151(5):2399-2408 |
| PD-1 | WB, IHC | Rabbit anti-mouse | Cell Signaling | 84651 | WB 1:1000<br>IHC 1:250 | | Yuan et al., JCI insight. 2022;7(11):e157788. |
| PD-1 | IHC | Rabbit anti-human | Cell Signaling | 86163 | IHC 1:250 | | Ju et al., Nat Commun. 2022; 13(1):5378 |
| CD3ε | WB, IHC | Rabbit anti-mouse | Cell Signaling | 99940 | WB 1:1000<br>IHC 1:500 | | Wallace-Povirk et al., Sci Rep. 2022;5;12(1):11346 |
| CD19 | WB, IHC | Rabbit anti-mouse | Cell Signaling | 90176 | WB 1:1000<br>IHC 1:2000 | | Tedder et al. Immunity. 1997;6, 107-18. |
| GAPDH | WB | Rabbit anti-mouse | Cell Signaling | 2118 | 1:5000 | | Wong et al., J Clin Invest. 2022;132(15):e152635 |
| AR | ChIP | Mouse anti-mouse | Santa Cruz | sc7305 | 5 μg/ChIP | | Sharma et al., Front Oncol. 2022;12: 824594 |
| AR | ChIP | Rabbit anti-mouse | Millipore | 17-10489 | 1 μg/ChIP | | Nigro et al., Diabetes. 2021;70(6):1250–1264 |
| CD45 | FCM | Rat anti-mouse | Biolegend | 103128 | 0.25 μg/$10^6$ cells | Alexa Fluor 700 | Tian et al., Nat Commun. 2016;7:13283. |
| CD45 | FCM | Rat anti-mouse | Biolegend | 103126 | 0.25 μg/$10^6$ cells | Pacific blue | Pattabiraman et al., Science. 2016;351(6277):aad3680 |
| CD3ε | FCM | Rat anti-mouse | Biolegend | 100320 | 0.5 μg/$10^6$ cells | PE/Cy7 | Cabañero et al. Elife. 2020;9:e55582 |
| CD3 | FCM | Rat anti-mouse | Biolegend | 100240 | 0.25 μg/$10^6$ cells | Alexa Fluor 594 | Lederer et al., Immunity. 2020;53(6):1281-1295.e5. |
| CD19 | FCM | Rat anti-mouse | Biolegend | 115534 | 0.25 μg/$10^6$ cells | PerCP | Faust et al., J Clin Invest. 2020;130(10):5493-5507 |
| PD-1 | FCM | Rat anti-mouse | Biolegend | 135210 | 0.25 μg/$10^6$ cells | APC | Mandal et al. Cell Reports. 2021;35(6):109094. |
| PD-1 | FCM | Rat anti-mouse | Biolegend | 135219 | 0.125 μg/$10^6$ cells | BV605 | Mogilenko et al., Immunity 2021;54(1):99-115.e12 |
| Ly6G | FCM | Rat anti-mouse | Biolegend | 127608 | 0.25 μg/$10^6$ cells | PE | Okada et al., J Biol Chem. 2014;289(47):32926-36 |
| F4/80 | FCM | Rat anti-mouse | Biolegend | 123120 | 1 μg/$10^6$ cells | Alexa Fluor 488 | Wheeler et al., Nat Commun. 2015;6: 8964. |
| F4/80 | FCM | Rat anti-mouse | Biolegend | 123110 | 1 μg/$10^6$ cells | PE | Xiang et al., PNAS 2014;111(48):E5159-68 |

*Note:* IHC, immunocytochemistry. WB, western blot. ChIP, Chromatin immunoprecipitation. FCM, flow cytometry.

#### Supplemental Table 6. A complete list of mouse PCR primers used in the current study

| Gene | Primer | Sequence | Application |
|---|---|---|---|
| Ar | Forward | 5'-GGACCATGTTTTACCCATCG-3' | Real-time PCR |
| | Reverse | 5'-TCGTTTCTGCTGGCACATAG-3' | |
| Nr3c2 | Forward | 5'-ATGGGTACCCGGTCCTAGAG-3' | Real-time PCR |
| | Reverse | 5'-ACCAAGCAGATCTTGGAAGG-3' | |
| Sgk1 | Forward | 5'-TCAGAGCGGAATGTTCTGTTG-3' | Real-time PCR |
| | Reverse | 5'-AGCGGTCTGGAATGAGAAGTG-3' | |
| Scnn1a | Forward | 5'-TACTTCAGCTACCCCGTGAGT-3' | Real-time PCR |
| | Reverse | 5'-AAAAAGCGTCTGTTCCGTGAT-3' | |
| Scnn1b | Forward | 5'-ACCCGGTGGTTCTCAATTTGT-3' | Real-time PCR |
| | Reverse | 5'-AAGTTCCGCAAGGTACACACA-3' | |
| Scnn1g | Forward | 5'-GGCACCGACCATTAAGGACC-3' | Real-time PCR |
| | Reverse | 5'-CTGTCAGCGTGAACGCAATC-3' | |
| Arntl | Forward | 5'-ATCAGCGACTTCATGTCTCC-3' | Real-time PCR |
| | Reverse | 5'-CTCCCTTGCATTCTTGATCC-3' | |
| Tgfb2 | Forward | 5'-CCATCCCGCCCACTTTCTAC-3' | Real-time PCR |
| | Reverse | 5'-TCTGGTTTTCACAACCTTGCT-3' | |
| Mmp2 | Forward | 5'-GGACAAGTGGTCCGCGTAAA-3' | Real-time PCR |
| | Reverse | 5'-CCGACCGTTGAACAGGAAGG-3' | |
| Il1b | Forward | 5'-TCGCTC AGGGTCACAAGAAA-3' | Real-time PCR |
| | Reverse | 5'-CATCAGAGGCAAGGAGGAAAAC-3' | |
| Il6 | Forward | 5'-ACAAGTCGGAGGCTTAATTACACAT-3' | Real-time PCR |
| | Reverse | 5'-TTGCCATTGCACAACTCTTTTC-3' | |
| Il6ra | Forward | 5'-AGCGACACTGGGGACTATTTA-3' | Real-time PCR |
| | Reverse | 5'-ACAGCCTTCGTGGTTGGAG-3' | |
| Il6st | Forward | 5'-TGGAGTGAGGAGGCTAGTGG-3' | Real-time PCR |
| | Reverse | 5'-ATTTTCCCATTGGCTTCAGA-3' | |
| Ccl2 | Forward | 5'-CTTCCTCCACCACCATGCA-3' | Real-time PCR |
| | Reverse | 5'-CCAGCCGGCAACTGTGA-3' | |
| Ccl4 | Forward | 5'-TTCCTGCTGTTTCTCTTACACCT-3' | Real-time PCR |
| | Reverse | 5'-CTGTCTGCCTCTTTTGGTCAG-3' | |
| Tnf | Forward | 5'-CCCTCACACTCAGATCATCTTCT-3' | Real-time PCR |
| | Reverse | 5'-GCTACGACGTGGGCTACAG-3' | |
| 36B4 | Forward | 5'-CCCTGAAGTGCTCGACATCA-3' | Real-time PCR |
| | Reverse | 5'-TGCGGACACCCTCCAGAA-3' | |
| Pdcd1 | Forward | 5'-ACCCTGGTCATTCACTTGGG-3' | Real-time PCR |
| | Reverse | 5'- CATTTGCTCCCTCTGACACTG-3' | |
| Pdcd1 | Forward | 5'-TCATTCCACTCACAAGTCAATCAA-3' | ChIP Exon 6 |
| | Reverse | 5'-TCTTCCCTTCTCATCTCATTGTGA-3' | |
| Pdcd1 | Forward | 5'-CCTTTAGCTTCTGGGAAATGTTT-3' | ChIP Exon 4 |
| | Reverse | 5'-CCCTGGAGGTAATGGCAAGTTTCC-3' | |
